# Training Strategy Optimization to Mitigate Shortcut Learning in Pan-Cancer Drug Response Prediction

**DOI:** 10.64898/2026.05.23.725295

**Authors:** Kazuki Shimamoto, Takafumi Ito, Artem Lysenko, Tatsuhiko Tsunoda

## Abstract

**Background:** Prediction of *in vivo* drug response is a central challenge in precision medicine, but the scarcity of labeled clinical data still necessitates the use of large-scale cancer cell line resources for model training. Domain adaptation methods, which aim to transfer knowledge learned from a source domain (cell lines) to a target domain (patients) by aligning feature distributions across domains, are a promising approach to bridge the gap between *in vitro* models and *in vivo* patients. However, we observed that these methods can exhibit a significant discrepancy between pan-cancer evaluation metrics and cancer type-specific prediction accuracy. This performance gap warrants a detailed investigation into their underlying predictive characteristics.

**Results:** We discovered that cancer-type-specific class imbalances in training data can lead domain adaptation models to engage in shortcut learning, where they primarily discriminate between cancer types rather than capturing the actual biological determinants of drug sensitivity. To address this, we propose a strategy of combining two approaches: (1) excluding cancer types causing imbalance from the training data, and (2) adjusting class balance through oversampling and class weighting while retaining cancer types causing the imbalance. Among all configurations tested in conjunction with the CODE-AE (Context-aware Deconfounding AutoEncoder) framework, the combination of moderate oversampling (30% non-responder ratio) with class weighting achieved the best performance, significantly improving prediction accuracy in 5 out of 11 external patient cohorts from TCGA and GEO.

**Conclusions:** Our findings demonstrate that appropriate class imbalance correction—rather than wholesale exclusion of imbalanced cancer subtypes—enables effective utilization of biologically relevant information shared across cancer types for drug response prediction. This study highlights the critical importance of jointly optimizing training data composition and class balance adjustment strategies in developing robust pan-cancer drug response prediction models for precision medicine applications.

**Highlights:**

- Identified a critical discrepancy in current domain adaptation models for drug response prediction: high pan-cancer accuracy often masks poor performance within specific cancer types.
- Revealed the root cause as “shortcut learning,” where models tend to distinguish between cancer tissue types (hematological vs. solid) rather than learning individual drug sensitivity.
- Discovered severe class imbalance in training data, with hematological cell lines being disproportionately drug-responsive across multiple chemotherapeutics.
- Proposed an architecture-agnostic fix using the CODE-AE framework: moderate oversampling (30% minority ratio) combined with class weighting.
- Demonstrated significant improvements in 5 of 11 external patient cohorts, showing that correcting class bias is more effective than simply excluding problematic data.

## 1. Introduction

The increasing complexity of modern biomedical research necessitates a cross-disciplinary approach, often resulting in divergent perspectives and disparate analytical methodologies [1]. In the case of data science and AI, this can increasingly often lead to misalignment of objectives, where predictive models achieve high statistical accuracy but lack biological relevance [2]. One example of this is often observed in AI-driven pan-cancer studies, where broad patterns are often prioritized over specific disease pathologies. For example, if such models fail to sufficiently control for differences between specific cancers, these can completely dominate the predicted response and mask critical biological differences that actually determine the treatment optimality for specific patients.

In this study we demonstrate how particular structural features of pan-cancer datasets lead to emergence of this problem and explore various strategies to address it. The proposed weighted training strategy was effective at allowing the model to correctly resolve differences in drug response within individual cancer types, which were previously lost. Importantly, this was achieved without the need to substantially overhaul the design of the original neural network itself.

### 1.1. Precision Medicine and the Utilization of Cell Lines

A key challenge in precision medicine is creation of high-quality drug response prediction models that can guide treatment strategies and lead to optimal outcomes for individual patients. Drug efficacy is usually determined by complex interactions among multiple genes [3], and advanced machine learning methods, such as deep neural networks, are highly effective in capturing such multidimensional relationships from high-throughput gene expression data [4, 5]. However, collecting sufficient clinical and, in particular, *in vivo* data for training these models remains difficult. Consequently, cancer cell line data from resources such as the Cancer Cell Line Encyclopedia (CCLE) [6] and Genomics of Drug Sensitivity in Cancer (GDSC) [7], which are readily accessible and amenable to large-scale screening, have been widely used as surrogates for clinical samples.

### 1.2. Domain Adaptation from *In Vitro* to *In Vivo*

When models trained on cell lines are applied to actual patients, differences between culture and *in vivo* environments pose a major barrier, resulting in a significant degradation in prediction accuracy—a challenge known as the out-of-distribution problem [8], which can be addressed by using domain adaptation methods [9–11]. One of such approaches is CODE-AE (Context-aware Deconfounding AutoEncoder) [11] that leverages special neural network design patterns to successfully extract biological signals shared between cell line and patient data while removing confounding factors. This work has demonstrated that a model trained solely on cell line drug response data can achieve high prediction accuracy on patient data.

### 1.3. Problem Statement: Discrepancy Between Hematological and Solid Cancers

CODE-AE aims to improve generalization performance by treating all cancer types collectively. However, in our replication study we have observed the following two issues. First, although the model showed reasonable prediction performance under pan-cancer evaluation, accuracy declined substantially when assessed within individual cancer types. Second, prediction score distributions differed systematically between hematological and solid cancers, suggesting that the model may be distinguishing between cancer types rather than learning drug sensitivity. Detailed results are presented in Section 3.1.

A closer examination of the training data revealed that for many drugs, hematological cancer cell lines contained a markedly higher proportion of drug-responsive samples compared to solid cancer cell lines. While hematological and solid cancers share common drug mechanisms of action at the cellular level, including DNA damage response and apoptosis pathways [12, 13], they differ substantially in gene expression profiles and tumor microenvironment [14–16]. When such biological differences combine with class imbalance in the training data, the model risks falling into shortcut learning [17], whereby it learns simple spurious correlations rather than complex features that truly underlie drug sensitivity. In simple terms, the model may learn the spurious rule “if hematological cancer, then responsive; if solid cancer, then non-responsive,” and consequently fail to capture patient-level differences in drug sensitivity within each cancer type.

### 1.4. Objectives and Approach

The objective of this study is to address the problems described above and construct a more accurate and robust drug response prediction model. We do not modify the CODE-AE model architecture; instead, we aim to improve performance solely by altering the training data composition and applying class weighting during training. To address the class imbalance in hematological cancers, we propose and comparatively evaluate two approaches:

The first approach excludes hematological cancers from the training data entirely, fitting the model on solid cancers alone. This is based on a hypothesis that large biological differences between hematological and solid cancers are primary drivers of shortcut learning, and that removing hematological cancers would encourage the model to learn drug response features common to solid cancers rather than to identify cancer types.

The second approach retains hematological cancers but adjusts for class imbalance. This strategy assumes that these cancers possess shared drug response features with solid tumors that can be leveraged if the data bias is corrected. Specifically, we apply oversampling and class weighting to the responder and non-responder groups. This would encourage the model to focus on universal biological markers of sensitivity rather than simply identifying cancer types.

By comparing the prediction accuracy and robustness of these two approaches, we aim to identify the optimal training data composition strategy for pan-cancer drug response prediction.

## 2. Materials and Methods

### 2.1. Target Drugs

Following the original CODE-AE paper [11], we selected five chemotherapeutic agents: 5-Fluorouracil (antimetabolite), Cisplatin (platinum-based agent), Gemcitabine (antimetabolite), Sorafenib (multi-kinase inhibitor), and Temozolomide (alkylating agent). For these drugs, cell line drug response data were obtained from CCLE (Section 2.2.1) and GDSC (Section 2.2.2), and clinical drug response data were obtained from TCGA (Section 2.2.3) and GEO (Section 2.2.4).

### 2.2. Datasets

#### 2.2.1. CCLE

CCLE is a project led by the Broad Institute and Novartis that provides detailed genetic information and pharmacological profiles for more than 1,000 human cancer cell lines [6]. We used CCLE gene expression data as input features for each cell line during model training. Gene expression values were log₂(TPM + 1) transformed.

#### 2.2.2. GDSC

GDSC is the world’s largest database of cancer cell line drug sensitivity, operated by the Wellcome Sanger Institute, and contains response data for hundreds of compounds [7]. We used the half-maximal inhibitory concentration (IC50) for each cell line as the ground-truth label for drug response during training.

Drug response was binarized following the methodology of the original CODE-AE paper [11]: for each drug, the median IC50 value across all samples was used as the threshold. Samples with IC50 values below the median (high drug sensitivity) were defined as responders, and those at or above the median (low drug sensitivity) were defined as non-responders.

#### 2.2.3. TCGA

TCGA (The Cancer Genome Atlas) is a large-scale cancer genome project led by the National Cancer Institute and other institutions, providing clinical information and multi-omics data from more than 11,000 patients across 33 cancer types [18]. We used TCGA data for two purposes. First, during training, TCGA served as the target domain (unlabeled data) for domain adaptation from cell lines to patients, aiming to correct the discrepancy in gene expression distributions between the two data sources. Second, during evaluation, patient cohorts who received specific chemotherapy were extracted from TCGA and used as test data to assess model prediction performance (total n = 145; Supplementary Table S1).

#### 2.2.4. GEO

GEO (Gene Expression Omnibus) is a public gene expression database operated by NCBI, providing gene expression data from microarrays and RNA-seq along with associated clinical information [19]. During the evaluation phase, patient cohorts who received specific chemotherapy were extracted from GEO and used as test data to assess model prediction performance (total n = 673; Supplementary Table S1).

### 2.3. Overview of CODE-AE algorithm and performance

#### 2.3.1. CODE-AE

CODE-AE (Context-aware Deconfounding AutoEncoder) [11] is a domain adaptation method developed to predict patient drug response from cell line data. Its core design is intended to explicitly separate biological signals shared between cell line and patient samples (“Shared Embedding”) from features specific to each domain (“Private Embedding”). The conceptual model architecture is shown in Figure 1.

**Figure 1.**
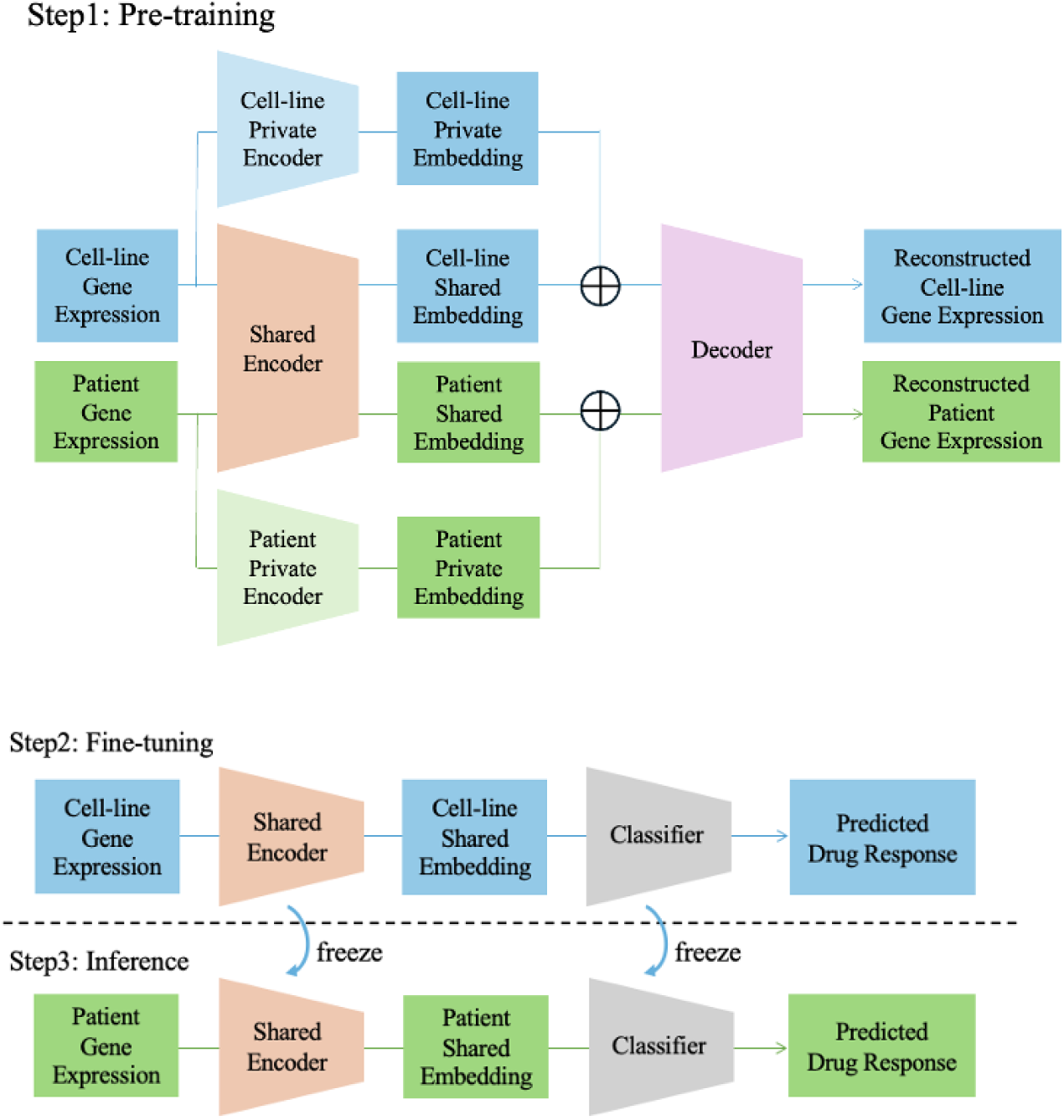
Architecture of CODE-AE (independently created based on Reference [11]).

CODE-AE consists of a three-stage training process. In the pre-training stage, gene expression data from both cell lines and patients are used as unlabeled data for unsupervised learning with an autoencoder [20]. An autoencoder learns to capture essential data features by compressing input data into a low-dimensional latent representation and then reconstructing the original input. Each sample is processed by two encoders— “Shared Encoder” and “Private Encoder” —which extract common and domain-specific features, respectively.

These two latent representations are concatenated and passed to a decoder that reconstructs the original gene expression profile, thereby encouraging the model to learn features of drug response shared across cell lines and patients.

In the fine-tuning stage, a classifier is connected to “Shared Encoder”, and supervised learning is performed using cell line drug response labels. “Shared Encoder” is not frozen at this stage, allowing the classifier to learn to predict drug response from features shared between patients and cell lines.

In the inference stage, the trained model is applied to patient samples to verify that a model trained solely on cell line data can generalize to patients.

The CODE-AE framework thus achieves drug response prediction for patients. This study uses CODE-AE as its foundation and aims to obtain insights for model improvement through detailed analyses of its prediction characteristics and training data composition.

#### 2.3.2. Evaluation of CODE-AE baseline performance

We started by examining the prediction behavior of CODE-AE using the original authors’ implementation. One particular point of interest was a scenario of training on a diverse set of cancer types, where we wanted to confirm whether CODE-AE captures essential biological signals of drug sensitivity or confounding factors—such as cancer-type-specific gene expression patterns— rather than just relying on the macro-level patterns for its predictions. If CODE-AE learned to exploit cancer type differences, integrated evaluation metrics across all cancer types will tend to appear favorable, yet the model will still fail to correctly distinguish drug response among individual patients within each cancer type. We therefore scrutinized the prediction accuracy for each cancer type and performed a multifaceted analysis of the features learned by CODE-AE.

##### Evaluation of prediction accuracy by cancer type

We first examined the extent to which the model captures within-cancer-type heterogeneity. The original paper reported only pan-cancer evaluation, but this metric may be overestimating true predictive performance by merely capturing systematic differences between cancer types. We therefore calculated AUROC for each cancer type individually to quantify how well drug response prediction performs within each cancer type.

##### Comparison of prediction score distributions

If the first analysis revealed a discrepancy between pan-cancer and cancer type-specific evaluation, the model may be using cancer type identity rather than individual sensitivity as a basis for prediction. To test this hypothesis, we visualized the distribution of prediction scores for each cancer type and examined whether distributions are systematically separated across cancer types.

##### Analysis of class distribution in training data

If systematic bias by cancer type was observed in the prediction score distributions, the underlying cause may reside not in the model architecture itself but in the composition of the training data. We therefore used drug response data from GDSC to calculate and compare the proportion of drug response labels across cancer types, evaluating the impact of inherent imbalances in the training data on model learning.

Through these analyses, we aimed to verify whether the prediction performance reported in the original paper [11] genuinely reflects drug response prediction or is an artifact of inter-cancer-type differences, and to inform the selection of appropriate improvement strategies.

### 2.4. Proposed Method: Optimization of Training Data Composition

To address the class imbalance in hematological cancers and the resulting shortcut learning, we examined two main approaches (Table 1).

**Table 1.** Main approaches.

| **Table 1. Main approaches.** |  |
| --- | --- |
| **Approach** | **Overview** |
| Hematological Cancer Exclusion | Exclude hematological cancers from training data and train on solid cancers only |
| Class Balance Adjustment | Retain hematological cancers while correcting class imbalance for training |

#### 2.4.1. Approach 1: Exclusion of Hematological Cancers

The first approach excludes all hematological cancers from the training data and trains the model solely on solid cancers. This is based on the hypothesis that the large biological differences between hematological and solid cancers drive shortcut learning, and that removing hematological cancers will encourage the model to learn biological features of drug response common to solid cancers rather than to identify cancer types. More specifically, based on metadata from CCLE, GDSC, and TCGA, all cell lines and patient samples classified as hematological cancers (leukemia, lymphoma, myeloma) were removed from the training dataset. Of the 1,305 total cell line samples used for CODE-AE training, 205 (approximately 15.7%) were hematological cancer cell lines, which were removed when we took this approach.

#### 2.4.2. Approach 2: Class Balance Adjustment

The second approach retains hematological cancers in the training data while adjusting the class balance. Although hematological and solid cancers differ greatly at the tissue level, they share common drug mechanisms of action at the cellular level. This approach is based on the hypothesis that appropriately correcting class imbalance enables the model to learn these shared drug response mechanisms rather than cancer type differences. Two techniques were used for class balance adjustment:

##### Oversampling (Duplication)

This is a standard technique for class imbalance problems [21] that adjusts the class ratio by duplicating samples from the minority class. We adopted a straightforward implementation in which non-responder samples were duplicated until their proportion of the overall dataset reached a specified target value.

##### Class Weighting

This technique assigns greater weight to the minority class in the loss function [21]. We used the inverse of each class’s sample count as a class weight in the loss function, thereby correcting the imbalanced data distribution at the loss-function level.

#### 2.4.3. Exploration of Hyperparameter Space

To identify the optimal parameter settings for the class balance adjustment approach, we examined the combinations shown in Table 2. Three parameters were varied:

**Table 2.** Different training data compositions. ‘Blood’ refers to hematological cancers. Blood/All = adjustment target, number (30%/40%/50%) = minority class ratio.

| Model Name | Adjustment Target | Minority Class Ratio | Class Weighting |
| --- | --- | --- | --- |
| Exclude Blood | N/A (hematological cancers excluded) | N/A | N/A |
| Blood 50% | Hematological cancers only | 50% | No |
| Blood 40% Duplication | Hematological cancers only | 40% | No |
| Blood 30% Duplication | Hematological cancers only | 30% | No |
| Blood 40% Duplication + Class Weight | Hematological cancers only | 40% | Yes |
| Blood 30% Duplication + Class Weight | Hematological cancers only | 30% | Yes |
| Blood Class Weight Only | Hematological cancers only | N/A | Yes |
| All 50% Duplication | All cancer types | 50% | No |
| All 30% Duplication | All cancer types | 30% | No |
| All 30% Duplication + Class Weight | All cancer types | 30% | Yes |

##### Scope of class balance adjustment

We compared applying class balance adjustment to hematological cancers only versus applying it to all cancer types.

##### Minority class ratio

The minority class ratio was set to 30%, 40%, or 50% through oversampling. A ratio of 50% completely equalizes both classes, while lower ratios apply more conservative adjustments.

##### Combination of techniques

We examined oversampling alone, class weighting alone, and both techniques combined.

### 2.5 Analysis of Sample Classification Difficulty

To investigate the mechanism underlying the complementary effects of oversampling and class weighting, we analyzed how each correction method affects samples with different classification difficulty. We defined each sample’s difficulty based on a classification margin under the original CODE-AE model calculated as follows. For each sample, we computed the mean predicted score across all 25 classifiers (5 seeds × 5 classifiers). The classification margin was defined as: for a responder sample (True Label = 1), margin = mean predicted score − 0.5; for a non-responder sample (True Label = 0), margin = 0.5 − mean predicted score. Positive margins indicate correct classifications, with larger values signifying higher confidence. Conversely, negative margins indicate a sample is misclassified.

Within each condition (drug × cancer type), samples were divided into three difficulty groups based on margin tertiles: Easy (top tertile), Medium (middle tertile), and Hard (bottom tertile). To quantify the effect of each correction method, we computed the margin change (Δmargin) for each sample as the difference between the margin under the correction method and the margin under the original CODE-AE. A positive Δmargin indicates improvement, while a negative Δmargin indicates deterioration. The analysis was conducted separately for responder and non-responder samples.

### 2.6. Application to Additional Domain Adaptation Methods

To assess whether the effectiveness of the training strategies proposed in this study can be generalized beyond CODE-AE, we additionally applied the same strategies to two established pan-cancer domain adaptation methods: TRANSACT [10] and VAEN [9].

TRANSACT (Tumor Response Assessment by Nonlinear Subspace Alignment of Cell lines and Tumors) [10] is a domain adaptation framework that builds a consensus space capturing biological processes common to both preclinical models and human tumors. It first computes nonlinear principal components (NLPCs) independently for cell lines and tumors using kernel methods, then aligns these two sets of NLPCs by identifying pairs of highly similar processes termed Principal Vectors. The top Principal Vectors are interpolated to construct consensus features, onto which both the cell line and the tumor samples are projected. Drug response predictors trained on the projected cell line data are then applied to predict patients’ responses without requiring any labeled patient data during training [10].

VAEN (VAE model followed by Elastic Net) [9] is a deep generative model that imputes drug response across domains using a variational autoencoder framework. It learns a shared latent space from both labeled cell line data and unlabeled patient data by optimizing a reconstruction loss combined with a KL divergence regularization term. The learned latent representations capture the underlying biological variation across domains, enabling the transfer of drug response predictions from the cell lines to the patients [9].

For both methods, we applied the same training data composition and class balance adjustment strategies described in Section 2.4 and evaluated prediction performance on the same external patient cohorts from TCGA and GEO used for the CODE-AE evaluation.

### 2.7. Model Evaluation

#### 2.7.1. Statistical Metrics

##### Area Under the Receiver Operating Characteristic Curve (AUROC)

a metric for evaluating the discriminative performance of binary classification models. AUROC can be interpreted as the probability that a randomly selected positive sample receives a higher prediction score than a randomly selected negative sample.

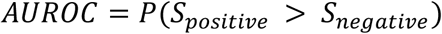

where 𝑆_𝑝𝑜𝑠𝑖𝑡𝑖𝑣𝑒_ and 𝑆_𝑛𝑒𝑔𝑎𝑡𝑖𝑣𝑒_ represent the prediction scores of positive and negative samples, respectively. AUROC ranges from 0 to 1, with values closer to 1 indicating better discriminative performance and 0.5 indicating performance equivalent to random prediction. We used AUROC to evaluate models’ ability to discriminate between responder and non-responder groups.

**Cohen’s d**: an effect size metric that expresses the standardized difference between the means of two groups [22]. Effect size quantifies the magnitude of an observed difference independently of statistical significance (p-value), enabling comparisons that are independent of sample size. Cohen’s d is calculated by the following formula.

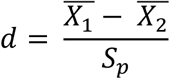

where 𝑋_1_ and 𝑋_2_ are the means of the two groups, and 𝑆_𝑝_ is the pooled standard deviation. The pooled standard deviation is calculated by the following formula.

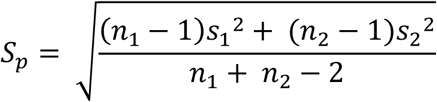

We obtained 25 AUROC values (5 seeds × 5 classifiers) for each model and evaluated the difference in AUROC distributions between the proposed model and the original CODE-AE using Cohen’s d, thereby providing an interpretable measure of the practical significance of any performance improvement.

#### 2.7.2. Statistical Tests

We used three statistical tests depending on the purpose of comparison, with a significance threshold of p < 0.05:

**Fisher’s exact test** [23]: a test that assesses independence between two categorical variables in a 2 × 2 contingency table. Because it calculates exact p-values based on the hypergeometric distribution, it is applicable even when sample sizes or expected frequencies are small. We used this test to evaluate the difference in the proportion of responder samples between hematological and solid cancers.

**Mann–Whitney U test** [24]: a nonparametric test for differences in the distributions of two independent groups. Because it is rank-based and does not assume normality, it is robust to outliers. We used this test to compare prediction score distributions between groups.

**Welch’s t-test** [25]: a parametric test for differences in the means of two independent groups. Unlike Student’s t-test [26], it estimates the variance of each group separately and adjusts the degrees of freedom via the Welch–Satterthwaite approximation, making it applicable when group variances differ. We used this test to compare prediction performance (AUROC) between models.

For each model, training was conducted with 5 random seeds, and 5 classifiers were obtained per seed through 5-fold cross-validation, yielding 25 AUROC values (5 seeds × 5 classifiers) per model. AUROC values from different models were compared as two groups using Welch’s t-test.

## 3. Results

### 3.1. Evaluation of the Original CODE-AE

First, to clarify whether the original CODE-AE learns essential features related to drug response or falls into shortcut learning based on cancer type identification, we performed a multifaceted analysis of its prediction characteristics on training data.

#### 3.1.1. Prediction Accuracy by Cancer Type

We used CCLE [6], which provides abundant response-labeled samples across diverse cancer types for individual drugs, to evaluate prediction accuracy by cancer type. Under pan-cancer evaluation (all cancer types combined), AUROC ranged from 0.57 to 0.68 depending on the drug. However, when the same data were evaluated by cancer type and the results were averaged, AUROC ranged from 0.53 to 0.61, which were significantly lower than pan-cancer AUROC for all drugs (Welch’s t-test, p < 0.05 for all drugs; Table 3). The gap between pan-cancer and cancer type-specific AUROC ranged from 0.02 to 0.11, with particularly large discrepancies for Temozolomide (difference: 0.11, p < 0.001) and Sorafenib (difference: 0.07, p < 0.001). One explanation for this consistent discrepancy may be that the model primarily identifies cancer types rather than individual drug sensitivity. Cancer type-specific evaluation may be eliminating this confounding and reflecting the model’s true drug sensitivity prediction capability.

**Table 3.** Comparison of prediction accuracy between pan-cancer evaluation and cancer type-specific evaluation. Values are expressed as mean ± standard deviation. n indicates the number of cancer types used for AUROC calculation. Significance values were determined using.

| Drug | Pan-cancer AUROC | Cancer Type-Specific AUROC (Mean) | p-value |
| --- | --- | --- | --- |
| 5-Fluorouracil | 0.61 ± 0.08 | 0.54 ± 0.06 (n=19) | $2.1 \times 10^{-3}$ |
| Cisplatin | 0.63 ± 0.02 | 0.61 ± 0.03 (n=20) | $4.1 \times 10^{-3}$ |
| Gemcitabine | 0.57 ± 0.05 | 0.53 ± 0.05 (n=20) | $2.8 \times 10^{-2}$ |
| Sorafenib | 0.63 ± 0.07 | 0.56 ± 0.08 (n=20) | $8.2 \times 10^{-4}$ |
| Temozolomide | 0.68 ± 0.05 | 0.57 ± 0.06 (n=20) | $2.4 \times 10^{-8}$ |

#### 3.1.2. Prediction Score Distribution and Training Data Class Imbalance

To test the hypothesis that the model primarily identifies cancer types rather than individual drug sensitivity, we analyzed prediction score distributions and the class composition of the training data. If the model is identifying cancer types, prediction score distributions should differ systematically across cancer types.

We compared the prediction score distributions for each combination of cancer type and drug, and found a clear separation of the scores between solid and hematological cancers. In this analysis, we constructed 25 predictors (5 seeds × 5 classifiers) for each drug and evaluated each predictor individually. Figure 2 shows one representative predictor for 5-Fluorouracil to illustrate this trend; the complete set of results for all 25 predictors across all five drugs is provided in Supplementary Figures S1. Across all five drugs, prediction scores of hematological cancer cell lines were consistently biased toward higher values (i.e., toward predicted sensitivity) compared to those of solid cancer cell lines. For all drugs, the majority of the 25 predictors (18 to 25) showed statistically significant differences in prediction score distributions between hematological and solid cancers (Mann–Whitney U test, p < 0.05; Table 4). Furthermore, Fisher’s exact test confirmed that for 4 of 5 drugs (5-Fluorouracil, Gemcitabine, Sorafenib, Temozolomide), the proportion of responder samples among hematological cancer cell lines was markedly higher than that of solid cancers in the training data (Table 5).

**Figure 2.**
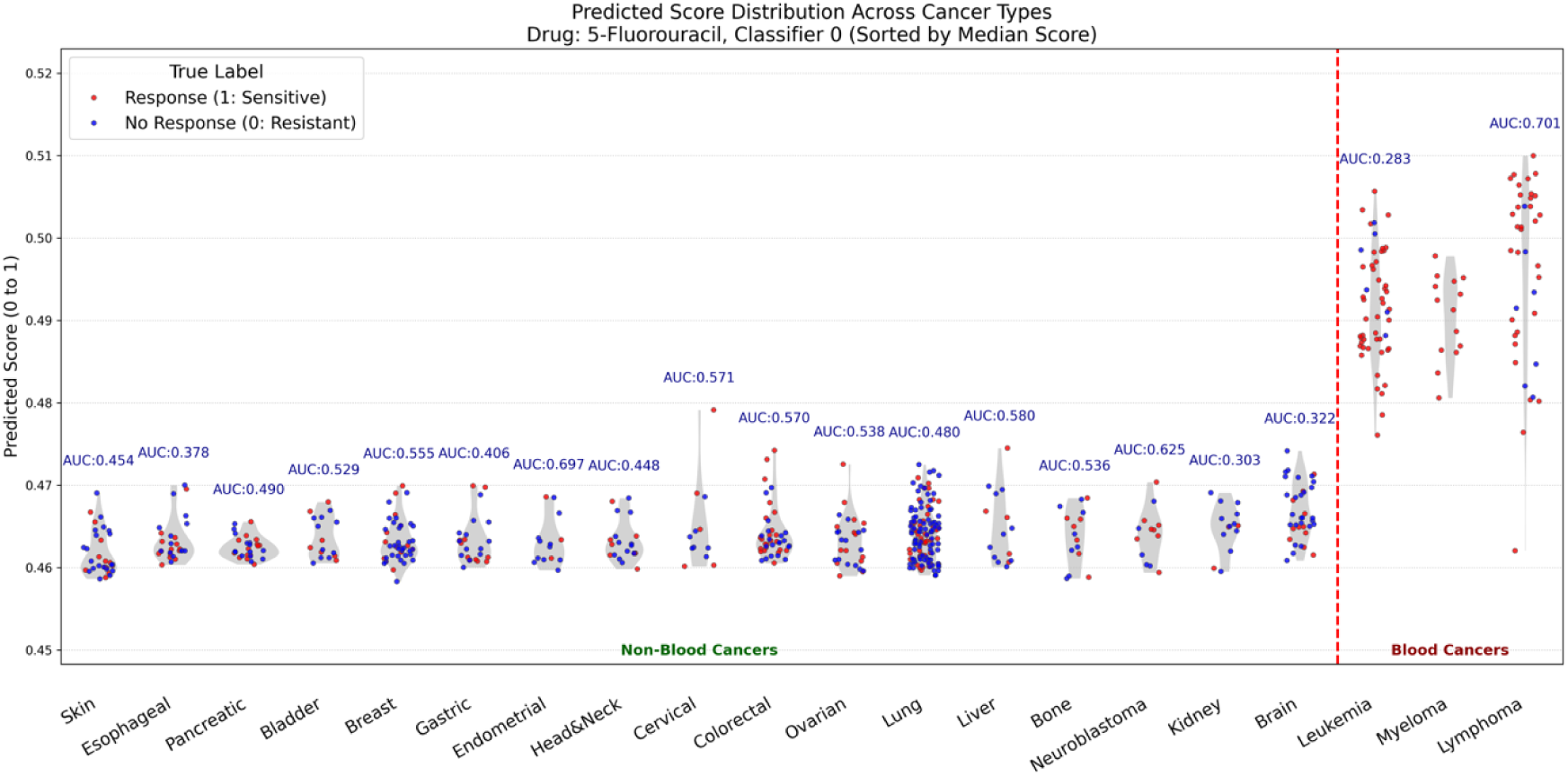
Comparison of prediction score distributions between hematological and solid cancers in CODE-AE for 5-Fluorouracil. Horizontal axis: cancer types (sorted by median score); Vertical axis: prediction score (0–1, higher = predicted response).

**Table 4.**
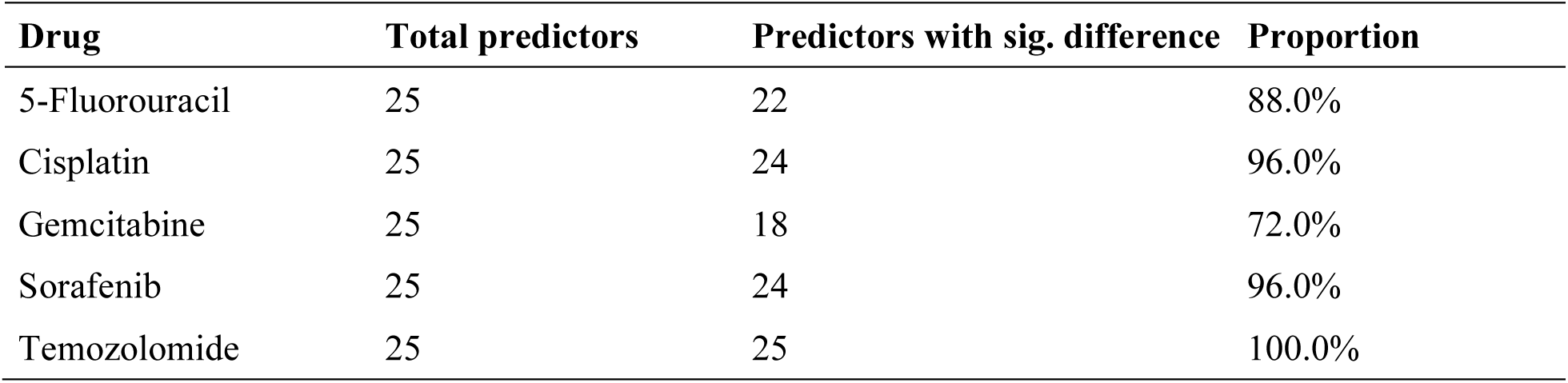
Differences in prediction score distributions between hematological and solid cancers.

| Drug | Total predictors | Predictors with sig. difference | Proportion |
| --- | --- | --- | --- |
| 5-Fluorouracil | 25 | 22 | 88.0% |
| Cisplatin | 25 | 24 | 96.0% |
| Gemcitabine | 25 | 18 | 72.0% |
| Sorafenib | 25 | 24 | 96.0% |
| Temozolomide | 25 | 25 | 100.0% |

**Table 5.** Comparison of the proportion of responder samples between hematological and solid cancers.

| Drug | Hematological (%) | Solid (%) | Difference (%) | p-value |
| --- | --- | --- | --- | --- |
| 5-Fluorouracil | 87.6 | 39.5 | 48.1 | $8.0 \times 10^{-21}$ |
| Cisplatin | 57.0 | 48.3 | 8.7 | 0.060 |
| Gemcitabine | 65.5 | 48.6 | 16.9 | $8.5 \times 10^{-4}$ |
| Sorafenib | 73.1 | 41.3 | 31.8 | $2.9 \times 10^{-9}$ |
| Temozolomide | 86.7 | 41.0 | 45.7 | $7.7 \times 10^{-19}$ |

These results suggest the following mechanism underlying the model’s predictions: because training data contained a disproportionately high fraction of responder samples among hematological cancers, the model learned to identify whether a sample is of hematological origin and to assign it a high prediction score (i.e., predicted response). This strategy achieves reasonable pan-cancer AUROC but is unable to capture the true differences in drug sensitivity within individual cancer types.

### 3.2. Accuracy Comparison Between Proposed Methods and Original CODE-AE

We compared two training strategies: exclusion of hematological cancers and class balance adjustment during retaining.

#### 3.2.1. Prediction Performance on CCLE

We first evaluated prediction performance on the cell line dataset (CCLE). The analysis was restricted to (drug, cancer type) pairs with 30 or more samples, yielding 25 pairs across 5 drugs and 6 cancer types (brain, breast, colorectal, lung, ovarian, skin). Table 6 summarizes changes in prediction performance relative to the original CODE-AE. Among all models tested, those combining oversampling with class weighting on hematological cancers—“Blood 30% Duplication + Class Weight” (7 significant improvements, 0 significant performance decreases; Table 6, Figure 3a) and “Blood 40% Duplication + Class Weight” (8 significant improvements and 1 significant decrease; Table 6, Figure 3b) —achieved the greatest net improvement. In contrast, “Exclude Blood”, which completely excluded hematological cancers, showed more significant worsening than improvements (five improvements vs. seven decreases; Table 6, Figure 3c), indicating that simply removing hematological cancers is not the optimal strategy. Furthermore, models combining oversampling with class weighting outperformed those using either technique alone (Table 6), suggesting that the two techniques have complementary effects: oversampling stabilizes training by reducing extreme class ratios, while class weighting ensures a minority class receives appropriate emphasis in the loss function. Models that extended class balance adjustment to all cancer types (“All 50% Duplication”, “All 30% Duplication”, “All 30% Duplication + Class Weight”) performed worse than those targeting only hematological cancers (Table 6). This is likely because enforcing a uniform class ratio across all cancer types distorted the native biological characteristics of individual cancer types and introduced noise.

**Figure 3.**
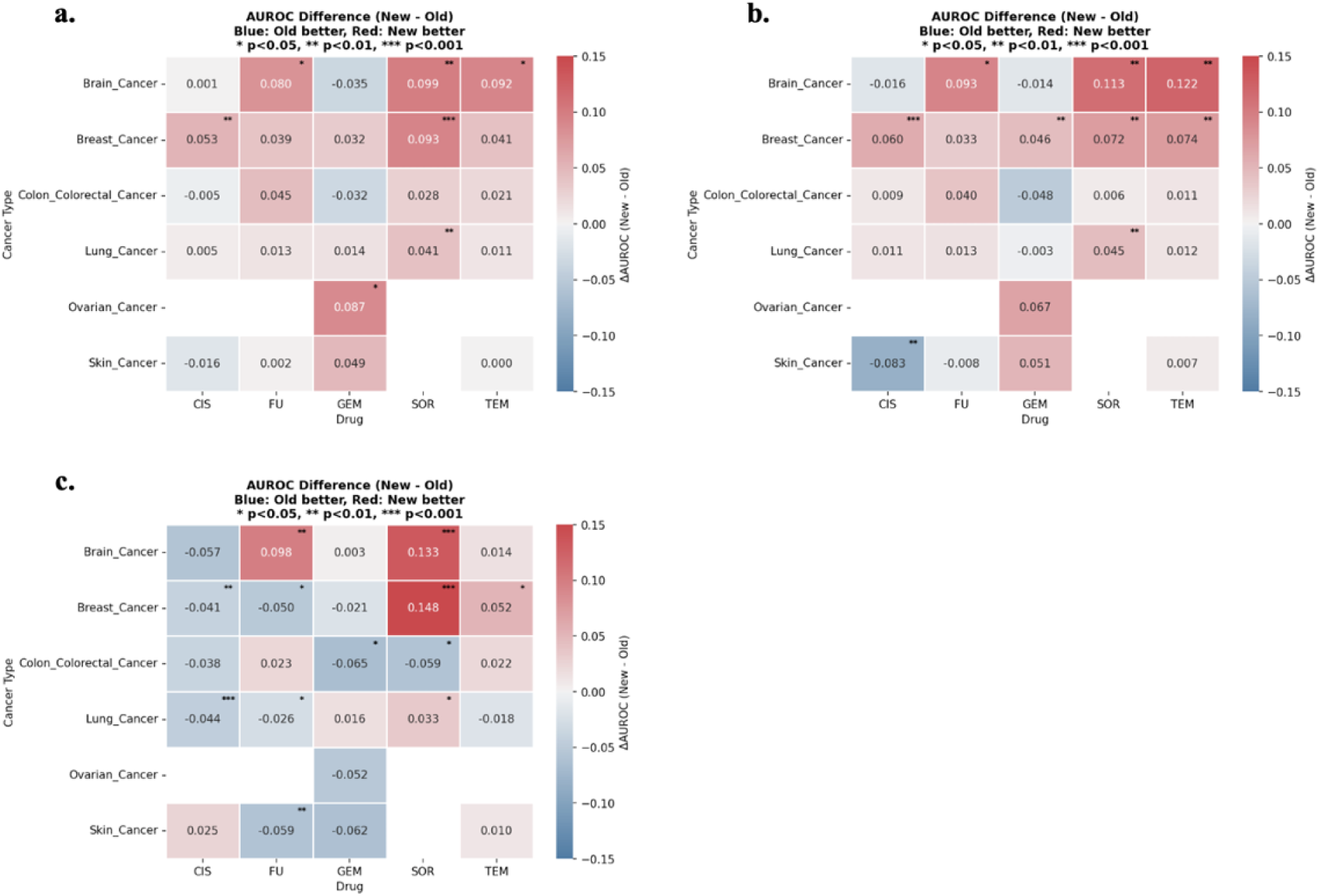
Comparison of prediction accuracy on CCLE cell line data. Red: proposed model superior; Blue: original superior; *: p < 0.05. (a) Blood 30% Duplication + Class Weight, (b) Blood 40% Duplication + Class Weight, (c) Exclude Blood.

**Table 6.**
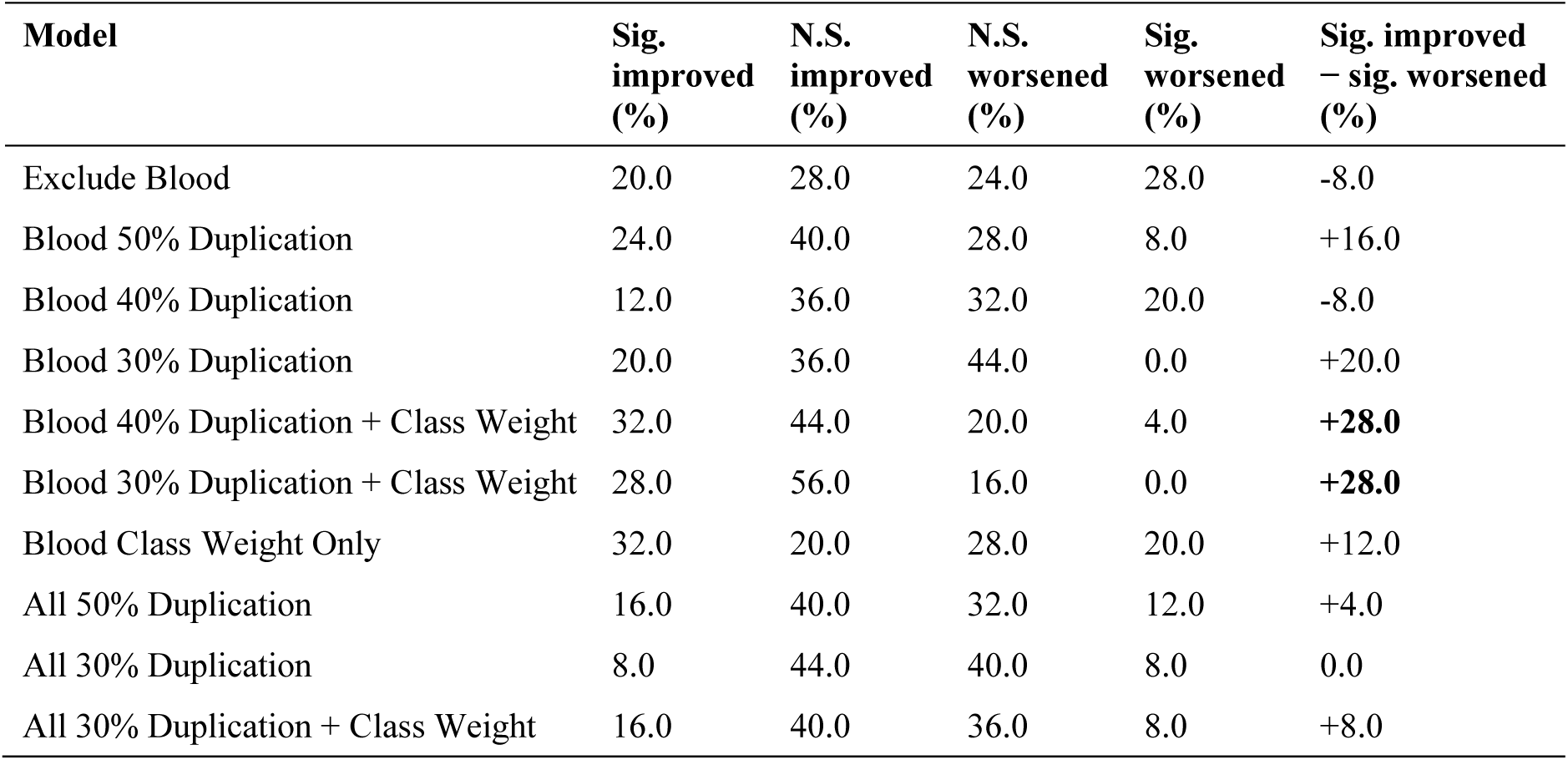
Changes in prediction performance for each proposed model on CCLE cell line data (n = 25 pairs). Sig.: p < 0.05; N.S.: no significant difference (Welch’s t-test). Imp. = Improved; Wors. = Worsened; Dup. = Duplication; CW = Class Weighting.

| Model | Sig. improved (%) | N.S. improved (%) | N.S. worsened (%) | Sig. worsened (%) | Sig. improved – sig. worsened (%) |
| --- | --- | --- | --- | --- | --- |
| Exclude Blood | 20.0 | 28.0 | 24.0 | 28.0 | -8.0 |
| Blood 50% Duplication | 24.0 | 40.0 | 28.0 | 8.0 | +16.0 |
| Blood 40% Duplication | 12.0 | 36.0 | 32.0 | 20.0 | -8.0 |
| Blood 30% Duplication | 20.0 | 36.0 | 44.0 | 0.0 | +20.0 |
| Blood 40% Duplication + Class Weight | 32.0 | 44.0 | 20.0 | 4.0 | **+28.0** |
| Blood 30% Duplication + Class Weight | 28.0 | 56.0 | 16.0 | 0.0 | **+28.0** |
| Blood Class Weight Only | 32.0 | 20.0 | 28.0 | 20.0 | +12.0 |
| All 50% Duplication | 16.0 | 40.0 | 32.0 | 12.0 | +4.0 |
| All 30% Duplication | 8.0 | 44.0 | 40.0 | 8.0 | 0.0 |
| All 30% Duplication + Class Weight | 16.0 | 40.0 | 36.0 | 8.0 | +8.0 |

#### 3.2.2. Samples’ Difficulty Analysis for Clarifying the Mechanisms of the Combined Correction Method

To elucidate why the combination of oversampling and class weighting outperforms either technique alone, we examined how each correction method affects samples with varying classification difficulty, using the same 25 (drug × cancer type) pairs across six solid cancer types described in Section 3.2.1.

All correction methods produced a consistent pattern: margins improved for non-responder samples (mean Δmargin ranging from +0.0152 to +0.0378) while deteriorating for responder samples (mean Δmargin ranging from −0.0178 to −0.0387) (Table 7, Figure 4). This indicates that correcting the class imbalance in hematological cancers shifts the model’s decision boundary toward predicting non-response, thereby improving the classification of non-responders at the expense of responders in solid cancers.

**Figure 4.**
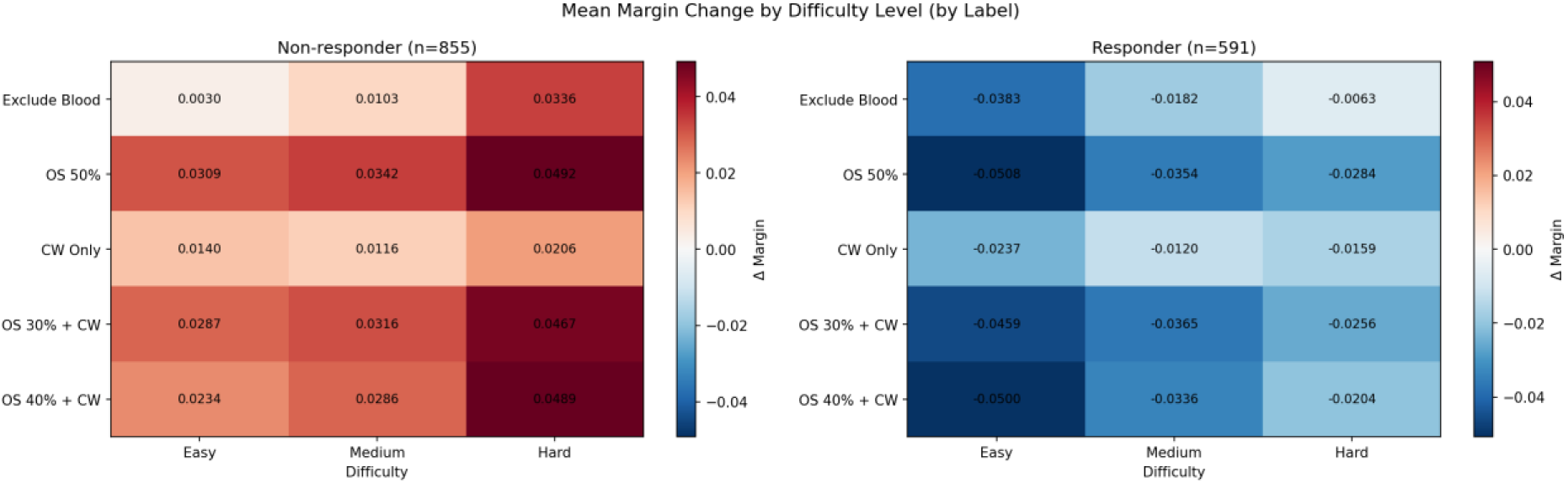
Mean margin change (Δmargin) by sample difficulty level for non-responder (left) and responder (right) samples across five correction methods. Red indicates improvement (positive Δmargin) and blue indicates deterioration (negative Δmargin).

**Table 7.** Mean margin change (Δmargin) by sample difficulty level for each correction method, analyzed separately for non-responder and responder samples. Non-responder samples (n = 855)

| Method | Difficulty | n | Δmargin (mean ± SD) | Improvement Rate |
| --- | --- | --- | --- | --- |
| Exclude Blood | Easy | 274 | $0.0030 \pm 0.0126$ | 69.0% |
| | Medium | 312 | $0.0103 \pm 0.0097$ | 83.3% |
| | Hard | 269 | $0.0336 \pm 0.0145$ | 100.0% |
| | All | 855 | $0.0153 \pm 0.0177$ | 84.0% |
| Blood 50% Duplication | Easy | 274 | $0.0309 \pm 0.0086$ | 99.6% |
| | Medium | 312 | $0.0342 \pm 0.0124$ | 100.0% |
| | Hard | 269 | $0.0492 \pm 0.0170$ | 100.0% |
| | All | 855 | $0.0378 \pm 0.0152$ | 99.9% |
| Blood Class Weight Only | Easy | 274 | $0.0140 \pm 0.0267$ | 78.8% |
| | Medium | 312 | $0.0116 \pm 0.0211$ | 70.2% |
| | Hard | 269 | $0.0206 \pm 0.0193$ | 85.9% |
| | All | 855 | $0.0152 \pm 0.0228$ | 77.9% |
| Blood 30% Duplication<br>+ Class Weight | Easy | 274 | $0.0287 \pm 0.0097$ | 100.0% |
| | Medium | 312 | $0.0316 \pm 0.0158$ | 100.0% |
| | Hard | 269 | $0.0467 \pm 0.0181$ | 100.0% |
| | All | 855 | $0.0354 \pm 0.0169$ | 100.0% |
| Blood 40% Duplication<br>+ Class Weight | Easy | 274 | $0.0234 \pm 0.0137$ | 100.0% |
| | Medium | 312 | $0.0286 \pm 0.0203$ | 100.0% |
| | Hard | 269 | $0.0489 \pm 0.0237$ | 100.0% |
| | All | 855 | $0.0333 \pm 0.0224$ | 100.0% |

Responder samples (n = 591)
| Method | Difficulty | n | Δmargin (mean ± SD) | Improvement Rate |
| --- | --- | --- | --- | --- |
| Exclude Blood | Easy | 221 | $-0.0383 \pm 0.0175$ | 0.0% |
| | Medium | 161 | $-0.0182 \pm 0.0103$ | 2.5% |
| | Hard | 209 | $-0.0063 \pm 0.0100$ | 23.4% |
| | All | 591 | $-0.0215 \pm 0.0192$ | 9.0% |
| Blood 50% Duplication | Easy | 221 | $-0.0508 \pm 0.0183$ | 0.0% |
| | Medium | 161 | $-0.0354 \pm 0.0146$ | 0.0% |
| | Hard | 209 | $-0.0284 \pm 0.0084$ | 0.0% |
| | All | 591 | $-0.0387 \pm 0.0174$ | 0.0% |
| Blood Class Weight Only | Easy | 221 | $-0.0237 \pm 0.0249$ | 19.9% |
| | Medium | 161 | $-0.0120 \pm 0.0155$ | 11.2% |
| | Hard | 209 | $-0.0159 \pm 0.0174$ | 11.5% |
| | All | 591 | $-0.0178 \pm 0.0207$ | 14.6% |
| Blood 30% Duplication<br>+ Class Weight | Easy | 221 | $-0.0459 \pm 0.0164$ | 0.0% |
| | Medium | 161 | $-0.0365 \pm 0.0164$ | 0.0% |
| | Hard | 209 | $-0.0256 \pm 0.0107$ | 0.0% |
| | All | 591 | $-0.0361 \pm 0.0170$ | 0.0% |
| Blood 40% Duplication<br>+ Class Weight | Easy | 221 | $-0.0500 \pm 0.0226$ | 0.0% |
| | Medium | 161 | $-0.0336 \pm 0.0216$ | 0.0% |
| | Hard | 209 | $-0.0204 \pm 0.0121$ | 0.0% |
| | All | 591 | $-0.0351 \pm 0.0230$ | 0.0% |

The magnitude of margin change was dependent on sample difficulty, but the direction was consistent within each class. For non-responder samples, the improvement was greatest for Hard samples and smallest for Easy samples across all methods (e.g., “Blood 30% Duplication + Class Weight”: Hard Δmargin = +0.0467, Medium = +0.0316, Easy = +0.0287). For responder samples, the deterioration was greatest for Easy samples and smallest for Hard samples (e.g., “Blood 30% Duplication + Class Weight”: Easy Δmargin = −0.0459, Medium = −0.0365, Hard = −0.0256). This gradient is consistent with a global decision boundary shift: samples closer to the original boundary are most affected, while samples far from the boundary are least affected.

Among the individual correction methods, class weighting alone exhibited a qualitatively different behavior from oversampling-based methods. While oversampling methods produced near-uniform improvement rates of 100% for non-responders, class weighting alone showed more variable improvement rates across difficulty levels (Easy: 78.8%, Medium: 70.2%, Hard: 85.9%). Notably, class weighting alone produced the smallest deterioration in responder samples (mean Δmargin = −0.0178 vs. −0.0387 for “Blood 50% Duplication” and −0.0361 for “Blood 30% Duplication + Class Weight”). Drug-specific analysis revealed that for Temozolomide and Cisplatin, class weighting alone improved responder margins in certain difficulty tiers where all other methods showed no improvement, suggesting that class weighting plays a unique role in preserving responder classification for specific drugs.

These results do not support the hypothesis that oversampling and class weighting improve distinct difficulty subsets of samples. Rather, the analysis suggests that their complementary effects operate through different mechanisms: oversampling provides a large, stable shift in the decision boundary that substantially improves non-responder classification, but this aggressive shift comes at a greater cost to responder classification. Class weighting alone produces a more conservative boundary adjustment that better preserves responder classification but provides less consistent improvement for non-responders. When combined, oversampling establishes the primary boundary correction while class weighting modulates the training dynamics to mitigate the excessive deterioration.

#### 3.2.3. Prediction Performance on TCGA and GEO External Patient Cohorts

We next evaluated the generalization of CODE-AE models, trained under different strategies, to clinical data using external patient cohorts from TCGA and GEO. Table 8 summarizes, for each of 11 patient cohorts, how each method’s prediction performance changed relative to the original CODE-AE. Among the class balance approaches, “Blood 30% Duplication + Class Weight” and “Blood Class Weight Only” achieved the strongest results, with significant accuracy improvements in five of 11 cohorts and no significant deterioration (Figure 5). In contrast, hematological cancer exclusion (“Exclude Blood”) and global class balance adjustment (“All 50% Duplication”, “All 30% Duplication”, “All 30% Duplication + Class Weight”) resulted in more significant worsening than improvements.

**Figure 5.**
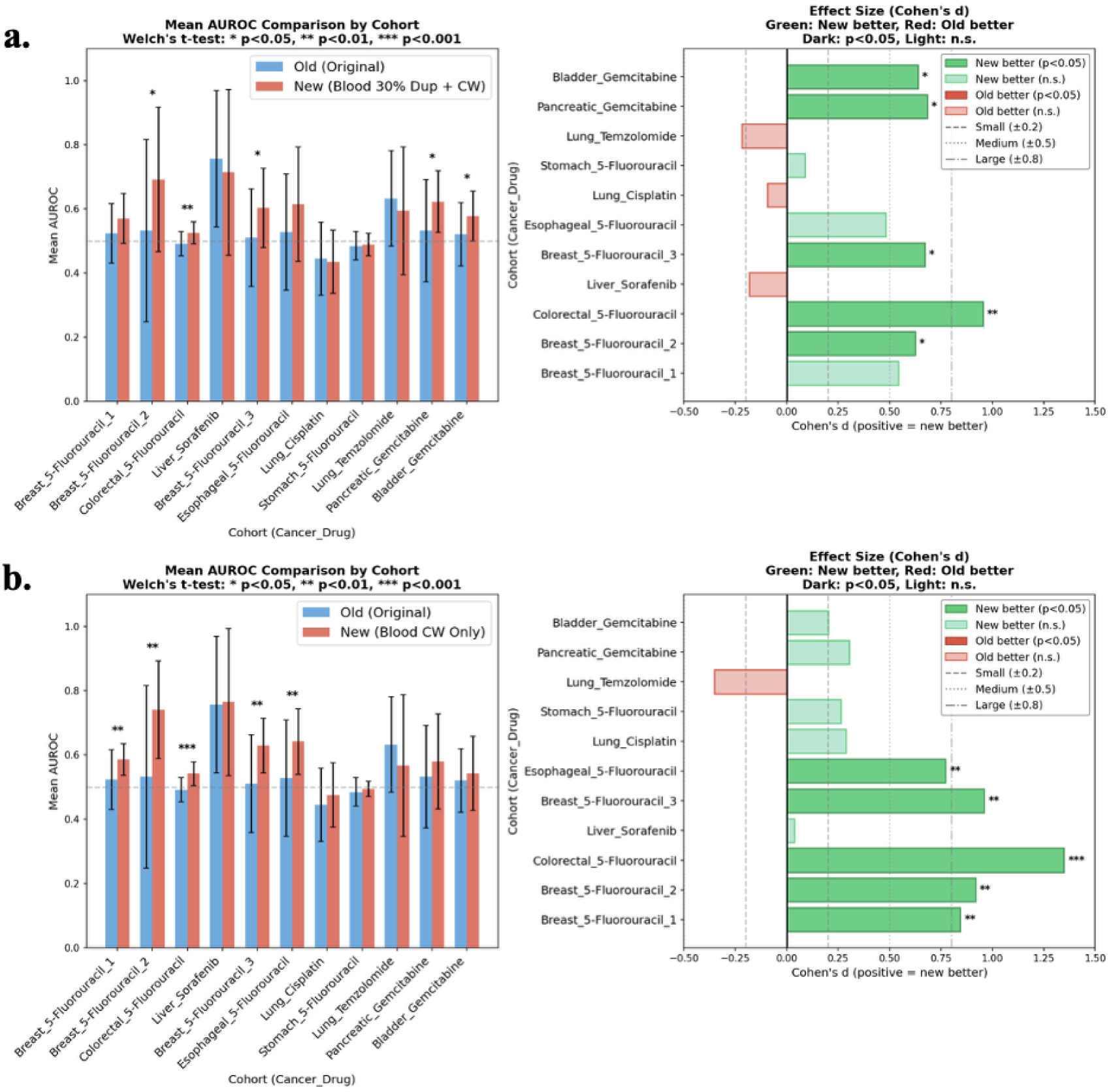
Comparison of prediction accuracy in external patient cohorts. Left: AUROC; Right: Effect size (Cohen’s d). Dark color: p < 0.05. (a) Blood 30% Duplication + Class Weight, (b) Blood Class Weight Only.

**Table 8.** Changes in prediction performance in external patient cohorts (TCGA/GEO, n = 11). Abbreviations as in. Table 6.

| Model | Sig. improved (%) | N.S. improved (%) | N.S. worsened (%) | Sig. worsened (%) | Sig. improved – sig. worsened (%) |
| --- | --- | --- | --- | --- | --- |
| Exclude Blood | 9.1 | 36.4 | 36.4 | 18.2 | -9.1 |
| Blood 50% Duplication | 27.3 | 45.5 | 27.3 | 0.0 | +27.3 |
| Blood 40% Duplication | 18.2 | 63.6 | 9.1 | 9.1 | +9.1 |
| Blood 30% Duplication | 9.1 | 72.7 | 18.2 | 0.0 | +9.1 |
| Blood 40% Duplication + Class Weight | 36.4 | 36.4 | 27.3 | 0.0 | +36.4 |
| Blood 30% Duplication + Class Weight | 45.5 | 27.3 | 27.3 | 0.0 | **+45.5** |
| Blood Class Weight Only | 45.5 | 45.5 | 9.1 | 0.0 | **+45.5** |
| All 50% Duplication | 9.1 | 63.6 | 9.1 | 18.2 | -9.1 |
| All 30% Duplication | 0.0 | 72.7 | 18.2 | 9.1 | -9.1 |
| All 30% Duplication + Class Weight | 0.0 | 27.3 | 63.6 | 9.1 | -9.1 |

These results demonstrate that appropriately controlling shortcut learning in hematological cancers is effective not only for cell line data but also for predicting clinical outcomes in patient data from a different domain.

### 3.3. Generalizability of the Proposed Strategy to Other Pan-Cancer Domain Adaptation Methods

#### 3.3.1 Generalizability to TRANSACT: Consistent Improvement via Class Balance Adjustment

Results obtained using TRANSACT were generally consistent with those observed in CODE-AE. As shown in Figure 6, all four class balance adjustment strategies tested—“Blood Class Weight Only,” “Blood 50% Duplication,” “Blood 30% Duplication + Class Weight,” and “Blood 40% Duplication + Class Weight”—improved prediction accuracy over the original model on external patient cohorts. Among them, “Blood Class Weight Only” (𝚫AUROC = +0.0267), “Blood 30% Duplication + Class Weight” (𝚫AUROC = +0.0262), and “Blood 40% Duplication + Class Weight” (𝚫AUROC = +0.0251) achieved the greatest improvements. In contrast, “Exclude Blood” showed a substantial decrease in prediction accuracy (𝚫AUROC = −0.0667), confirming that complete exclusion of imbalanced cancer subtypes is consistently detrimental across the different domain adaptation frameworks. These results suggest that the effectiveness of class imbalance correction is not specific to the CODE-AE architecture but is broadly applicable to domain adaptation methods that enforce explicit distributional alignment between cell lines and patients.

**Figure 6.**
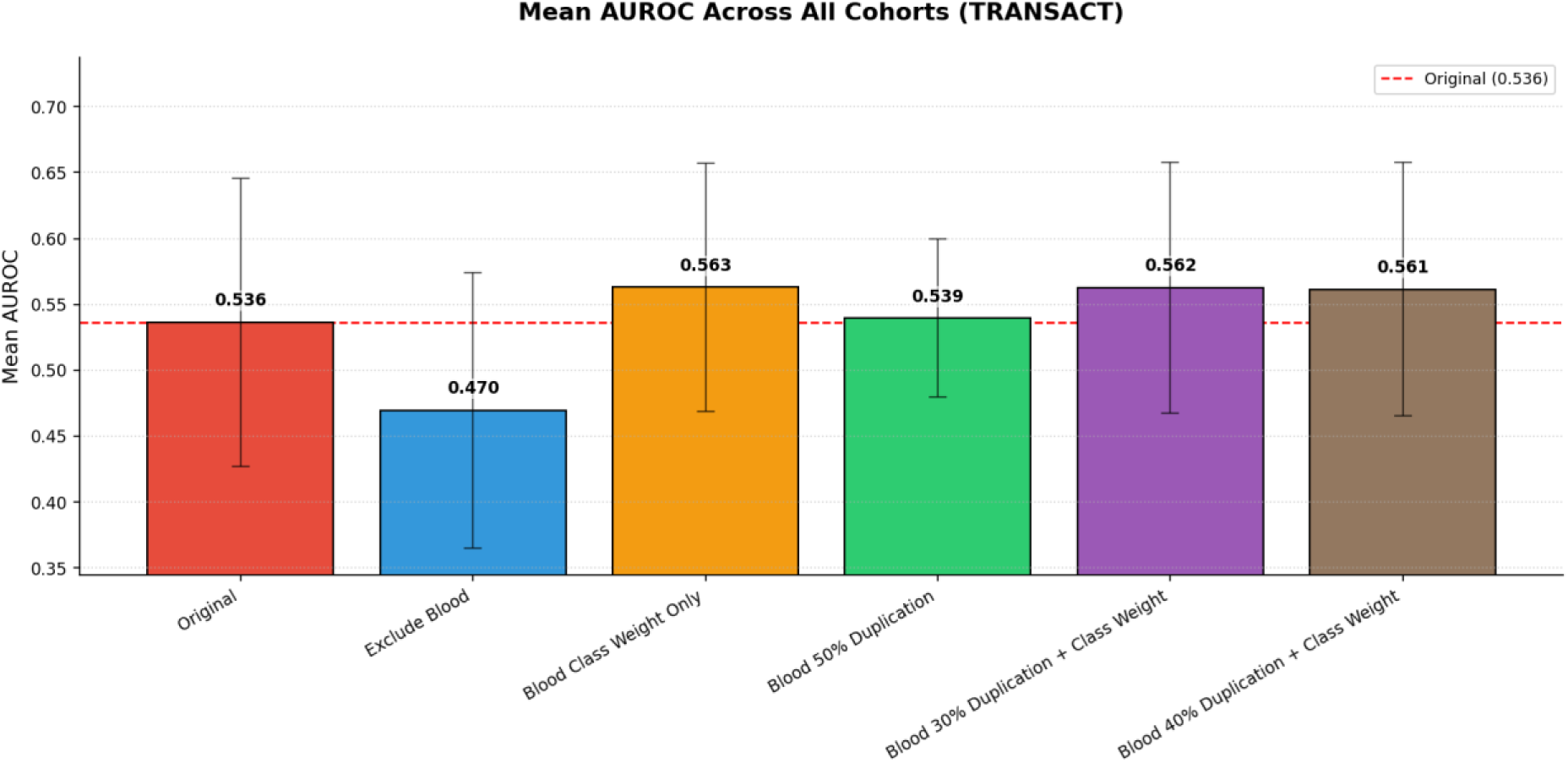
Average AUROC across all external patient cohorts for the original TRANSACT model and representative training strategies proposed in this study.

#### 3.3.2 Limitations of the Proposed Strategy: The Case of VAEN

In contrast, for VAEN, the original model without any training data optimization achieved the highest prediction accuracy (Figure 7). One possible explanation lies in the design of VAEN: unlike CODE-AE and TRANSACT, VAEN lacks a strong constraint to directly align the distributions of cell lines and patient tumors, and the authors themselves acknowledged the presence of numerous confounding factors in the learned representations. This suggests that the benefit of training data composition optimization may be contingent on the degree to which a method enforces distributional alignment between domains. In domain adaptation approaches that strongly align cell line and patient tumor distributions, correcting cancer-type-specific class imbalances in training data can be an effective strategy for improving prediction performance. Therefore, when applying training data optimization strategies, it is important to consider the underlying domain adaptation mechanism of the target method.

**Figure 7.**
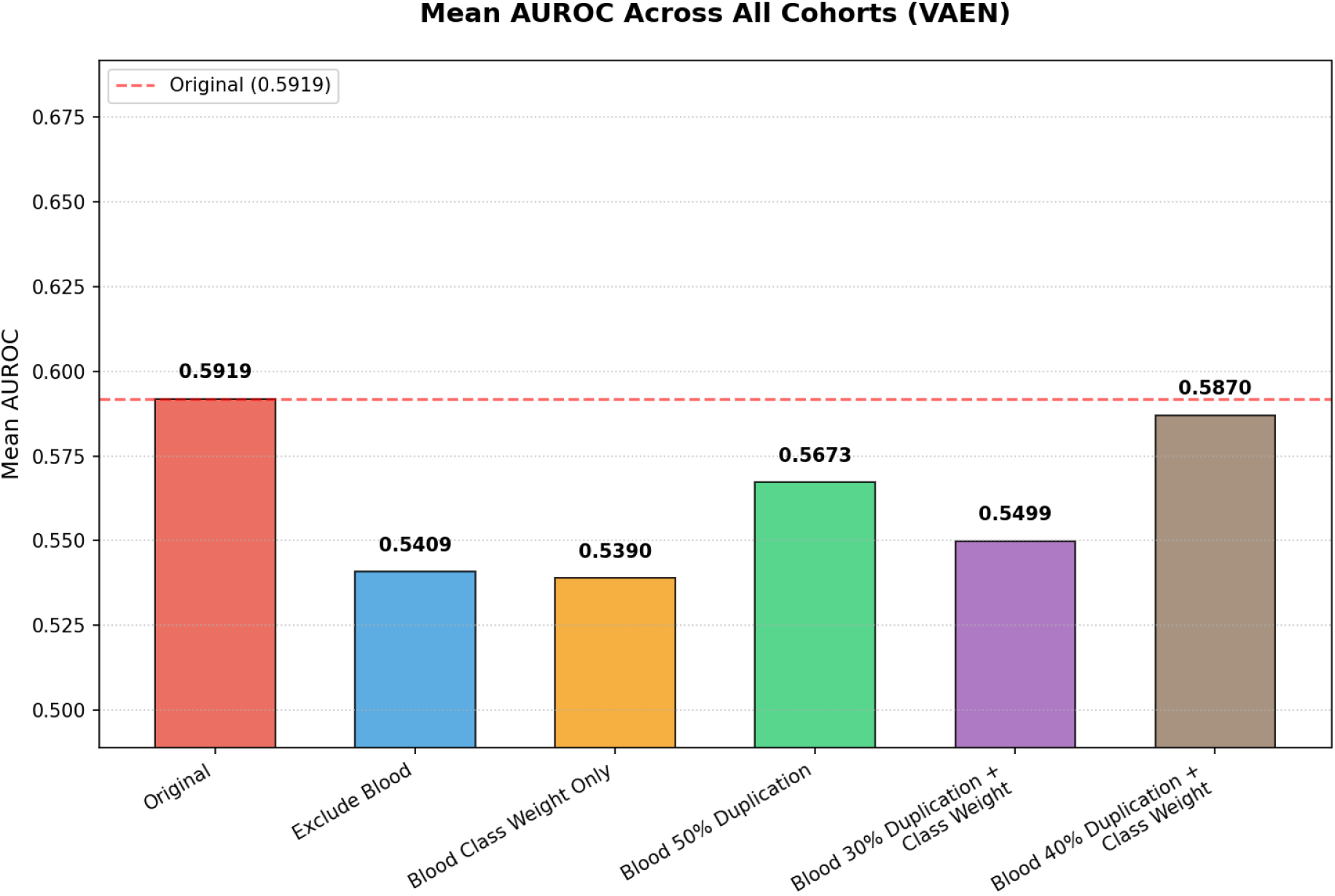
Average AUROC across all external patient cohorts for the original VAEN model and representative training strategies proposed in this study.

## 4. Discussion

In this study, we revealed that class imbalance between hematological and solid cancers causes shortcut learning in CODE-AE, a model that predicts patient drug response from cell line data, under the original authors’ training protocol. To address this problem, we proposed and evaluated methods for optimizing training data composition. Below, we discuss the interpretation and significance of the results, as well as the limitations of this study and future directions.

### 4.1. Effective Utilization of Hematological Cancer Data

Of the two approaches examined, although completely excluding hematological cancers from training improved accuracy for some of the cell line and patient cohorts, it caused performance degradation in many others, producing inconsistent results overall. In contrast, retaining hematological cancers while adjusting class balance achieved more consistent accuracy improvements across both cell lines and patient cohorts.

This finding suggests that hematological cancer data contain information relevant to drug response prediction and should be retained rather than excluded from pan-cancer training. The superior performance of class balance adjustment indicates that these data can be effectively utilized when their class imbalance is appropriately corrected. This result provides a practical guideline for improving pan-cancer drug response prediction using limited clinical data.

### 4.2. Complementary Effects of Oversampling and Class Weighting

Among the tested class balance adjustment models, those combining oversampling with class weighting generally outperformed those using either technique alone. In particular, “Blood 30% Duplication + Class Weight” demonstrated the highest prediction performance on both cell line and external patient cohorts (Tables 6 and 8).

When oversampling alone was applied (“Blood 50/40/30% Duplication”), increasing the minority class ratio to 50% may deviate excessively from the original class distribution in the training data. On the other hand, limiting the ratio to 30% or 40% may leave a residual class imbalance insufficiently corrected.

When class weighting alone was applied (“Blood Class Weight Only”), the original training data distribution was preserved, while the contribution of the minority class to the loss function was increased. This method achieved significant accuracy improvement in 5 of 11 patient cohorts, demonstrating strong generalization (Table 8). However, its improvement on CCLE cell line data was limited compared to the combined model (Table 6), likely because extremely skewed class ratios (e.g., 87.6% responder rate for 5-Fluorouracil in hematological cancers) produce extreme class weights, which may have overemphasized the minority class during optimization.

The combination of both techniques mitigated the extreme class ratio bias through oversampling to stabilize training while avoiding excessive duplication that would distort cancer type-specific drug response characteristics. Among the combined models tested at various oversampling ratios, “Blood 30% Duplication + Class Weight model”—with a non-responder ratio set at 30%—achieved the best accuracy: significant improvement in 7 of 25 pairs on CCLE (Table 6) and 5 of 11 cohorts on TCGA/GEO (Table 8). This result suggests that setting an appropriate oversampling ratio is crucial for maximizing the effect of class weighting while minimizing distortion of the data distribution.

### 4.3. Inefficiency of Global Class Balance Adjustment

Models that extended class balance adjustment to all cancer types (“All 50% Duplication”, “All 30% Duplication”, “All 30% Duplication + Class Weight”) showed poor overall prediction accuracy. Improvement on CCLE was extremely limited compared to models targeting only hematological cancers (Table 6), and performance on external patient cohorts was even worse than the original CODE-AE (Table 8).

These results indicate that class imbalance correction should be confined to hematological cancers, where the problem arises, and should not be applied uniformly to solid cancers.

### 4.4. Limitations and Future Perspectives

First limitation of this study is relatively modest absolute prediction accuracy, which therefore remains a concern. Although the proposed class balance adjustment strategy achieved statistically significant improvements over the original CODE-AE implementation, AUROC on external patient cohorts remained in the range of approximately 0.5 to 0.7. The pan-cancer approach is useful for compensating for the limited number of samples, but it inherently faces a trade-off in that it is difficult to fully capture the unique characteristics of each cancer type. For future clinical applications, appropriate use of pan-cancer and cancer type-specific models, or the development of hybrid approaches integrating both will be necessary. Additionally, integrating additional multi-omics data such as mutation profiles, copy number alterations, and methylation may improve capture of multiple biological facets of drug response and potentially increase prediction accuracy.

Second, the range of explored drugs was insufficiently diverse. We examined only five chemotherapeutic agents: 5-Fluorouracil, Cisplatin, Gemcitabine, and Temozolomide, which are conventional cytotoxic anticancer drugs, and Sorafenib, which is a small-molecule multikinase inhibitor. In contemporary cancer treatment, new modalities such as immune checkpoint inhibitors (anti-PD-1 antibodies, anti-CTLA-4 antibodies) and CAR-T cell therapy are rapidly becoming widely used, and the effectiveness of our approach for these drugs remains unverified. The response to immune checkpoint inhibitors, in particular, depends heavily on the tumor microenvironment and immune cell state in addition to tumor cell gene expression. Therefore, the current framework—which learns solely from the cell line data—may face fundamental limitations.

However, these novel agents have shorter clinical track records, and patient cohorts with accumulated drug response data are considerably smaller than those available for conventional chemotherapies. In this context, pan-cancer approaches like CODE-AE holds particular promise as they can secure sufficient training sample numbers by integrating data from multiple cancer types. For novel drug applications, constructing pan-cancer models that maximize limited data while applying the training data composition optimization identified in this study may be an effective strategy. Future work can verify the applicability of this approach to novel therapeutic agents and, to the extent it is needed, expand the feature sets and training data accordingly.

Some points that we would like to confirm in future investigations are whether the proposed strategy can be further improved by optimizing the selection of cancer types included during pan-cancer training. In addition, we would like to more comprehensively profile the applicability of this approach by evaluating it in a broader range of pan-cancer drug response prediction methods beyond CODE-AE and TRANSACT. Such studies may help establish general principles for improving domain adaptation-based drug response prediction.

### 4.5. Conclusion

In this study, we have explored an important challenge for pan-cancer based models where differences between individual cancer types and their representation in the data can adversely affect the results. In this selected case, a specific class imbalance between hematological and solid cancers caused shortcut learning in the pan-cancer drug response prediction model CODE-AE when trained according to the protocol developed by its original authors. To address this problem, we proposed methods for optimizing the training data composition. We demonstrated that rather than excluding hematological cancers, appropriately combining oversampling and class weighting can suppress shortcut learning while leveraging the valuable biological information contained in hematological cancer data. Importantly, we have observed that an original idea behind the method was a sound one and its effectiveness increased after the proposed changes.

Our findings demonstrate that in constructing drug response prediction models, training data composition and preprocessing have a substantial impact on prediction performance, beyond model architecture improvements alone. Careful attention to data characteristics and training strategy selection can maximize the utility of limited clinical data and ultimately advance precision medicine.

## References

[1] Gong J, Zhao Z, Niu X, Ji Y, Sun H, Shen Y, Chen B and Wu B (2025) AI reshaping life sciences: intelligent transformation, application challenges, and future convergence in neuroscience, biology, and medicine. Front. Digit. Health 7:1666415. doi: 10.3389/fdgth.2025.1666415

[2] Philip Ball. 2023. “Is AI Leading to a Reproducibility Crisis in Science?” Nature 624: 22–25.

[3] Muhammad A, Aka IT, Birdwell KA, Gordon AS, Roden DM, Wei WQ, Mosley JD, Van Driest SL. Genome-Wide Approach to Measure Variant-Based Heritability of Drug Outcome Phenotypes. Clin Pharmacol Ther. 2021 Sep;110(3):714–722. doi: 10.1002/cpt.2323. Epub 2021 Jul 12. PMID: 34151428; PMCID: PMC8376753.

[4] Li Y, Umbach DM, Krahn JM, Shats I, Li X, Li L. Predicting tumor response to drugs based on gene-expression biomarkers of sensitivity learned from cancer cell lines. BMC Genomics. 2021 Apr 15;22(1):272. doi: 10.1186/s12864-021-07581-7. PMID: 33858332; PMCID: PMC8048084.

[5] Geeleher P, Cox NJ, Huang RS. Clinical drug response can be predicted using baseline gene expression levels and in vitro drug sensitivity in cell lines. Genome Biol. 2014 Mar 3;15(3):R47. doi: 10.1186/gb-2014-15-3-r47. PMID: 24580837; PMCID: PMC4054092.

[6] Barretina, J., Caponigro, G., Stransky, N. et al. The Cancer Cell Line Encyclopedia enables predictive modelling of anticancer drug sensitivity. Nature 483, 603–607 (2012). 10.1038/nature11003

[7] Wanjuan Yang, Jorge Soares, Patricia Greninger, Elena J. Edelman, Howard Lightfoot, Simon Forbes, Nidhi Bindal, Dave Beare, James A. Smith, I. Richard Thompson, Sridhar Ramaswamy, P. Andrew Futreal, Daniel A. Haber, Michael R. Stratton, Cyril Benes, Ultan McDermott, Mathew J. Garnett, Genomics of Drug Sensitivity in Cancer (GDSC): a resource for therapeutic biomarker discovery in cancer cells, Nucleic Acids Research, Volume 41, Issue D1, 1 January 2013, Pages D955–D961, 10.1093/nar/gks1111

[8] Sharifi-Noghabi, H., Harjandi, P.A., Zolotareva, O. et al. Out-of-distribution generalization from labelled and unlabelled gene expression data for drug response prediction. Nat Mach Intell 3, 962–972 (2021). 10.1038/s42256-021-00408-w

[9] Jia, P., Hu, R., Pei, G. et al. Deep generative neural network for accurate drug response imputation. Nat Commun 12, 1740 (2021). 10.1038/s41467-021-21997-5

[10] S.M.C. Mourragui, M. Loog, D.J. Vis, K. Moore, A.G. Manjon, M.A. van de Wiel, M.J.T. Reinders, & L.F.A. Wessels, Predicting patient response with models trained on cell lines and patient-derived xenografts by nonlinear transfer learning, Proc. Natl. Acad. Sci. U.S.A. 118 (49) e2106682118, 10.1073/pnas.2106682118 (2021).

[11] He, D., Liu, Q., Wu, Y. et al. A context-aware deconfounding autoencoder for robust prediction of personalized clinical drug response from cell-line compound screening. Nat Mach Intell 4, 879–892 (2022). 10.1038/s42256-022-00541-0

[12] Li, Q., Qian, W., Zhang, Y. et al. A new wave of innovations within the DNA damage response. Sig Transduct Target Ther 8, 338 (2023). 10.1038/s41392-023-01548-8

[13] Seula Jeong, Yuheon Chung, Soomin Heo, Kyungjae Myung, Targeting DNA repair mechanisms in cancer therapy: the role of small molecule DNA repair inhibitors, NAR Cancer, Volume 7, Issue 4, December 2025, zcaf040, 10.1093/narcan/zcaf040

[14] Zhang H, Qureshi MA, Wahid M, Charifa A, Ehsan A, Ip A, De Dios I, Ma W, Sharma I, McCloskey J, Donato M, Siegel D, Gutierrez M, Pecora A, Goy A, Albitar M. Differential Diagnosis of Hematologic and Solid Tumors Using Targeted Transcriptome and Artificial Intelligence. Am J Pathol. 2023 Jan;193(1):51–59. doi: 10.1016/j.ajpath.2022.09.006. Epub 2022 Oct 13. PMID: 36243045.

[15] Montironi C, Muñoz-Pinedo C, Eldering E. Hematopoietic versus Solid Cancers and T Cell Dysfunction: Looking for Similarities and Distinctions. Cancers (Basel). 2021 Jan 14;13(2):284. doi: 10.3390/cancers13020284. PMID: 33466674; PMCID: PMC7828769.

[16] Zetian Zheng, Lei Huang, Fuzhou Wang, Linjing Liu, Jixiang Yu, Weidun Xie, Xingjian Chen, Xiangtao Li, Ka-Chun Wong. Drug Response Modeling across Cancers: Proteomics vs. Transcriptomics. bioRxiv 2024.12.04.626700; 10.1101/2024.12.04.626700

[17] Geirhos, R., Jacobsen, JH., Michaelis, C. et al. Shortcut learning in deep neural networks. Nat Mach Intell 2, 665–673 (2020). 10.1038/s42256-020-00257-z

[18] Tomczak K, Czerwińska P, Wiznerowicz M. The Cancer Genome Atlas (TCGA): an immeasurable source of knowledge. Contemp Oncol (Pozn). 2015;19(1A):A68–77. doi: 10.5114/wo.2014.47136. PMID: 25691825; PMCID: PMC4322527.

[19] Edgar, Ron et al. “Gene Expression Omnibus: NCBI gene expression and hybridization array data repository.” Nucleic acids research 30 1 (2002): 207–10.

[20] G. E. Hinton, R. R. Salakhutdinov, Reducing the Dimensionality of Data with Neural Networks.Science313,504–507(2006).DOI:10.1126/science.1127647

[21] H. He and E. A. Garcia, “Learning from Imbalanced Data,” in IEEE Transactions on Knowledge and Data Engineering, vol. 21, no. 9, pp. 1263–1284, Sept. 2009, doi: 10.1109/TKDE.2008.239.

[22] Cohen, J. (1988). Statistical Power Analysis for the Behavioral Sciences (2nd ed.). Routledge. 10.4324/9780203771587

[23] Fisher, R. A. “On the Interpretation of χ2 from Contingency Tables, and the Calculation of P.” Journal of the Royal Statistical Society, vol. 85, no. 1, 1922, pp. 87–94. JSTOR, 10.2307/2340521. Accessed 28 Dec. 2025.

[24] Mann, H. B., and D. R. Whitney. “On a Test of Whether One of Two Random Variables Is Stochastically Larger than the Other.” The Annals of Mathematical Statistics, vol. 18, no. 1, 1947, pp. 50–60. JSTOR, http://www.jstor.org/stable/2236101. Accessed 10 Dec. 2025.

[25] Welch, B. L. “The Generalization of ‘Student’s’ Problem When Several Different Population Variances Are Involved.” Biometrika, vol. 34, no. 1/2, 1947, pp. 28–35. JSTOR, 10.2307/2332510. Accessed 28 Dec. 2025.

[26] Student. “The Probable Error of a Mean.” Biometrika, vol. 6, no. 1, 1908, pp. 1–25. JSTOR, 10.2307/2331554. Accessed 28 Dec. 2025.

