## Supplementary Materials for "Training Strategy Optimization to Mitigate Shortcut Learning in Pan-Cancer Drug Response Prediction"

**Table S1. Summary of the TCGA and GEO patient datasets receiving specific anticancer drugs for independent testing.**

| Source | Drug name | Cancer type | Sample Size |
|---|---|---|---|
| GSE20194 | Cyclophosphamide+Doxorubicin+Fluorouracil+Paclitaxel | Breast cancer | 211 |
| GSE20271 | Cyclophosphamide+Doxorubicin+Fluorouracil | Breast cancer | 82 |
| GSE28702 | Fluorouracil+Leucovorin+Oxaliplatin | Colorectal cancer | 83 |
| GSE109211 | Sorafenib | Liver cancer | 67 |
| GSE6861 | Cyclophosphamide+Epirubicin+Fluorouracil | Breast cancer | 103 |
| GSE104958 | Cisplatin+Docetaxel+Fluorouracil | Esophageal cancer | 41 |
| GSE37138 | Bevacizumab+Cisplatin+Gemcitabine | Lung cancer | 42 |
| GSE14209 | Cisplatin+Fluorouracil | Stomach cancer | 44 |
| TCGA-LGG | Temozolomide | Brain cancer | 55 |
| TCGA-PAAD | Gemcitabine | Pancreatic cancer | 46 |
| TCGA-BLCA | Cisplatin+Gemcitabine | Urinary bladder cancer | 44 |

[5-Fluorouracil]

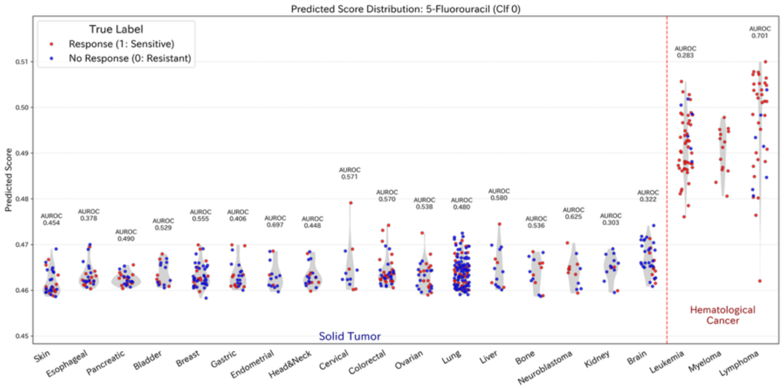


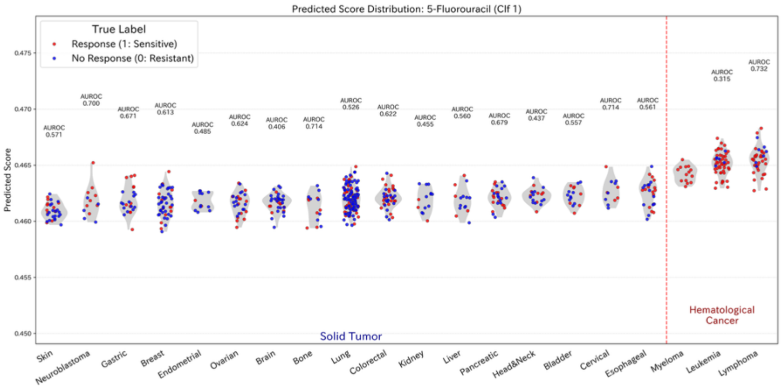


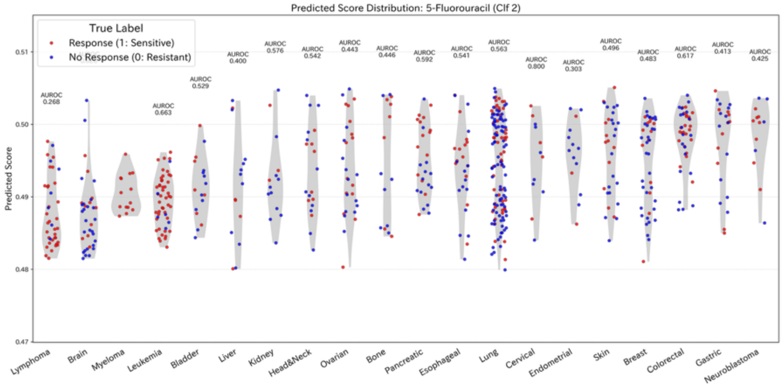


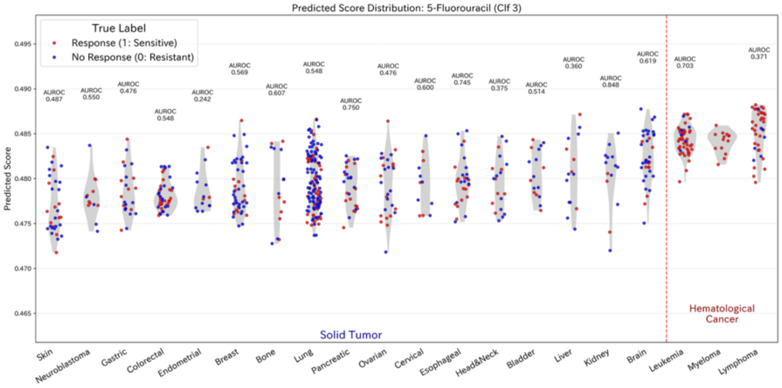


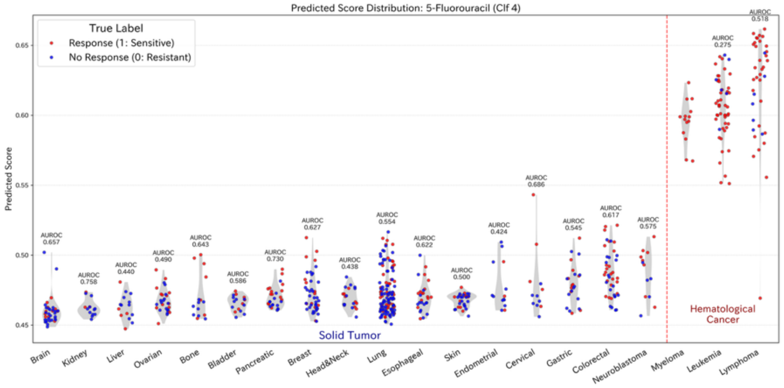


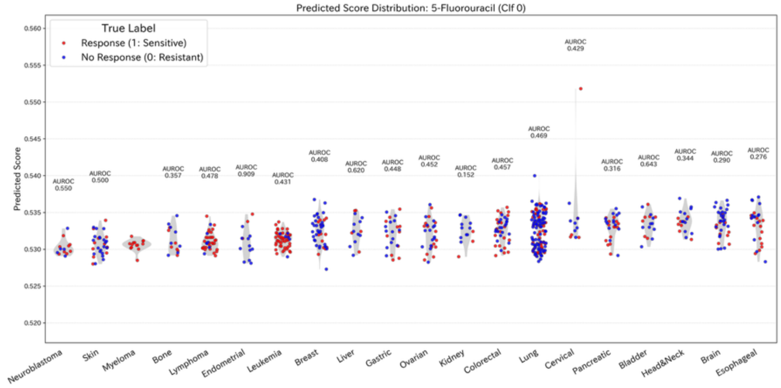


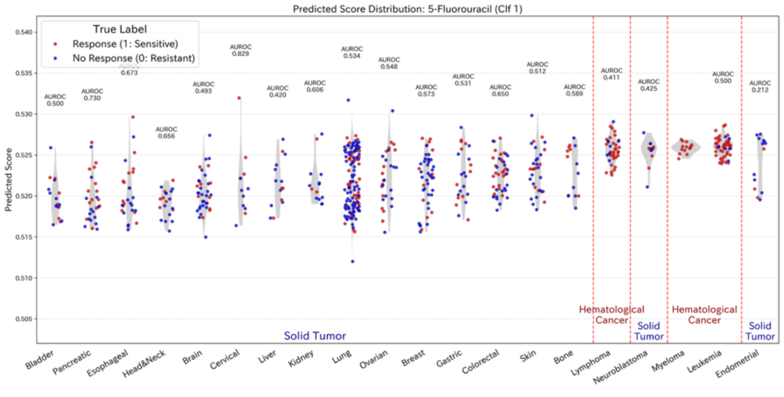


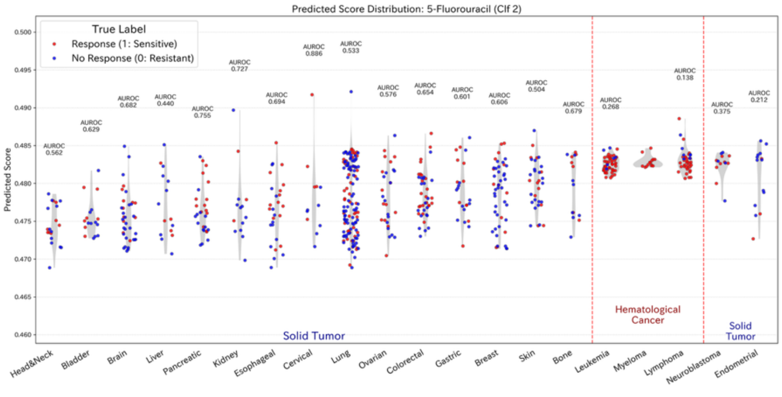

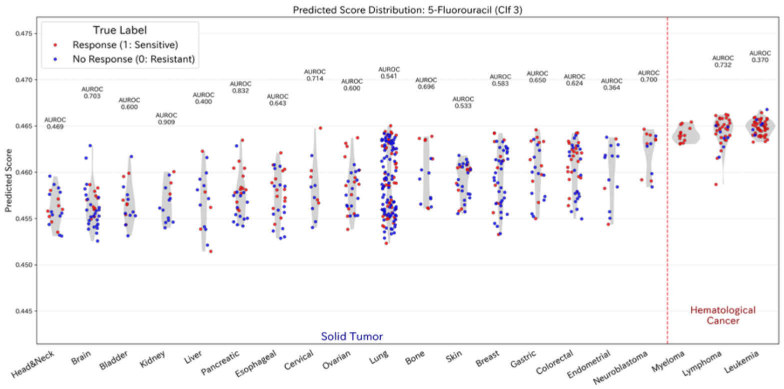
Predicted Score Distribution: 5-Fluorouracil (Clf 3)
True Label
Response (1: Sensitive)
No Response (0: Resistant)
Predicted Score
Solid Tumor
Hematological Cancer
Head&Neck Brain Bladder Kidney Liver Pancreatic Esophageal Cervical Ovarian Lung Bone Skin Breast Gastric Colorectal Endometrial Neuroblastoma Myeloma Lymphoma Leukemia

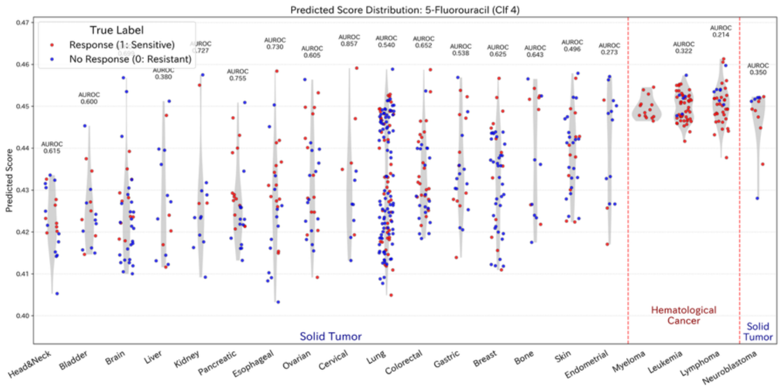
Predicted Score Distribution: 5-Fluorouracil (Clf 4)
True Label
Response (1: Sensitive)
No Response (0: Resistant)
Predicted Score
Solid Tumor
Hematological Cancer
Solid Tumor
Head&Neck Bladder Brain Liver Kidney Pancreatic Esophageal Ovarian Cervical Lung Colorectal Gastric Breast Bone Skin Endometrial Myeloma Leukemia Lymphoma Neuroblastoma

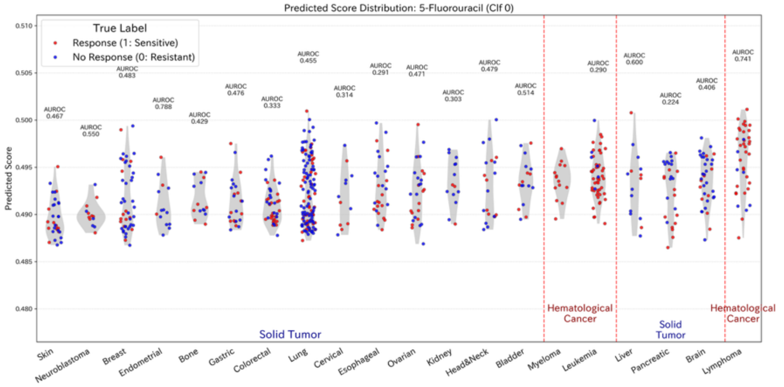
Predicted Score Distribution: 5-Fluorouracil (Clf 0)
True Label
Response (1: Sensitive)
No Response (0: Resistant)
Predicted Score
Solid Tumor
Hematological Cancer
Solid Tumor
Hematological Cancer
Skin Neuroblastoma Breast Endometrial Bone Gastric Colorectal Lung Cervical Esophageal Ovarian Kidney Head&Neck Bladder Myeloma Leukemia Liver Pancreatic Brain Lymphoma

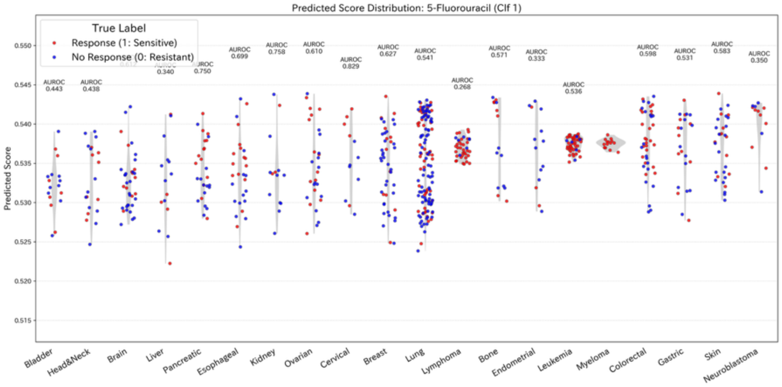
Predicted Score Distribution: 5-Fluorouracil (Clf 1)
True Label
Response (1: Sensitive)
No Response (0: Resistant)
Predicted Score
Bladder Head&Neck Brain Liver Pancreatic Esophageal Kidney Ovarian Cervical Breast Lung Lymphoma Bone Endometrial Leukemia Myeloma Colorectal Gastric Skin Neuroblastoma

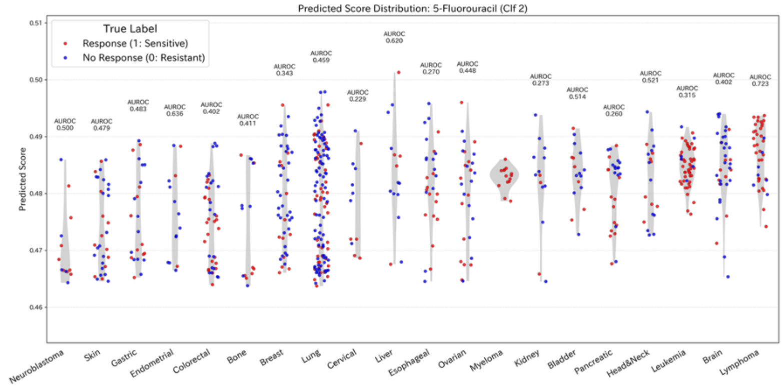
Predicted Score Distribution: 5-Fluorouracil (Clf 2)
True Label
Response (1: Sensitive)
No Response (0: Resistant)
Predicted Score
Neuroblastoma Skin Gastric Endometrial Colorectal Bone Breast Lung Cervical Liver Esophageal Ovarian Myeloma Kidney Bladder Pancreatic Head&Neck Leukemia Brain Lymphoma

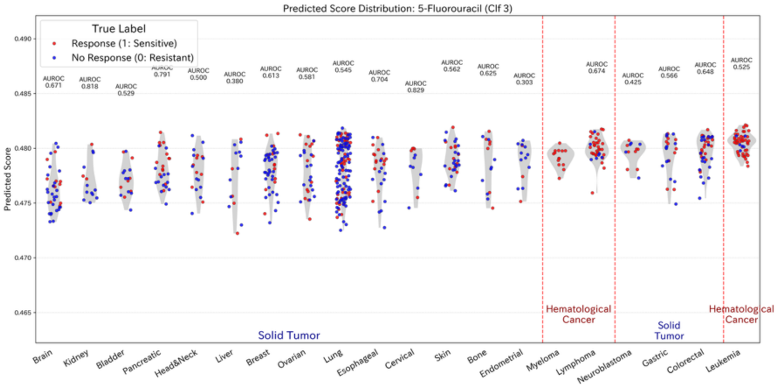
Predicted Score Distribution: 5-Fluorouracil (Clf 3)
True Label
Response (1: Sensitive)
No Response (0: Resistant)
Predicted Score
Solid Tumor
Hematological Cancer
Solid Tumor
Hematological Cancer
Brain Kidney Bladder Pancreatic Head&Neck Liver Breast Ovarian Lung Esophageal Cervical Skin Bone Endometrial Myeloma Lymphoma Neuroblastoma Gastric Colorectal Leukemia

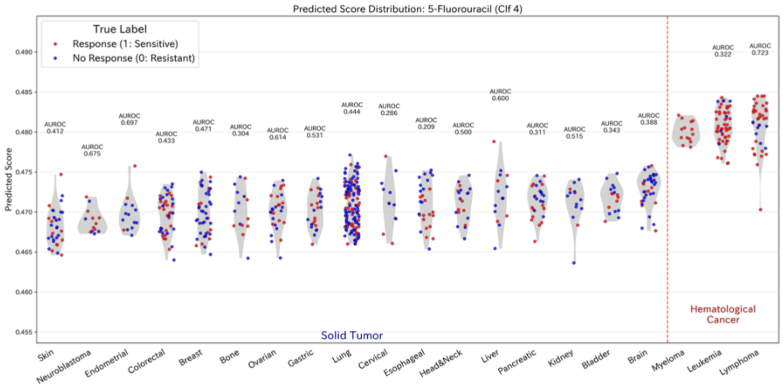
Predicted Score Distribution: 5-Fluorouracil (Clf 4)
True Label
Response (1: Sensitive)
No Response (0: Resistant)
Predicted Score
Solid Tumor
Hematological Cancer
Skin Neuroblastoma Endometrial Colorectal Breast Bone Ovarian Gastric Lung Cervical Esophageal Head&Neck Liver Pancreatic Kidney Bladder Brain Myeloma Leukemia Lymphoma

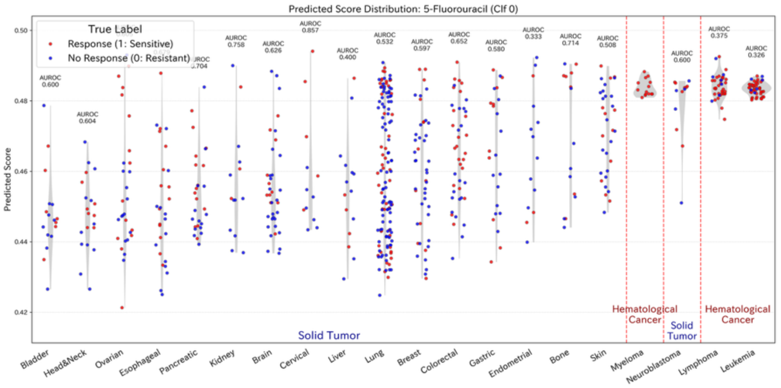
Predicted Score Distribution: 5-Fluorouracil (Clf 0)
True Label
Response (1: Sensitive)
No Response (0: Resistant)
Predicted Score
Solid Tumor
Hematological Cancer
Solid Tumor
Hematological Cancer
Bladder Head&Neck Ovarian Esophageal Pancreatic Kidney Brain Cervical Liver Lung Breast Colorectal Gastric Endometrial Bone Skin Myeloma Neuroblastoma Lymphoma Leukemia

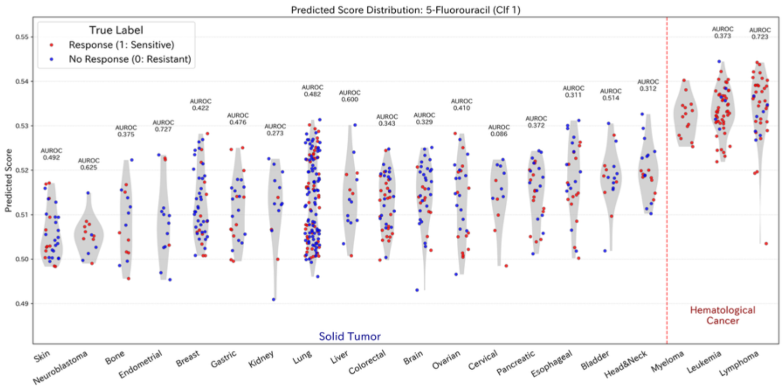
Predicted Score Distribution: 5-Fluorouracil (Clf 1)
True Label
Response (1: Sensitive)
No Response (0: Resistant)
Predicted Score
Solid Tumor
Hematological Cancer
Skin
Neuroblastoma
Bone
Endometrial
Breast
Gastric
Kidney
Lung
Liver
Colorectal
Brain
Ovarian
Cervical
Pancreatic
Esophageal
Bladder
Head&Neck
Myeloma
Leukemia
Lymphoma

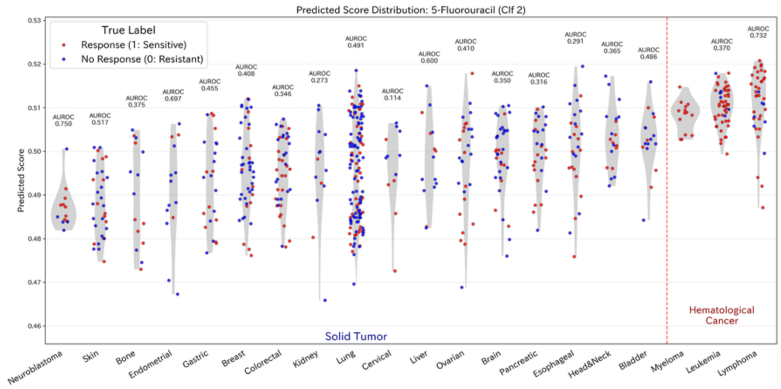
Predicted Score Distribution: 5-Fluorouracil (Clf 2)
True Label
Response (1: Sensitive)
No Response (0: Resistant)
Predicted Score
Solid Tumor
Hematological Cancer
Neuroblastoma
Skin
Bone
Endometrial
Gastric
Breast
Colorectal
Kidney
Lung
Cervical
Liver
Ovarian
Brain
Pancreatic
Esophageal
Head&Neck
Bladder
Myeloma
Leukemia
Lymphoma

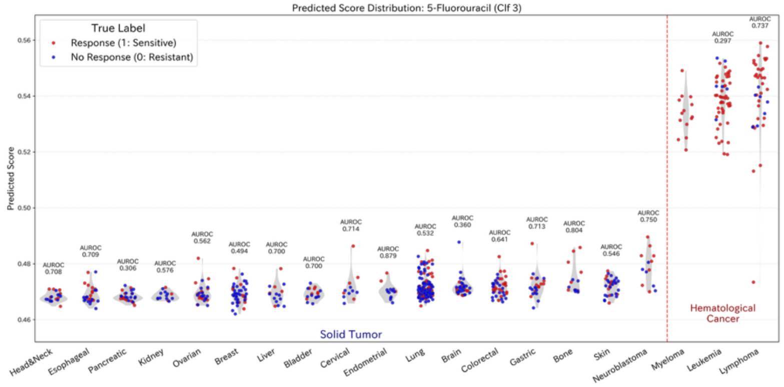
Predicted Score Distribution: 5-Fluorouracil (Clf 3)
True Label
Response (1: Sensitive)
No Response (0: Resistant)
Predicted Score
Solid Tumor
Hematological Cancer
Head&Neck
Esophageal
Pancreatic
Kidney
Ovarian
Breast
Liver
Bladder
Cervical
Endometrial
Lung
Brain
Colorectal
Gastric
Bone
Skin
Neuroblastoma
Myeloma
Leukemia
Lymphoma

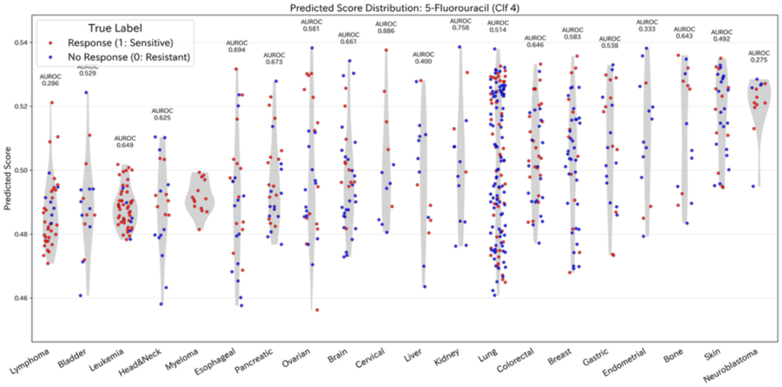
Predicted Score Distribution: 5-Fluorouracil (Clf 4)
True Label
Response (1: Sensitive)
No Response (0: Resistant)
Predicted Score
Lymphoma
Bladder
Leukemia
Head&Neck
Myeloma
Esophageal
Pancreatic
Ovarian
Brain
Cervical
Liver
Kidney
Lung
Colorectal
Breast
Gastric
Endometrial
Bone
Skin
Neuroblastoma

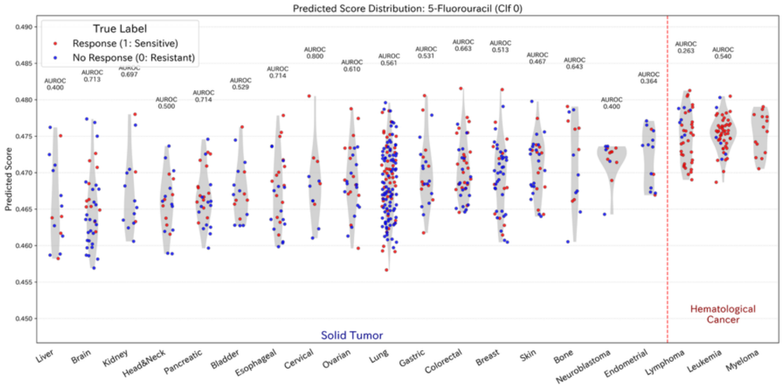
Predicted Score Distribution: 5-Fluorouracil (Clf 0)
True Label
Response (1: Sensitive)
No Response (0: Resistant)
Predicted Score
Solid Tumor
Hematological Cancer
Liver
Brain
Kidney
Head&Neck
Pancreatic
Bladder
Esophageal
Cervical
Ovarian
Lung
Gastric
Colorectal
Breast
Skin
Bone
Neuroblastoma
Endometrial
Lymphoma
Leukemia
Myeloma

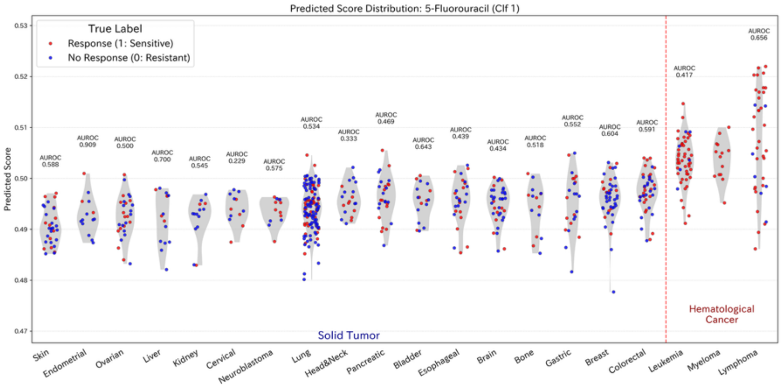
Predicted Score Distribution: 5-Fluorouracil (Clf 1)
True Label
Response (1: Sensitive)
No Response (0: Resistant)
Predicted Score
Solid Tumor
Hematological Cancer
Skin
Endometrial
Ovarian
Liver
Kidney
Cervical
Neuroblastoma
Lung
Head&Neck
Pancreatic
Bladder
Esophageal
Brain
Bone
Gastric
Breast
Colorectal
Leukemia
Myeloma
Lymphoma

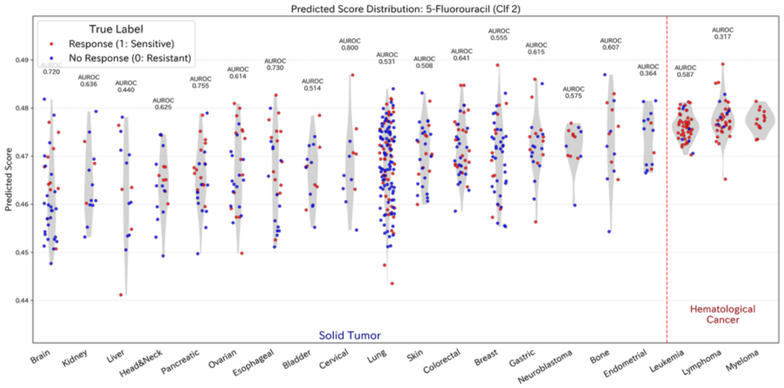
Predicted Score Distribution: 5-Fluorouracil (Clf 2)
True Label
Response (1: Sensitive)
No Response (0: Resistant)
Predicted Score
Solid Tumor
Hematological Cancer
Brain
Kidney
Liver
Head&Neck
Pancreatic
Ovarian
Esophageal
Bladder
Cervical
Lung
Skin
Colorectal
Breast
Gastric
Neuroblastoma
Bone
Endometrial
Leukemia
Lymphoma
Myeloma

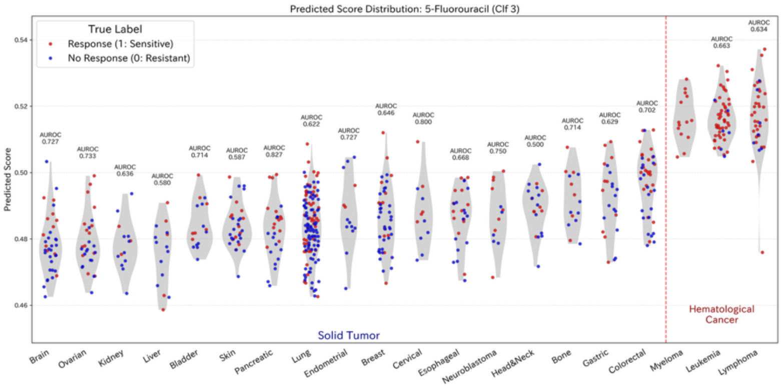
Predicted Score Distribution: 5-Fluorouracil (Clf 3)
True Label
Response (1: Sensitive)
No Response (0: Resistant)
Predicted Score
Solid Tumor
Hematological Cancer
Brain
Ovarian
Kidney
Liver
Bladder
Skin
Pancreatic
Lung
Endometrial
Breast
Cervical
Esophageal
Neuroblastoma
Head&Neck
Bone
Gastric
Colorectal
Myeloma
Leukemia
Lymphoma

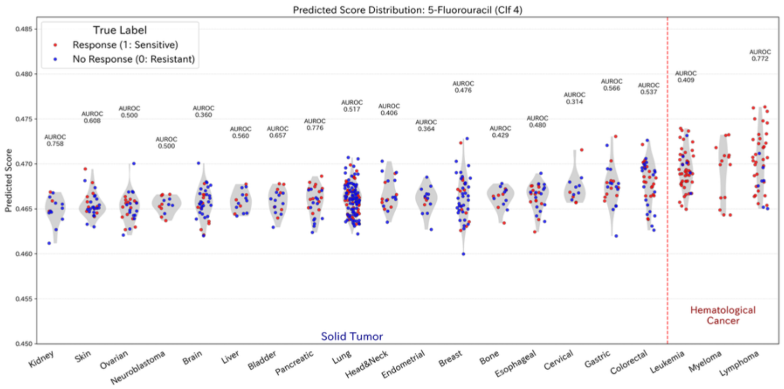

Predicted Score Distribution: 5-Fluorouracil (Clf 4)
True Label
Response (1: Sensitive)
No Response (0: Resistant)
Predicted Score
Solid Tumor
Hematological Cancer
Kidney
Skin
Ovarian
Neuroblastoma
Brain
Liver
Bladder
Pancreatic
Lung
Head&Neck
Endometrial
Breast
Bone
Esophageal
Cervical
Gastric
Colorectal
Leukemia
Myeloma
Lymphoma


[Cisplatin]

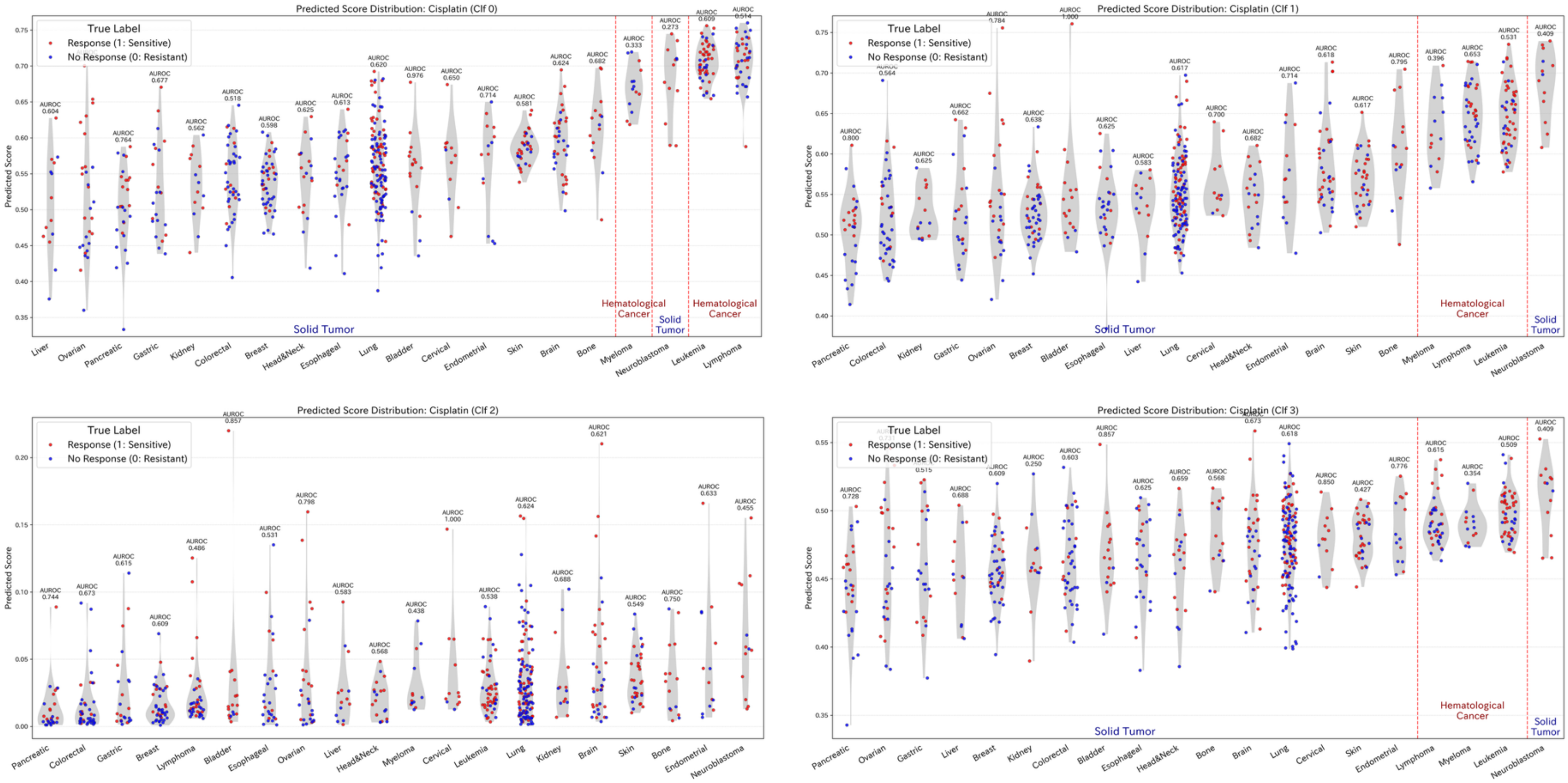

Predicted Score Distribution: Cisplatin (Clf 0)
Predicted Score Distribution: Cisplatin (Clf 1)
Predicted Score Distribution: Cisplatin (Clf 2)
Predicted Score Distribution: Cisplatin (Clf 3)
True Label
Response (1: Sensitive)
No Response (0: Resistant)
Predicted Score
Solid Tumor
Hematological Cancer

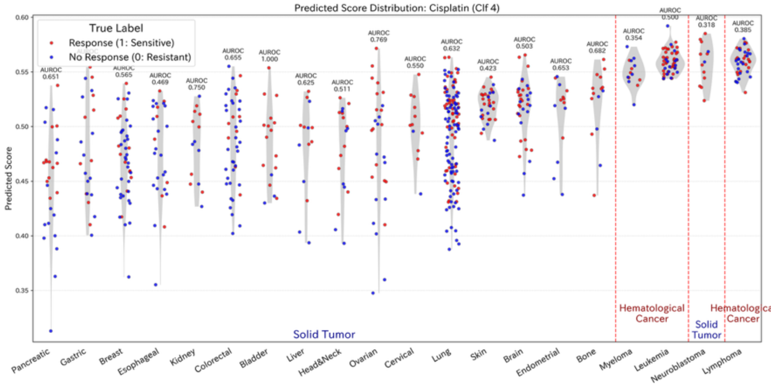
Predicted Score Distribution: Cisplatin (Clf 4)
True Label
Response (1: Sensitive)
No Response (0: Resistant)
Predicted Score
Solid Tumor
Hematological Cancer
Solid Tumor
Hematological Cancer
Pancreatic
Gastric
Breast
Esophageal
Kidney
Colorectal
Bladder
Liver
Head&Neck
Ovarian
Cervical
Lung
Skin
Brain
Endometrial
Bone
Myeloma
Leukemia
Neuroblastoma
Lymphoma

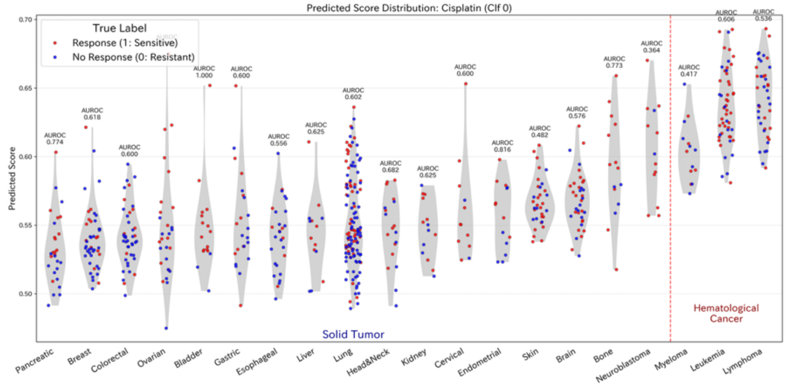
Predicted Score Distribution: Cisplatin (Clf 0)
True Label
Response (1: Sensitive)
No Response (0: Resistant)
Predicted Score
Solid Tumor
Hematological Cancer
Pancreatic
Breast
Colorectal
Ovarian
Bladder
Gastric
Esophageal
Liver
Lung
Head&Neck
Kidney
Cervical
Endometrial
Skin
Brain
Bone
Neuroblastoma
Myeloma
Leukemia
Lymphoma

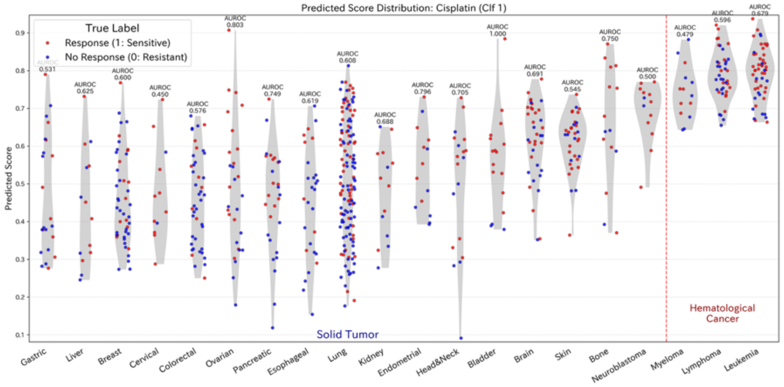
Predicted Score Distribution: Cisplatin (Clf 1)
True Label
Response (1: Sensitive)
No Response (0: Resistant)
Predicted Score
Solid Tumor
Hematological Cancer
Gastric
Liver
Breast
Cervical
Colorectal
Ovarian
Pancreatic
Esophageal
Lung
Kidney
Endometrial
Head&Neck
Bladder
Brain
Skin
Bone
Neuroblastoma
Myeloma
Lymphoma
Leukemia

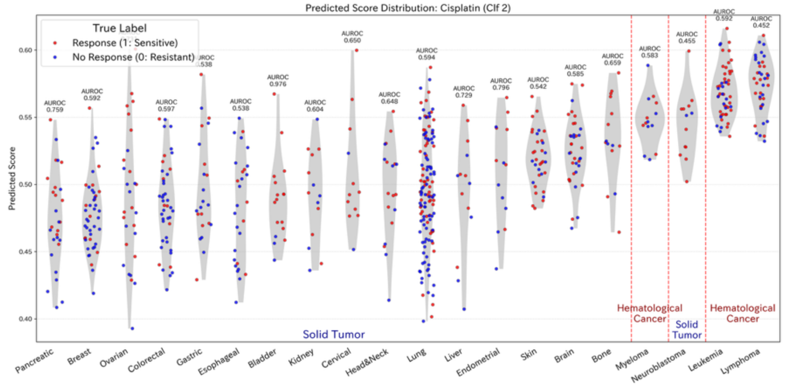
Predicted Score Distribution: Cisplatin (Clf 2)
True Label
Response (1: Sensitive)
No Response (0: Resistant)
Predicted Score
Solid Tumor
Hematological Cancer
Solid Tumor
Hematological Cancer
Pancreatic
Breast
Ovarian
Colorectal
Gastric
Esophageal
Bladder
Kidney
Cervical
Head&Neck
Lung
Liver
Endometrial
Skin
Brain
Bone
Myeloma
Neuroblastoma
Leukemia
Lymphoma

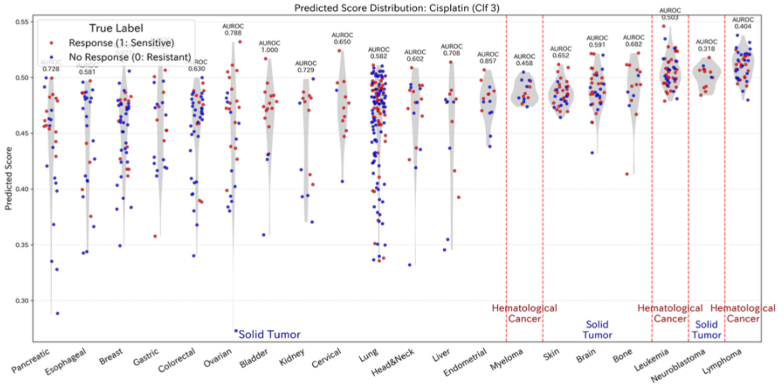
Predicted Score Distribution: Cisplatin (Clf 3)
True Label
Response (1: Sensitive)
No Response (0: Resistant)
Predicted Score
Solid Tumor
Hematological Cancer
Solid Tumor
Hematological Cancer
Solid Tumor
Hematological Cancer
Pancreatic
Esophageal
Breast
Gastric
Colorectal
Ovarian
Bladder
Kidney
Cervical
Lung
Head&Neck
Liver
Endometrial
Myeloma
Skin
Brain
Bone
Leukemia
Neuroblastoma
Lymphoma

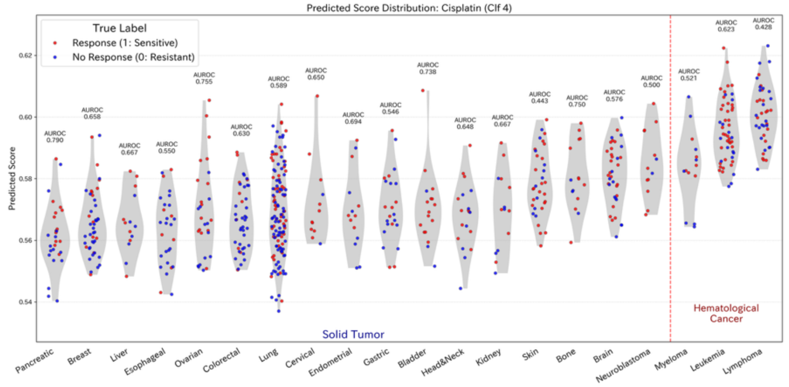
Predicted Score Distribution: Cisplatin (Clf 4)
True Label
Response (1: Sensitive)
No Response (0: Resistant)
Predicted Score
Solid Tumor
Hematological Cancer
Pancreatic
Breast
Liver
Esophageal
Ovarian
Colorectal
Lung
Cervical
Endometrial
Gastric
Bladder
Head&Neck
Kidney
Skin
Bone
Brain
Neuroblastoma
Myeloma
Leukemia
Lymphoma

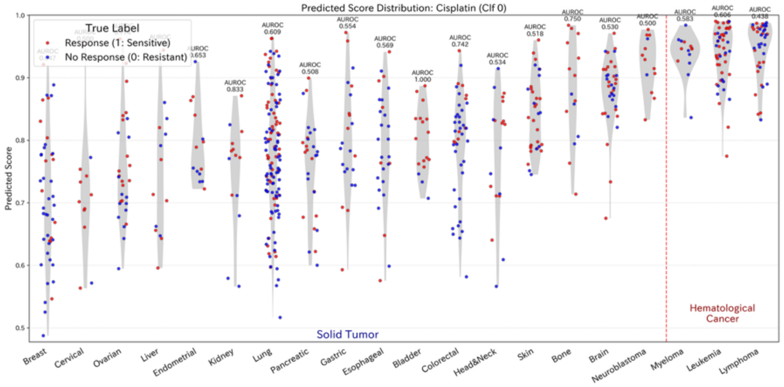
Predicted Score Distribution: Cisplatin (Clf 0)
True Label
Response (1: Sensitive)
No Response (0: Resistant)
Predicted Score
Solid Tumor
Hematological Cancer
Breast
Cervical
Ovarian
Liver
Endometrial
Kidney
Lung
Pancreatic
Gastric
Esophageal
Bladder
Colorectal
Head&Neck
Skin
Bone
Brain
Neuroblastoma
Myeloma
Leukemia
Lymphoma

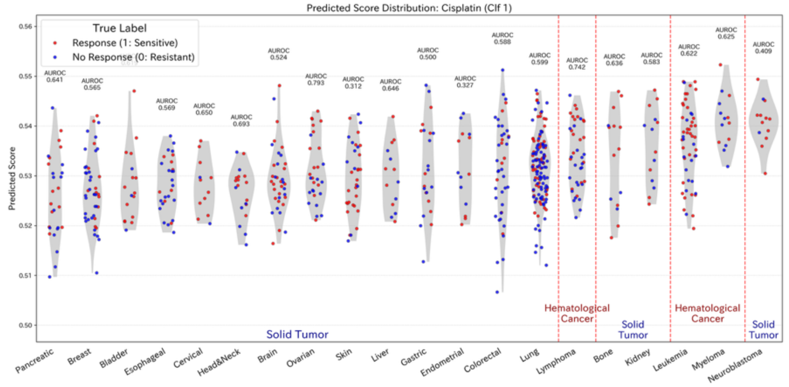
Predicted Score Distribution: Cisplatin (Clf 1)
True Label
Response (1: Sensitive)
No Response (0: Resistant)
Predicted Score
Solid Tumor
Hematological Cancer
Solid Tumor
Hematological Cancer
Solid Tumor
Pancreatic
Breast
Bladder
Esophageal
Cervical
Head&Neck
Brain
Ovarian
Skin
Liver
Gastric
Endometrial
Colorectal
Lung
Lymphoma
Bone
Kidney
Leukemia
Myeloma
Neuroblastoma

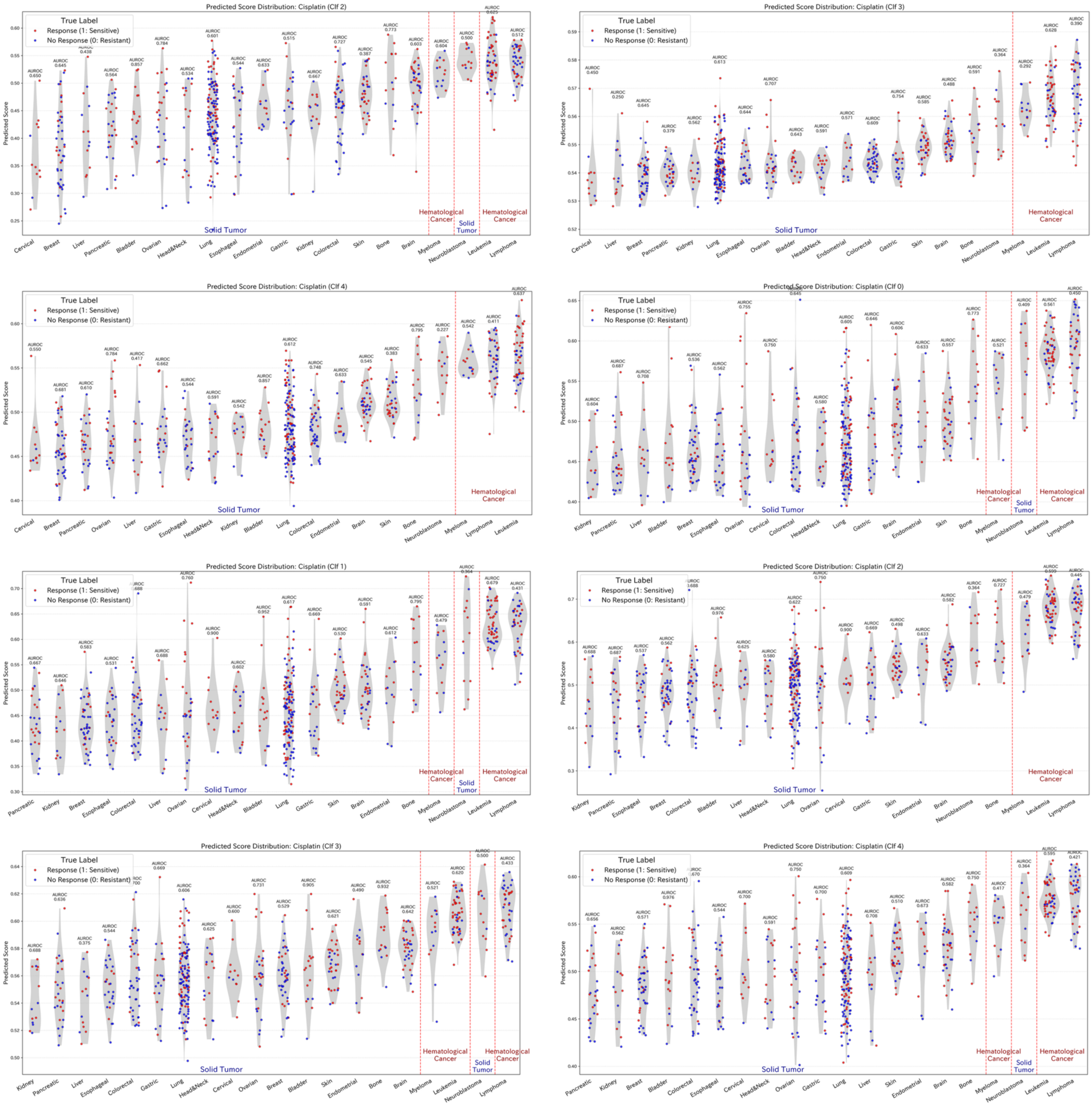
Predicted Score Distribution: Cisplatin (Clf 2)
Predicted Score Distribution: Cisplatin (Clf 3)
Predicted Score Distribution: Cisplatin (Clf 4)
Predicted Score Distribution: Cisplatin (Clf 0)
Predicted Score Distribution: Cisplatin (Clf 1)
Predicted Score Distribution: Cisplatin (Clf 2)
Predicted Score Distribution: Cisplatin (Clf 3)
Predicted Score Distribution: Cisplatin (Clf 4)
True Label
Response (1: Sensitive)
No Response (0: Resistant)
Predicted Score
Solid Tumor
Hematological Cancer

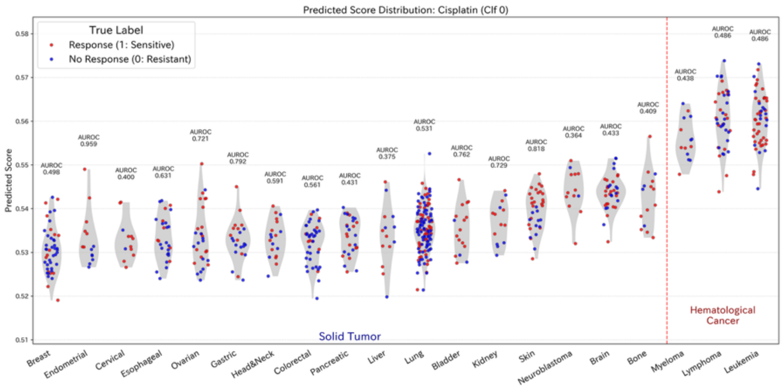
Predicted Score Distribution: Cisplatin (Clf 0)
True Label
Response (1: Sensitive)
No Response (0: Resistant)
Predicted Score
Solid Tumor
Hematological Cancer
Breast
Endometrial
Cervical
Esophageal
Ovarian
Gastric
Head&Neck
Colorectal
Pancreatic
Liver
Lung
Bladder
Kidney
Skin
Neuroblastoma
Brain
Bone
Myeloma
Lymphoma
Leukemia

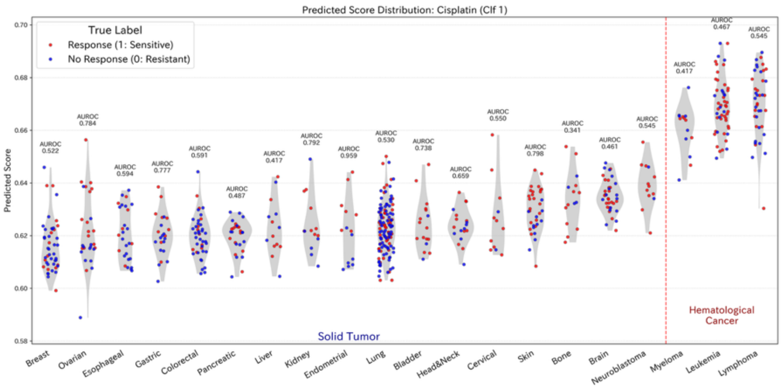
Predicted Score Distribution: Cisplatin (Clf 1)
True Label
Response (1: Sensitive)
No Response (0: Resistant)
Predicted Score
Solid Tumor
Hematological Cancer
Breast
Ovarian
Esophageal
Gastric
Colorectal
Pancreatic
Liver
Kidney
Endometrial
Lung
Bladder
Head&Neck
Cervical
Skin
Bone
Brain
Neuroblastoma
Myeloma
Leukemia
Lymphoma

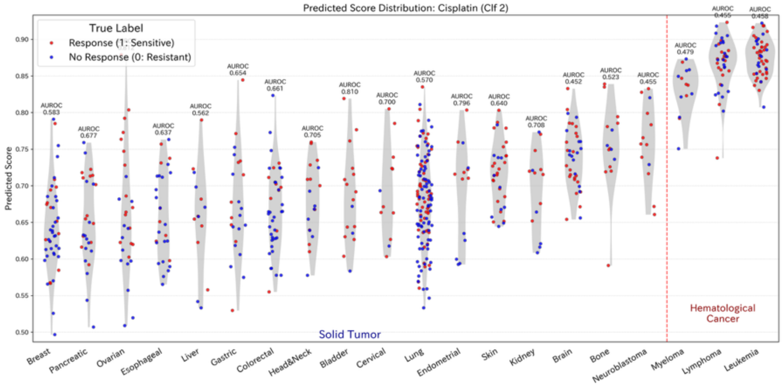
Predicted Score Distribution: Cisplatin (Clf 2)
True Label
Response (1: Sensitive)
No Response (0: Resistant)
Predicted Score
Solid Tumor
Hematological Cancer
Breast
Pancreatic
Ovarian
Esophageal
Liver
Gastric
Colorectal
Head&Neck
Bladder
Cervical
Lung
Endometrial
Skin
Kidney
Brain
Bone
Neuroblastoma
Myeloma
Lymphoma
Leukemia

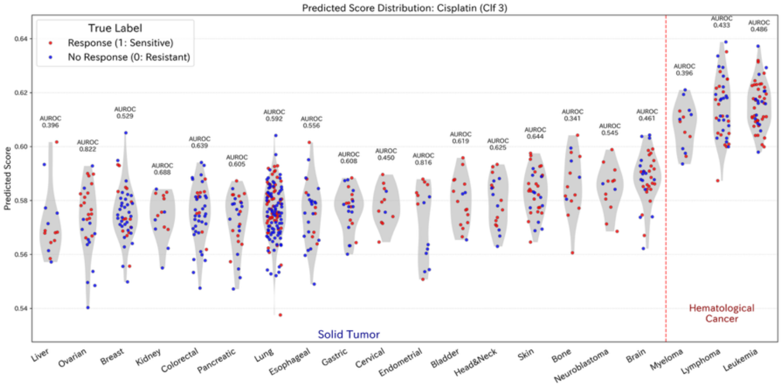
Predicted Score Distribution: Cisplatin (Clf 3)
True Label
Response (1: Sensitive)
No Response (0: Resistant)
Predicted Score
Solid Tumor
Hematological Cancer
Liver
Ovarian
Breast
Kidney
Colorectal
Pancreatic
Lung
Esophageal
Gastric
Cervical
Endometrial
Bladder
Head&Neck
Skin
Bone
Neuroblastoma
Brain
Myeloma
Lymphoma
Leukemia

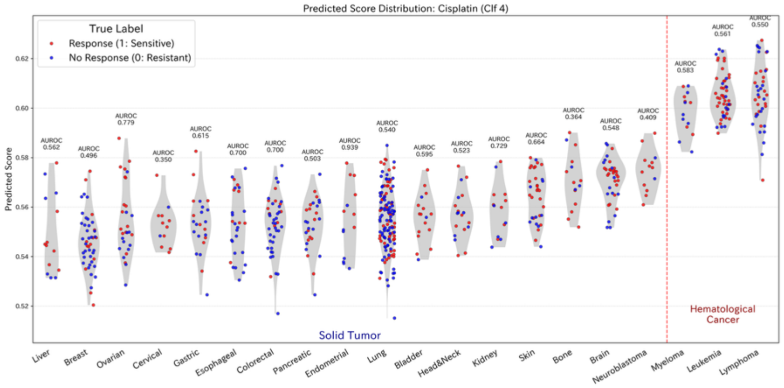
Predicted Score Distribution: Cisplatin (Clf 4)
True Label
Response (1: Sensitive)
No Response (0: Resistant)
Predicted Score
Solid Tumor
Hematological Cancer
Liver
Breast
Ovarian
Cervical
Gastric
Esophageal
Colorectal
Pancreatic
Endometrial
Lung
Bladder
Head&Neck
Kidney
Skin
Bone
Brain
Neuroblastoma
Myeloma
Leukemia
Lymphoma

[Gemcitabine]

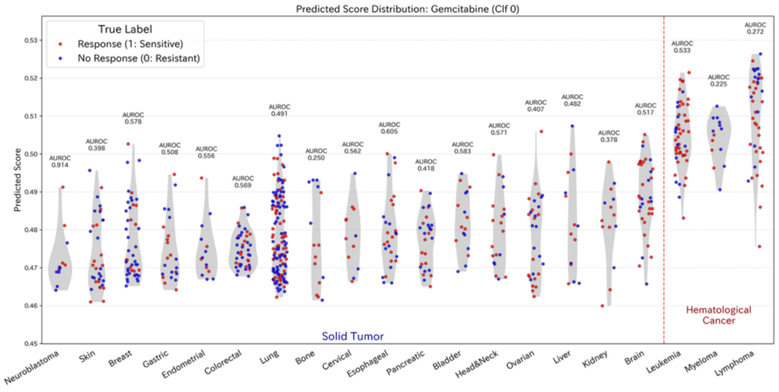


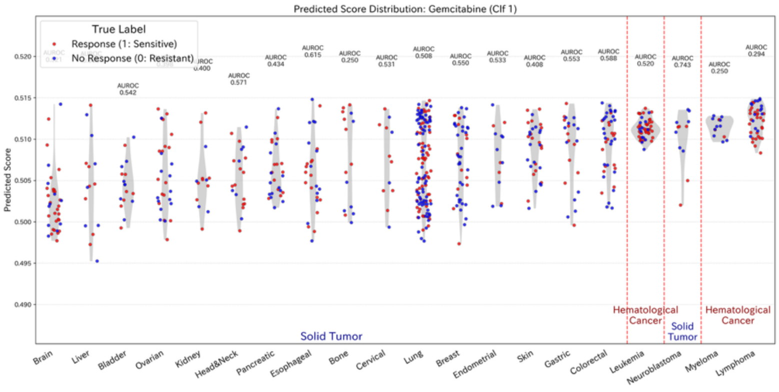


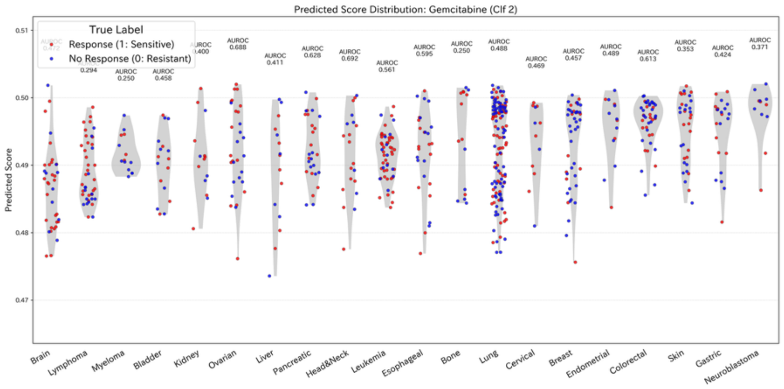


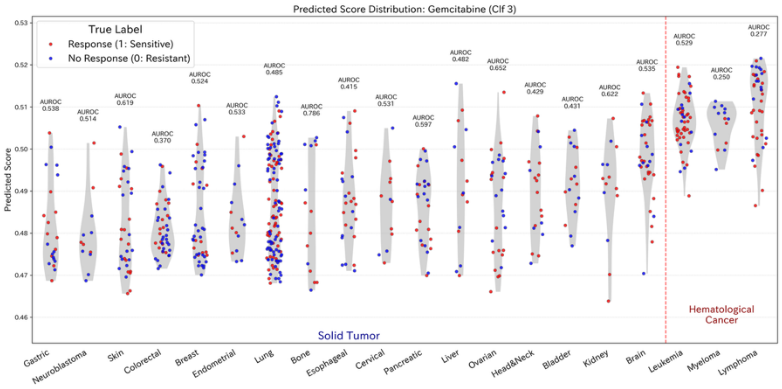


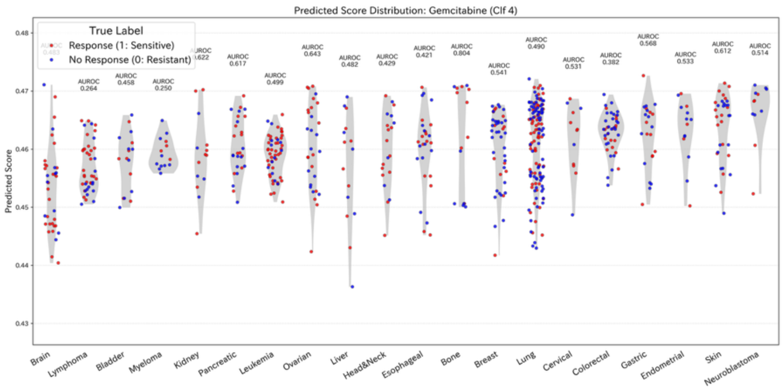


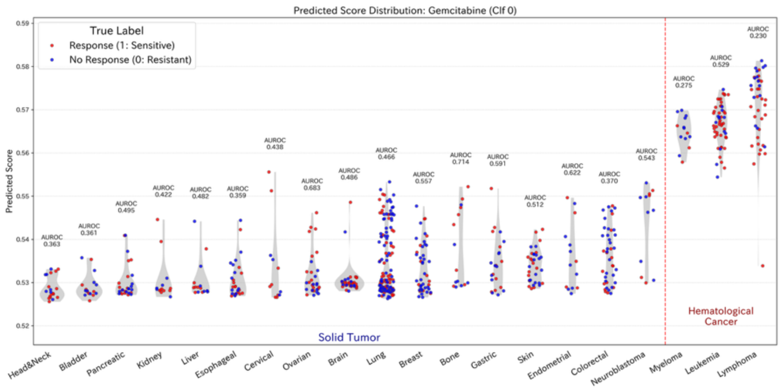


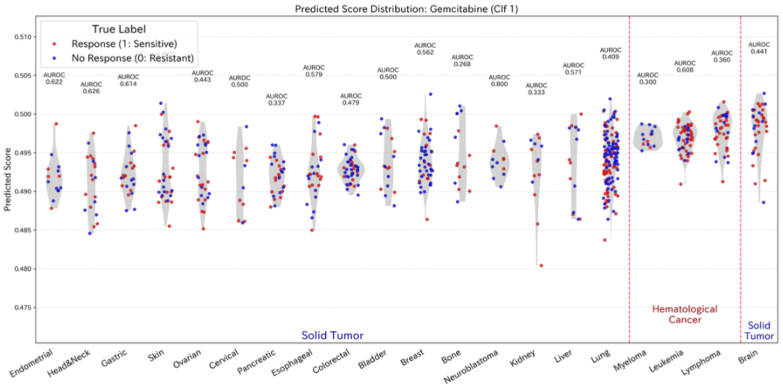


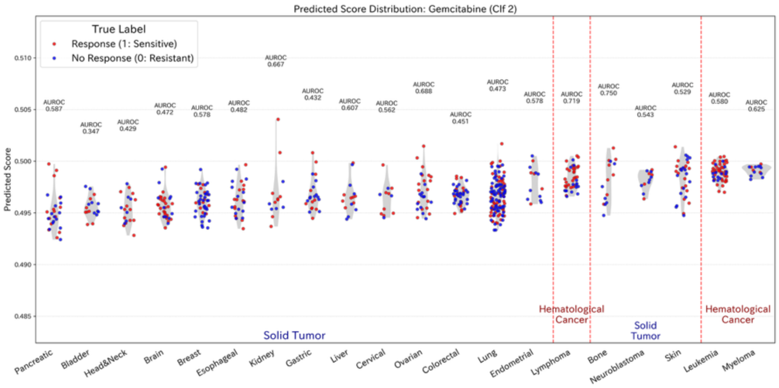

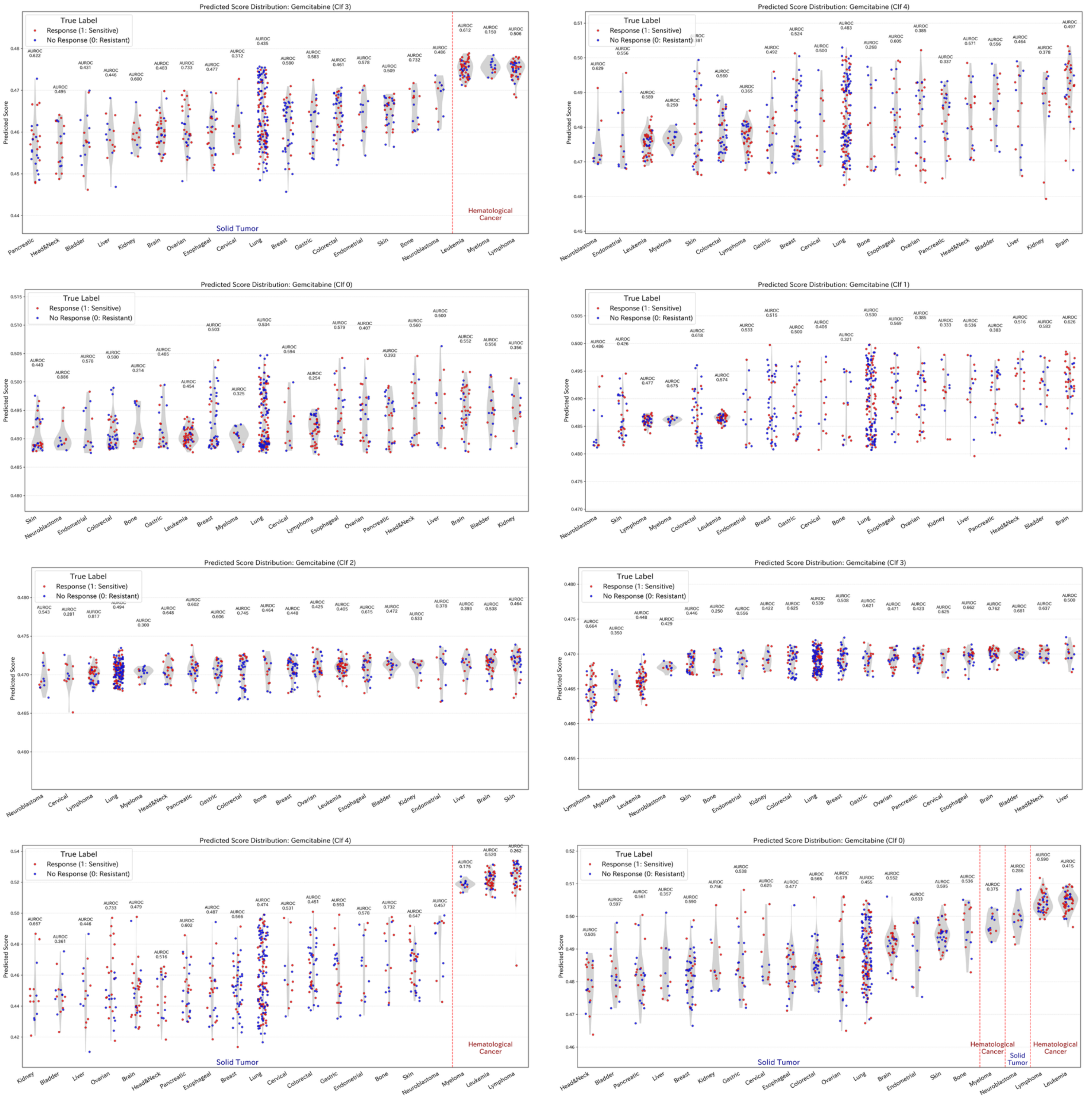
Predicted Score Distribution: Gemcitabine (Clf 3)
Predicted Score Distribution: Gemcitabine (Clf 4)
Predicted Score Distribution: Gemcitabine (Clf 0)
Predicted Score Distribution: Gemcitabine (Clf 1)
Predicted Score Distribution: Gemcitabine (Clf 2)
Predicted Score Distribution: Gemcitabine (Clf 3)
Predicted Score Distribution: Gemcitabine (Clf 4)
Predicted Score Distribution: Gemcitabine (Clf 0)
True Label
Response (1: Sensitive)
No Response (0: Resistant)
Predicted Score
Solid Tumor
Hematological Cancer

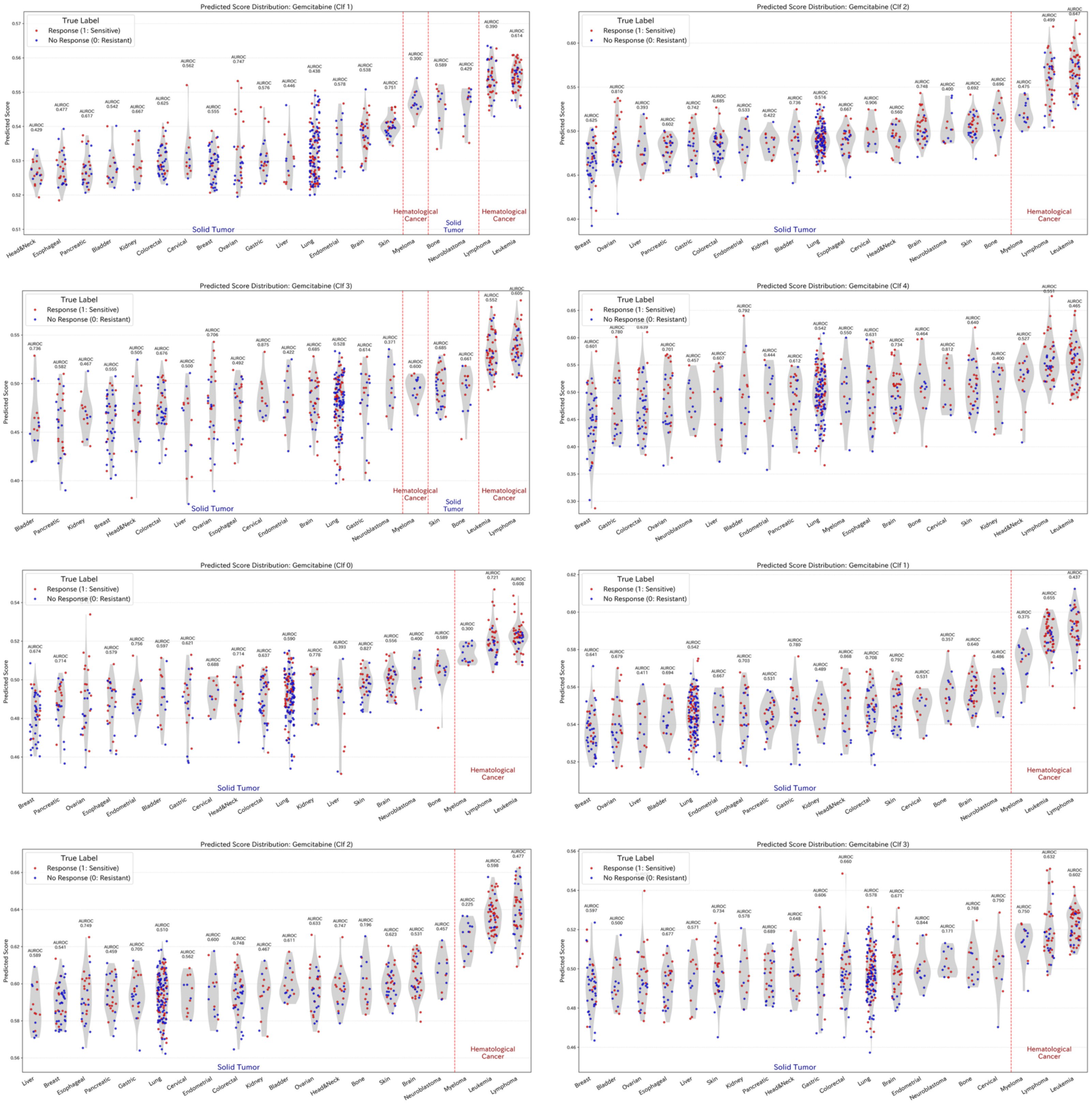
Predicted Score Distribution: Gemcitabine (Clf 1)
True Label
Response (1: Sensitive)
No Response (0: Resistant)
Predicted Score
Solid Tumor
Hematological Cancer
Solid Tumor
Hematological Cancer
Head&Neck
Esophageal
Pancreatic
Bladder
Kidney
Colorectal
Cervical
Breast
Ovarian
Gastric
Liver
Lung
Endometrial
Brain
Skin
Myeloma
Bone
Neuroblastoma
Lymphoma
Leukemia
Predicted Score Distribution: Gemcitabine (Clf 2)
True Label
Response (1: Sensitive)
No Response (0: Resistant)
Predicted Score
Solid Tumor
Hematological Cancer
Breast
Ovarian
Liver
Pancreatic
Gastric
Colorectal
Endometrial
Kidney
Bladder
Lung
Esophageal
Cervical
Head&Neck
Brain
Neuroblastoma
Skin
Bone
Myeloma
Lymphoma
Leukemia
Predicted Score Distribution: Gemcitabine (Clf 3)
True Label
Response (1: Sensitive)
No Response (0: Resistant)
Predicted Score
Solid Tumor
Hematological Cancer
Solid Tumor
Hematological Cancer
Bladder
Pancreatic
Kidney
Breast
Head&Neck
Colorectal
Liver
Ovarian
Esophageal
Cervical
Endometrial
Brain
Lung
Gastric
Neuroblastoma
Myeloma
Skin
Bone
Leukemia
Lymphoma
Predicted Score Distribution: Gemcitabine (Clf 4)
True Label
Response (1: Sensitive)
No Response (0: Resistant)
Predicted Score
Breast
Gastric
Colorectal
Ovarian
Neuroblastoma
Liver
Bladder
Endometrial
Pancreatic
Lung
Myeloma
Esophageal
Brain
Bone
Cervical
Skin
Kidney
Head&Neck
Lymphoma
Leukemia
Predicted Score Distribution: Gemcitabine (Clf 0)
True Label
Response (1: Sensitive)
No Response (0: Resistant)
Predicted Score
Solid Tumor
Hematological Cancer
Breast
Pancreatic
Ovarian
Esophageal
Endometrial
Bladder
Gastric
Cervical
Head&Neck
Colorectal
Lung
Kidney
Liver
Skin
Brain
Neuroblastoma
Bone
Myeloma
Lymphoma
Leukemia
Predicted Score Distribution: Gemcitabine (Clf 1)
True Label
Response (1: Sensitive)
No Response (0: Resistant)
Predicted Score
Solid Tumor
Hematological Cancer
Breast
Ovarian
Liver
Bladder
Lung
Endometrial
Esophageal
Pancreatic
Gastric
Kidney
Head&Neck
Colorectal
Skin
Cervical
Bone
Brain
Neuroblastoma
Myeloma
Leukemia
Lymphoma
Predicted Score Distribution: Gemcitabine (Clf 2)
True Label
Response (1: Sensitive)
No Response (0: Resistant)
Predicted Score
Solid Tumor
Hematological Cancer
Liver
Breast
Esophageal
Pancreatic
Gastric
Lung
Cervical
Endometrial
Colorectal
Kidney
Bladder
Ovarian
Head&Neck
Bone
Skin
Brain
Neuroblastoma
Myeloma
Leukemia
Lymphoma
Predicted Score Distribution: Gemcitabine (Clf 3)
True Label
Response (1: Sensitive)
No Response (0: Resistant)
Predicted Score
Solid Tumor
Hematological Cancer
Breast
Bladder
Ovarian
Esophageal
Liver
Skin
Kidney
Pancreatic
Head&Neck
Gastric
Colorectal
Lung
Brain
Endometrial
Neuroblastoma
Bone
Cervical
Myeloma
Lymphoma
Leukemia

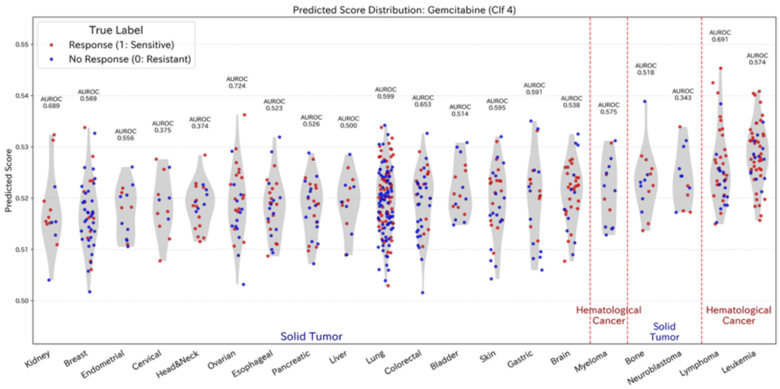
Predicted Score Distribution: Gemcitabine (Clf 4)
True Label
Response (1: Sensitive)
No Response (0: Resistant)
Predicted Score
Solid Tumor
Hematological Cancer
Solid Tumor
Hematological Cancer
Kidney
Breast
Endometrial
Cervical
Head&Neck
Ovarian
Esophageal
Pancreatic
Liver
Lung
Colorectal
Bladder
Skin
Gastric
Brain
Myeloma
Bone
Neuroblastoma
Lymphoma
Leukemia

[Sorafenib]

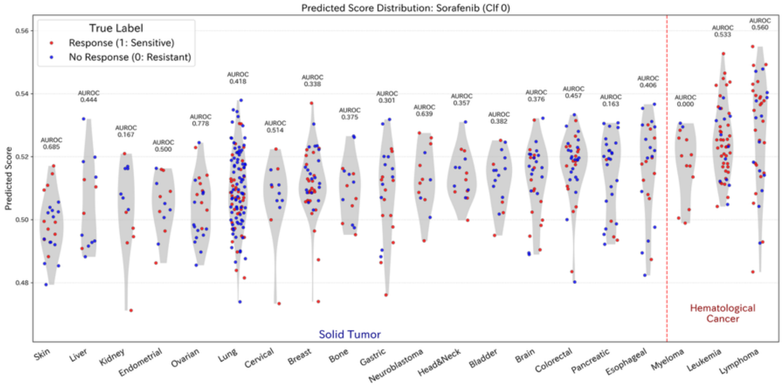
Predicted Score Distribution: Sorafenib (Clf 0)
True Label
Response (1: Sensitive)
No Response (0: Resistant)
Predicted Score
Solid Tumor
Hematological Cancer
Skin
Liver
Kidney
Endometrial
Ovarian
Lung
Cervical
Breast
Bone
Gastric
Neuroblastoma
Head&Neck
Bladder
Brain
Colorectal
Pancreatic
Esophageal
Myeloma
Leukemia
Lymphoma

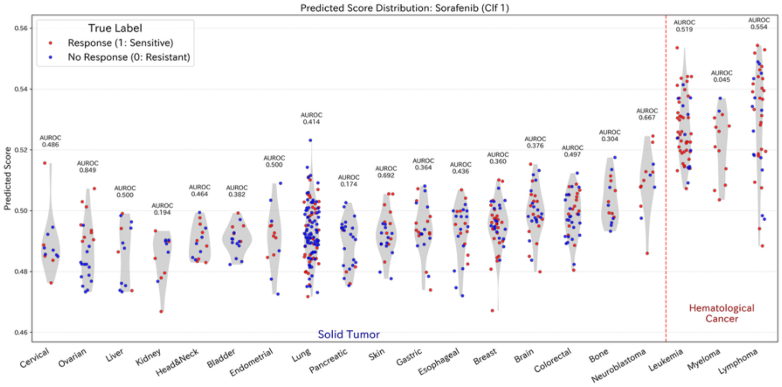
Predicted Score Distribution: Sorafenib (Clf 1)
True Label
Response (1: Sensitive)
No Response (0: Resistant)
Predicted Score
Solid Tumor
Hematological Cancer
Cervical
Ovarian
Liver
Kidney
Head&Neck
Bladder
Endometrial
Lung
Pancreatic
Skin
Gastric
Esophageal
Breast
Brain
Colorectal
Bone
Neuroblastoma
Leukemia
Myeloma
Lymphoma

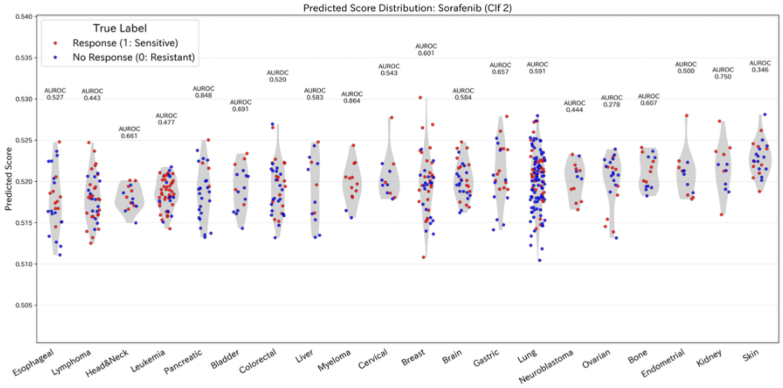
Predicted Score Distribution: Sorafenib (Clf 2)
True Label
Response (1: Sensitive)
No Response (0: Resistant)
Predicted Score
Esophageal
Lymphoma
Head&Neck
Leukemia
Pancreatic
Bladder
Colorectal
Liver
Myeloma
Cervical
Breast
Brain
Gastric
Lung
Neuroblastoma
Ovarian
Bone
Endometrial
Kidney
Skin

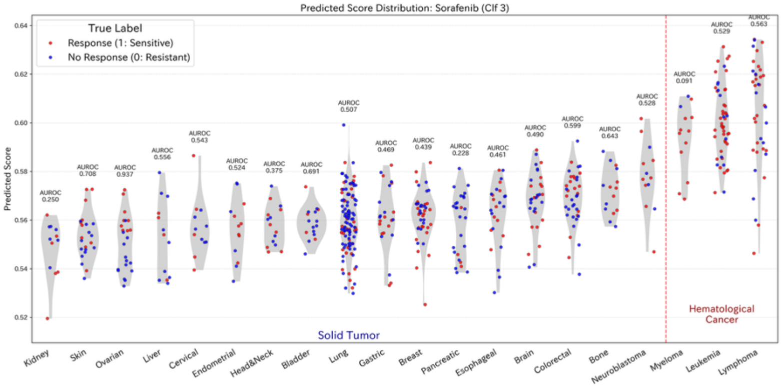
Predicted Score Distribution: Sorafenib (Clf 3)
True Label
Response (1: Sensitive)
No Response (0: Resistant)
Predicted Score
Solid Tumor
Hematological Cancer
Kidney
Skin
Ovarian
Liver
Cervical
Endometrial
Head&Neck
Bladder
Lung
Gastric
Breast
Pancreatic
Esophageal
Brain
Colorectal
Bone
Neuroblastoma
Myeloma
Leukemia
Lymphoma

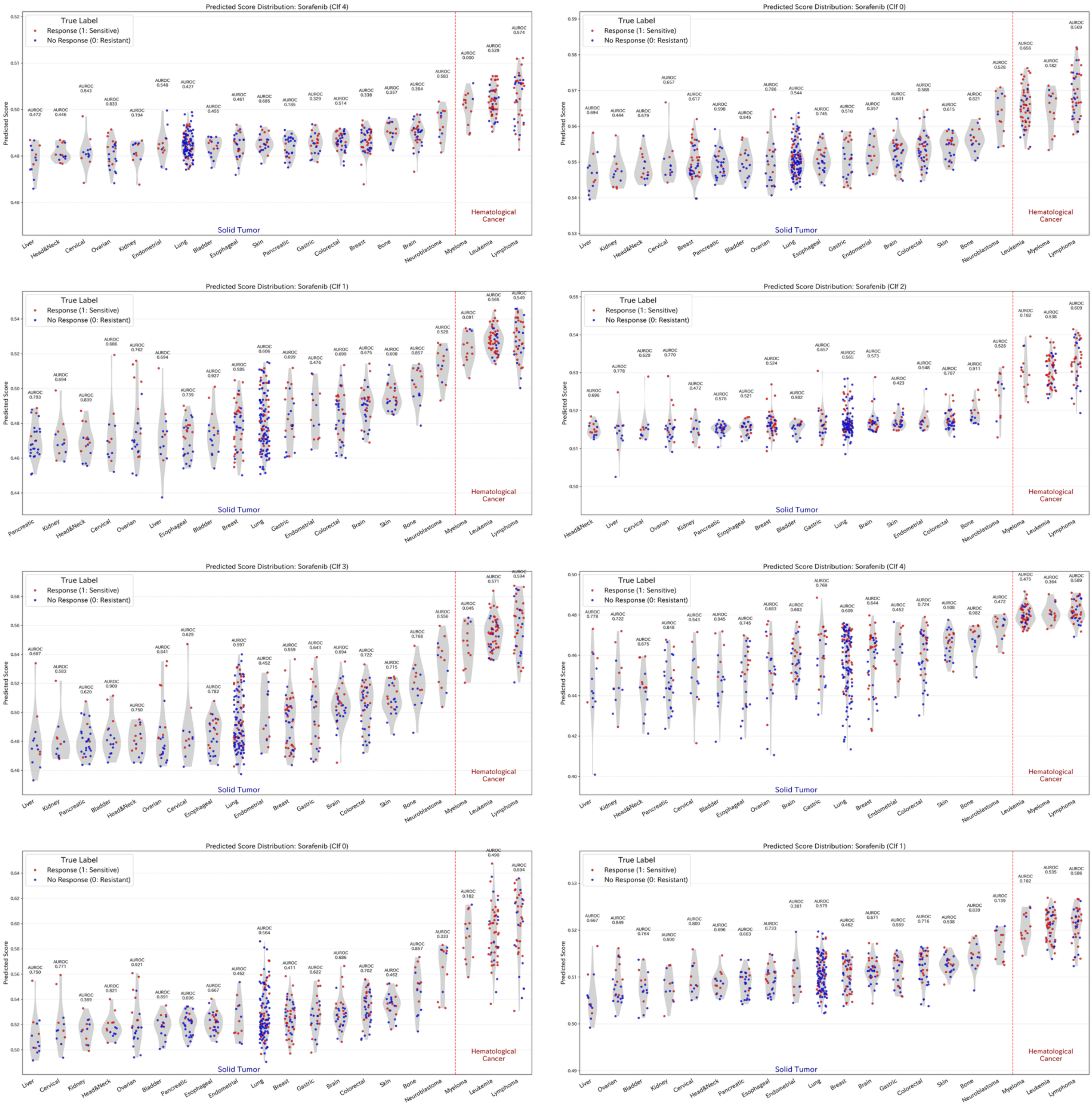
Predicted Score Distribution: Sorafenib (Clf 4)
Predicted Score Distribution: Sorafenib (Clf 0)
Predicted Score Distribution: Sorafenib (Clf 1)
Predicted Score Distribution: Sorafenib (Clf 2)
Predicted Score Distribution: Sorafenib (Clf 3)
Predicted Score Distribution: Sorafenib (Clf 4)
Predicted Score Distribution: Sorafenib (Clf 0)
Predicted Score Distribution: Sorafenib (Clf 1)
True Label
Response (1: Sensitive)
No Response (0: Resistant)
Predicted Score
Solid Tumor
Hematological Cancer

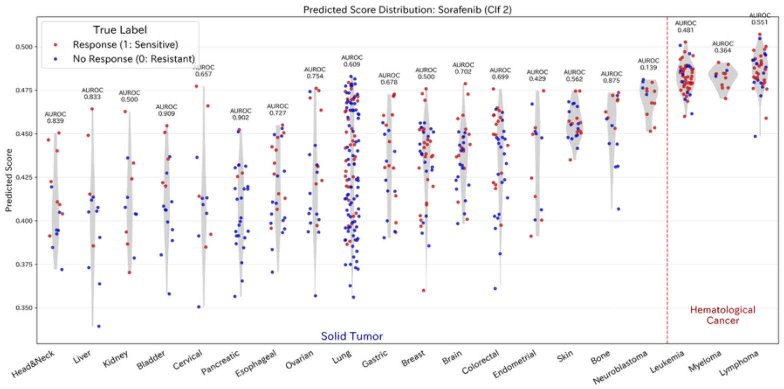
Predicted Score Distribution: Sorafenib (Clf 2)
True Label
Response (1: Sensitive)
No Response (0: Resistant)
Predicted Score
Solid Tumor
Hematological Cancer
Head&Neck
Liver
Kidney
Bladder
Cervical
Pancreatic
Esophageal
Ovarian
Lung
Gastric
Breast
Brain
Colorectal
Endometrial
Skin
Bone
Neuroblastoma
Leukemia
Myeloma
Lymphoma

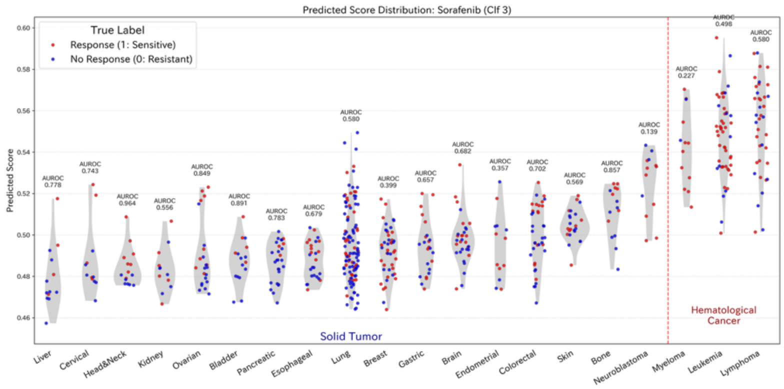
Predicted Score Distribution: Sorafenib (Clf 3)
True Label
Response (1: Sensitive)
No Response (0: Resistant)
Predicted Score
Solid Tumor
Hematological Cancer
Liver
Cervical
Head&Neck
Kidney
Ovarian
Bladder
Pancreatic
Esophageal
Lung
Breast
Gastric
Brain
Endometrial
Colorectal
Skin
Bone
Neuroblastoma
Myeloma
Leukemia
Lymphoma

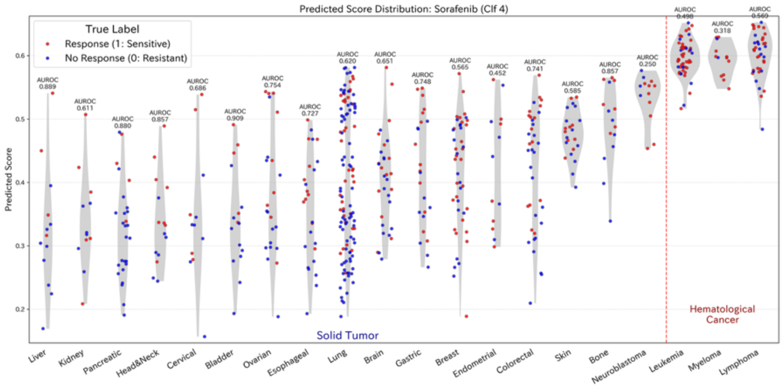
Predicted Score Distribution: Sorafenib (Clf 4)
True Label
Response (1: Sensitive)
No Response (0: Resistant)
Predicted Score
Solid Tumor
Hematological Cancer
Liver
Kidney
Pancreatic
Head&Neck
Cervical
Bladder
Ovarian
Esophageal
Lung
Brain
Gastric
Breast
Endometrial
Colorectal
Skin
Bone
Neuroblastoma
Leukemia
Myeloma
Lymphoma

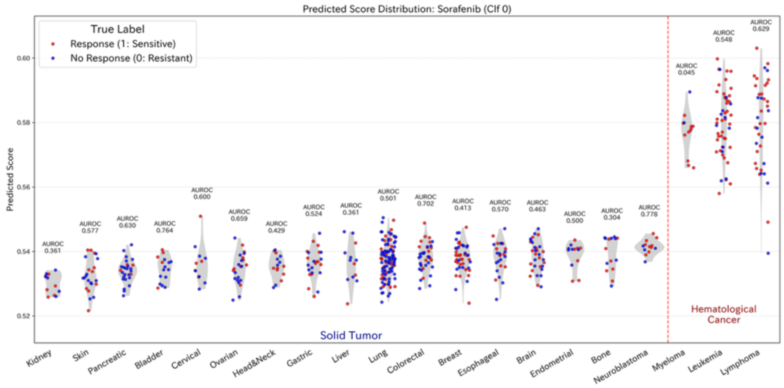
Predicted Score Distribution: Sorafenib (Clf 0)
True Label
Response (1: Sensitive)
No Response (0: Resistant)
Predicted Score
Solid Tumor
Hematological Cancer
Kidney
Skin
Pancreatic
Bladder
Cervical
Ovarian
Head&Neck
Gastric
Liver
Lung
Colorectal
Breast
Esophageal
Brain
Endometrial
Bone
Neuroblastoma
Myeloma
Leukemia
Lymphoma

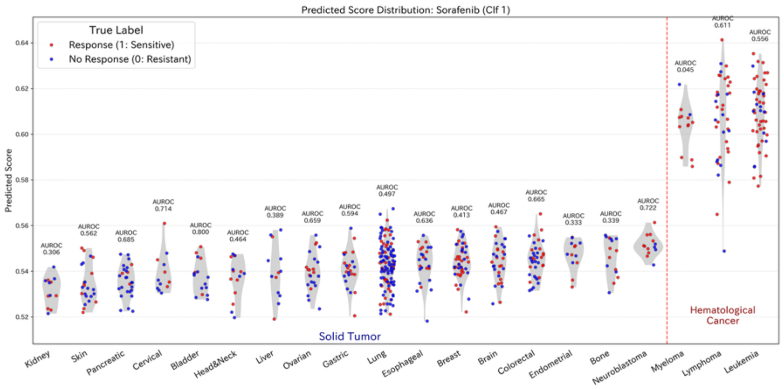
Predicted Score Distribution: Sorafenib (Clf 1)
True Label
Response (1: Sensitive)
No Response (0: Resistant)
Predicted Score
Solid Tumor
Hematological Cancer
Kidney
Skin
Pancreatic
Cervical
Bladder
Head&Neck
Liver
Ovarian
Gastric
Lung
Esophageal
Breast
Brain
Colorectal
Endometrial
Bone
Neuroblastoma
Myeloma
Lymphoma
Leukemia

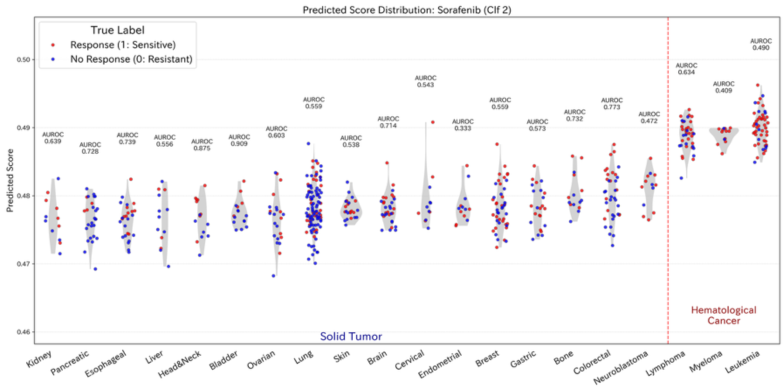
Predicted Score Distribution: Sorafenib (Clf 2)
True Label
Response (1: Sensitive)
No Response (0: Resistant)
Predicted Score
Solid Tumor
Hematological Cancer
Kidney
Pancreatic
Esophageal
Liver
Head&Neck
Bladder
Ovarian
Lung
Skin
Brain
Cervical
Endometrial
Breast
Gastric
Bone
Colorectal
Neuroblastoma
Lymphoma
Myeloma
Leukemia

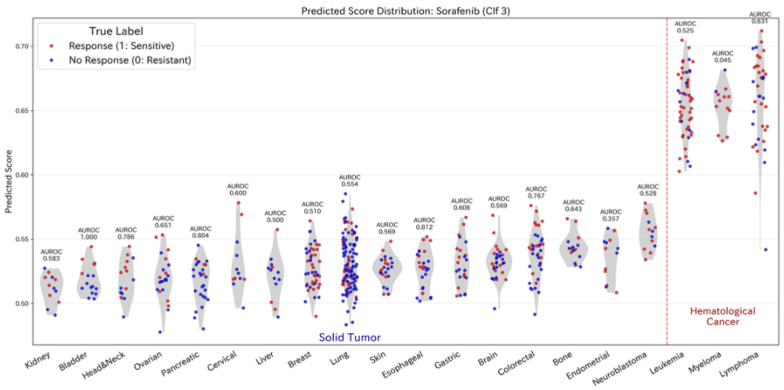
Predicted Score Distribution: Sorafenib (Clf 3)
True Label
Response (1: Sensitive)
No Response (0: Resistant)
Predicted Score
Solid Tumor
Hematological Cancer
Kidney
Bladder
Head&Neck
Ovarian
Pancreatic
Cervical
Liver
Breast
Lung
Skin
Esophageal
Gastric
Brain
Colorectal
Bone
Endometrial
Neuroblastoma
Leukemia
Myeloma
Lymphoma

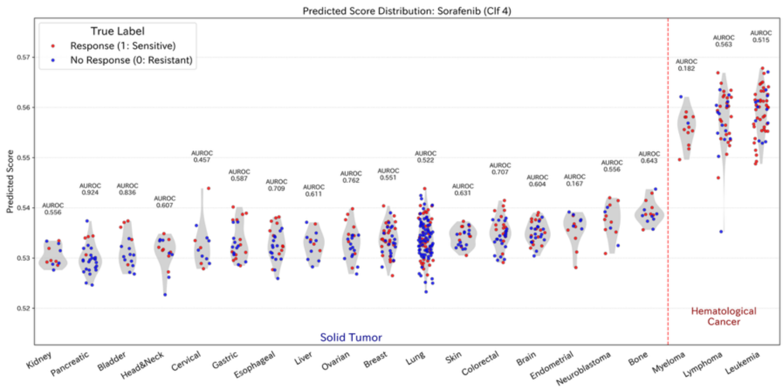
Predicted Score Distribution: Sorafenib (Clf 4)
True Label
Response (1: Sensitive)
No Response (0: Resistant)
Predicted Score
Solid Tumor
Hematological Cancer
Kidney
Pancreatic
Bladder
Head&Neck
Cervical
Gastric
Esophageal
Liver
Ovarian
Breast
Lung
Skin
Colorectal
Brain
Endometrial
Neuroblastoma
Bone
Myeloma
Lymphoma
Leukemia

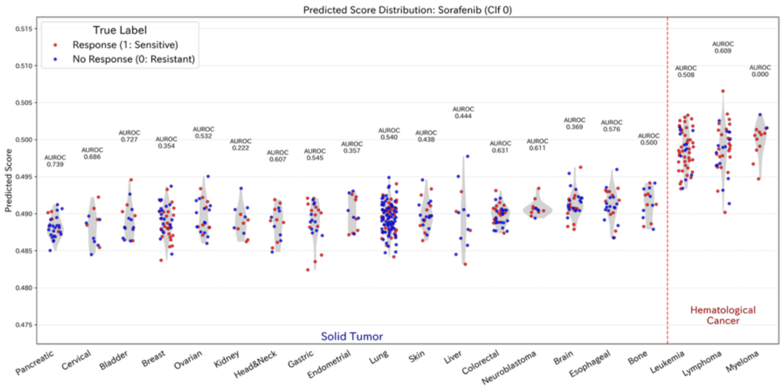
Predicted Score Distribution: Sorafenib (Clf 0)
True Label
Response (1: Sensitive)
No Response (0: Resistant)
Predicted Score
Solid Tumor
Hematological Cancer
Pancreatic
Cervical
Bladder
Breast
Ovarian
Kidney
Head&Neck
Gastric
Endometrial
Lung
Skin
Liver
Colorectal
Neuroblastoma
Brain
Esophageal
Bone
Leukemia
Lymphoma
Myeloma

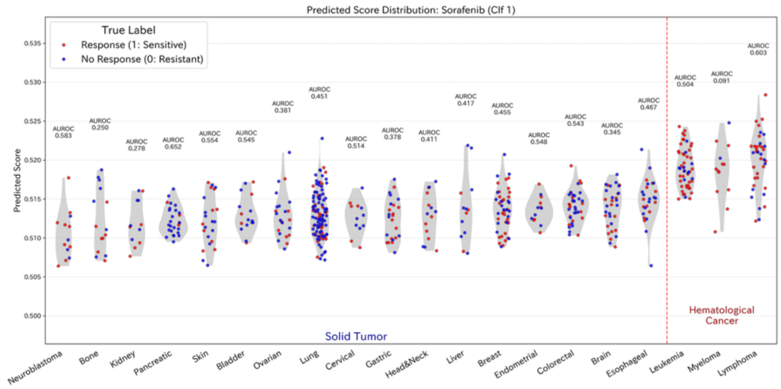
Predicted Score Distribution: Sorafenib (Clf 1)
True Label
Response (1: Sensitive)
No Response (0: Resistant)
Predicted Score
Solid Tumor
Hematological Cancer
Neuroblastoma
Bone
Kidney
Pancreatic
Skin
Bladder
Ovarian
Lung
Cervical
Gastric
Head&Neck
Liver
Breast
Endometrial
Colorectal
Brain
Esophageal
Leukemia
Myeloma
Lymphoma

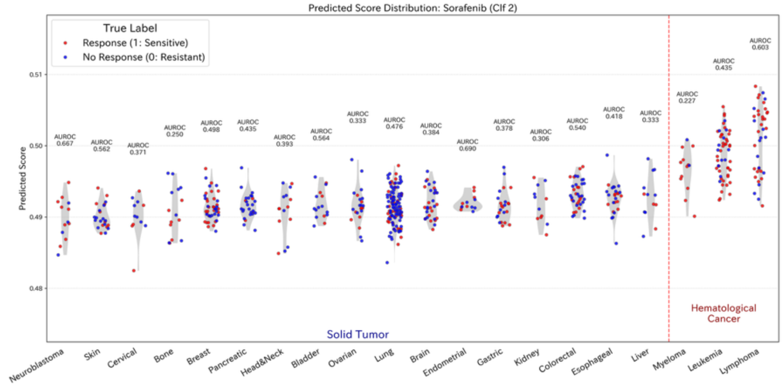
Predicted Score Distribution: Sorafenib (Clf 2)
True Label
Response (1: Sensitive)
No Response (0: Resistant)
Predicted Score
Solid Tumor
Hematological Cancer
Neuroblastoma
Skin
Cervical
Bone
Breast
Pancreatic
Head&Neck
Bladder
Ovarian
Lung
Brain
Endometrial
Gastric
Kidney
Colorectal
Esophageal
Liver
Myeloma
Leukemia
Lymphoma

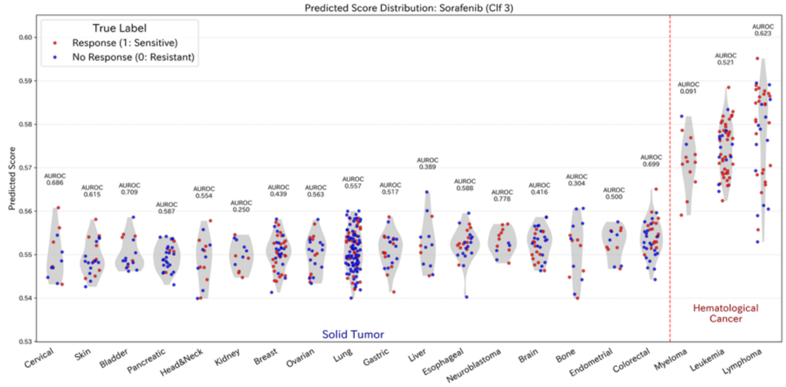
Predicted Score Distribution: Sorafenib (Clf 3)
True Label
Response (1: Sensitive)
No Response (0: Resistant)
Predicted Score
Solid Tumor
Hematological Cancer
Cervical
Skin
Bladder
Pancreatic
Head&Neck
Kidney
Breast
Ovarian
Lung
Gastric
Liver
Esophageal
Neuroblastoma
Brain
Bone
Endometrial
Colorectal
Myeloma
Leukemia
Lymphoma

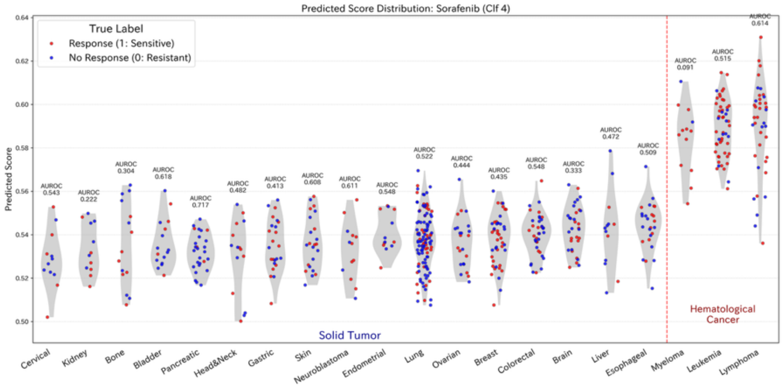
Predicted Score Distribution: Sorafenib (Clf 4)
True Label
Response (1: Sensitive)
No Response (0: Resistant)
Predicted Score
Solid Tumor
Hematological Cancer
Cervical
Kidney
Bone
Bladder
Pancreatic
Head&Neck
Gastric
Skin
Neuroblastoma
Endometrial
Lung
Ovarian
Breast
Colorectal
Brain
Liver
Esophageal
Myeloma
Leukemia
Lymphoma

[Temozolomide]

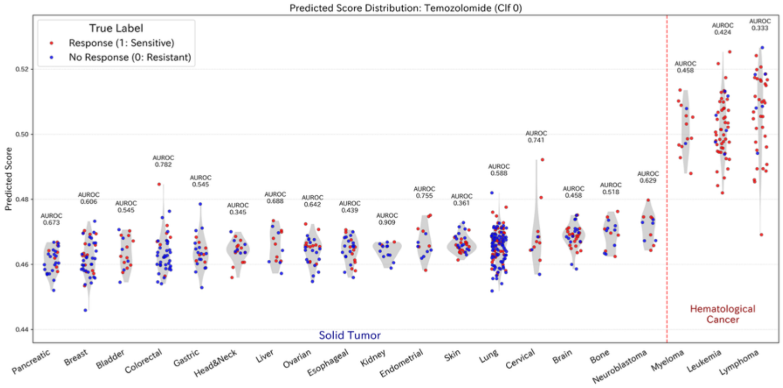


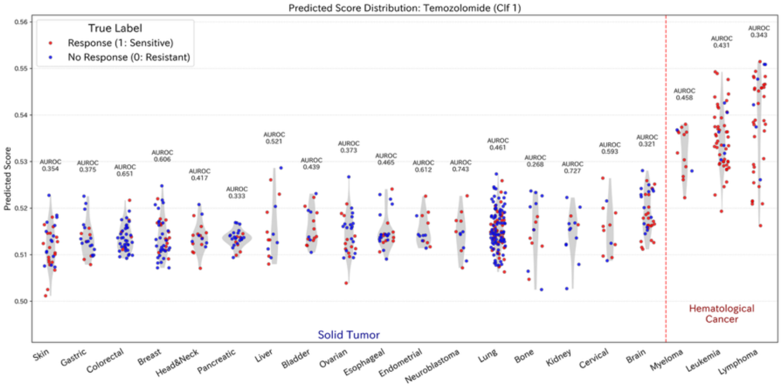


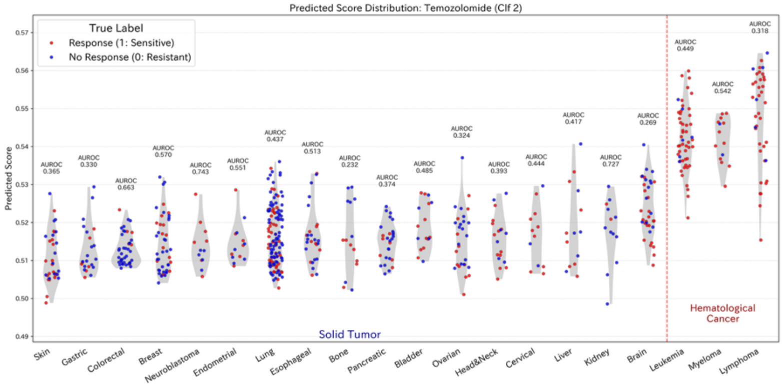


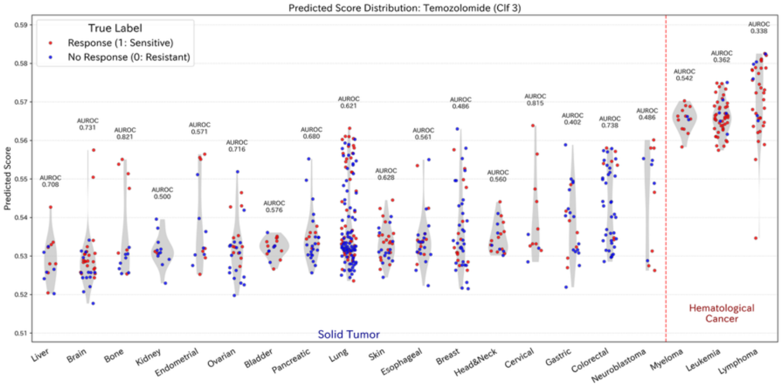


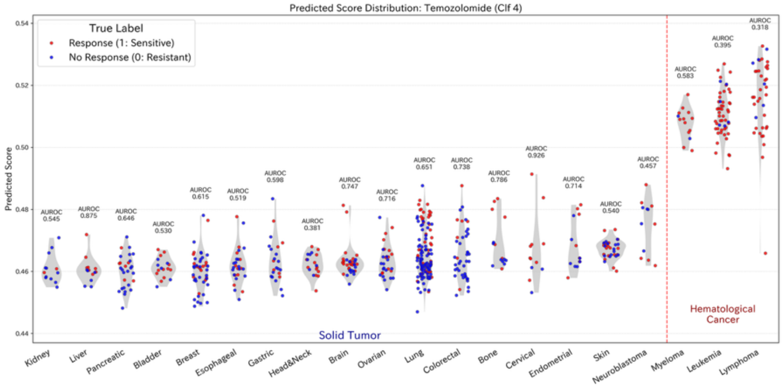


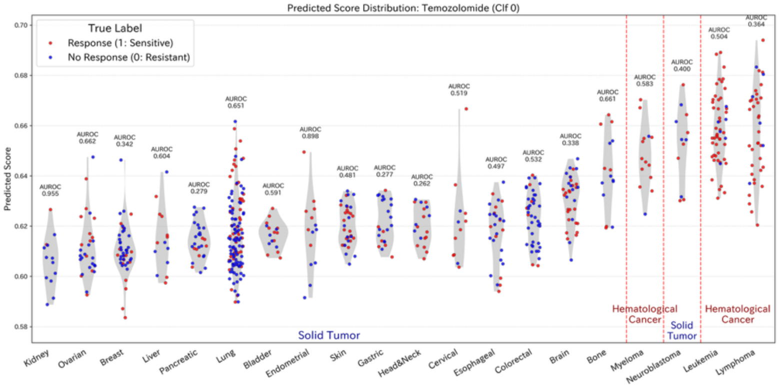


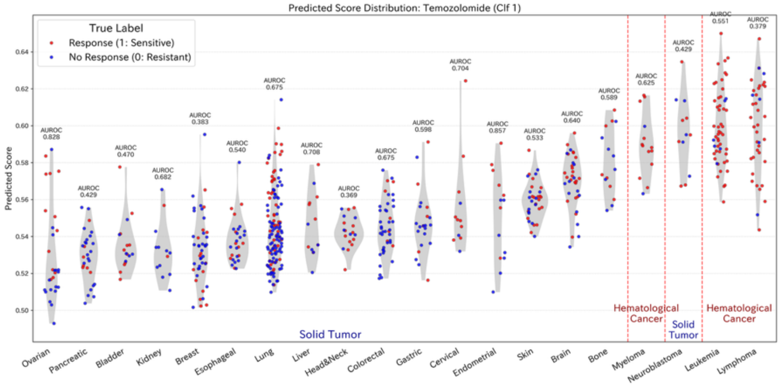


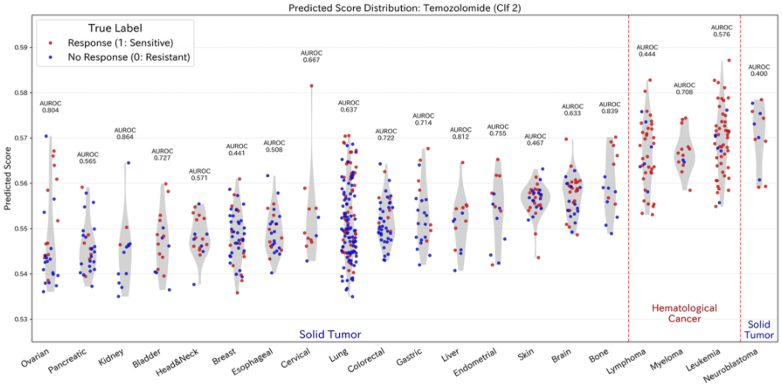

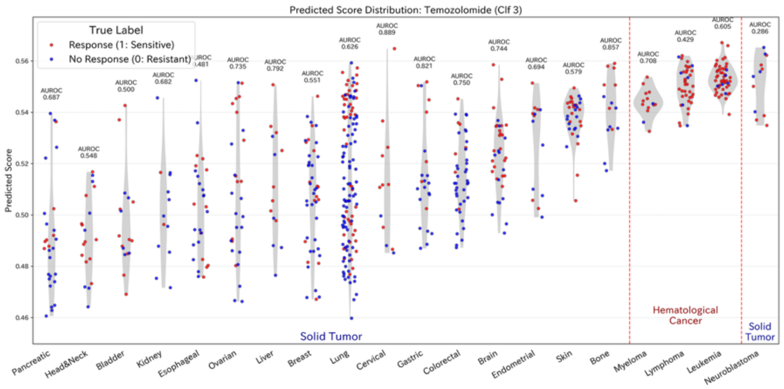
Predicted Score Distribution: Temozolomide (Clf 3)
True Label
Response (1: Sensitive)
No Response (0: Resistant)
Predicted Score
Solid Tumor
Hematological Cancer
Solid Tumor
Pancreatic
Head&Neck
Bladder
Kidney
Esophageal
Ovarian
Liver
Breast
Lung
Cervical
Gastric
Colorectal
Brain
Endometrial
Skin
Bone
Myeloma
Lymphoma
Leukemia
Neuroblastoma

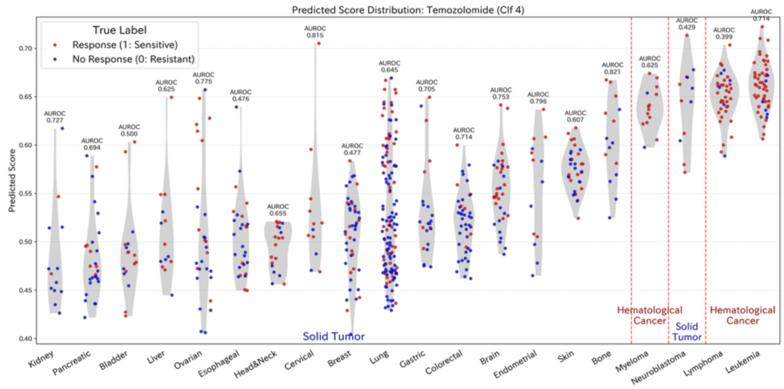
Predicted Score Distribution: Temozolomide (Clf 4)
True Label
Response (1: Sensitive)
No Response (0: Resistant)
Predicted Score
Solid Tumor
Hematological Cancer
Solid Tumor
Hematological Cancer
Kidney
Pancreatic
Bladder
Liver
Ovarian
Esophageal
Head&Neck
Cervical
Breast
Lung
Gastric
Colorectal
Brain
Endometrial
Skin
Bone
Myeloma
Neuroblastoma
Lymphoma
Leukemia

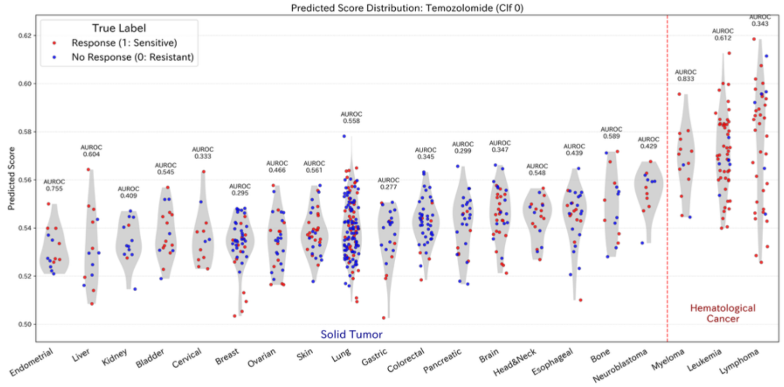
Predicted Score Distribution: Temozolomide (Clf 0)
True Label
Response (1: Sensitive)
No Response (0: Resistant)
Predicted Score
Solid Tumor
Hematological Cancer
Endometrial
Liver
Kidney
Bladder
Cervical
Breast
Ovarian
Skin
Lung
Gastric
Colorectal
Pancreatic
Brain
Head&Neck
Esophageal
Bone
Neuroblastoma
Myeloma
Leukemia
Lymphoma

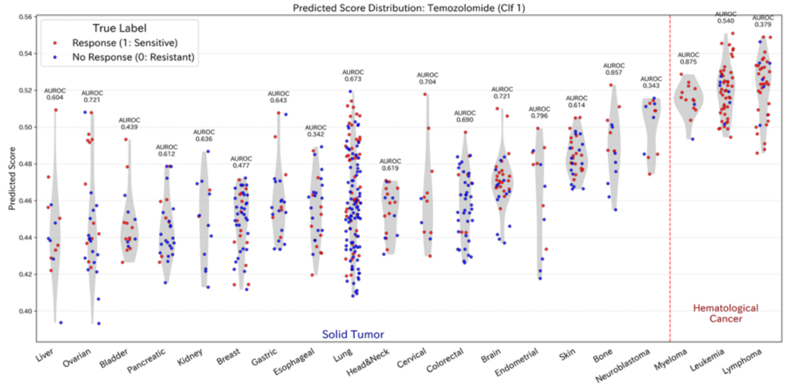
Predicted Score Distribution: Temozolomide (Clf 1)
True Label
Response (1: Sensitive)
No Response (0: Resistant)
Predicted Score
Solid Tumor
Hematological Cancer
Liver
Ovarian
Bladder
Pancreatic
Kidney
Breast
Gastric
Esophageal
Lung
Head&Neck
Cervical
Colorectal
Brain
Endometrial
Skin
Bone
Neuroblastoma
Myeloma
Leukemia
Lymphoma

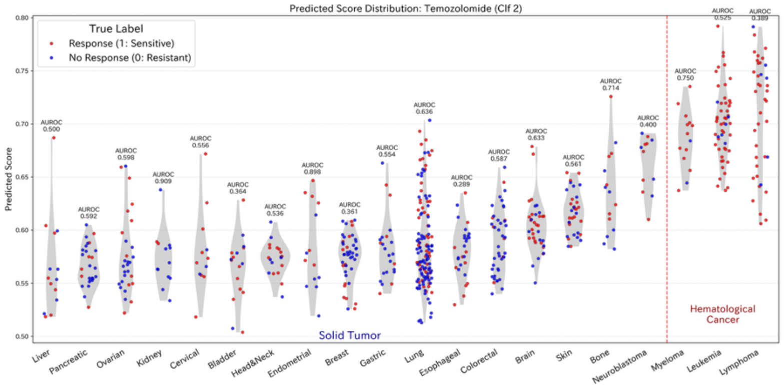
Predicted Score Distribution: Temozolomide (Clf 2)
True Label
Response (1: Sensitive)
No Response (0: Resistant)
Predicted Score
Solid Tumor
Hematological Cancer
Liver
Pancreatic
Ovarian
Kidney
Cervical
Bladder
Head&Neck
Endometrial
Breast
Gastric
Lung
Esophageal
Colorectal
Brain
Skin
Bone
Neuroblastoma
Myeloma
Leukemia
Lymphoma

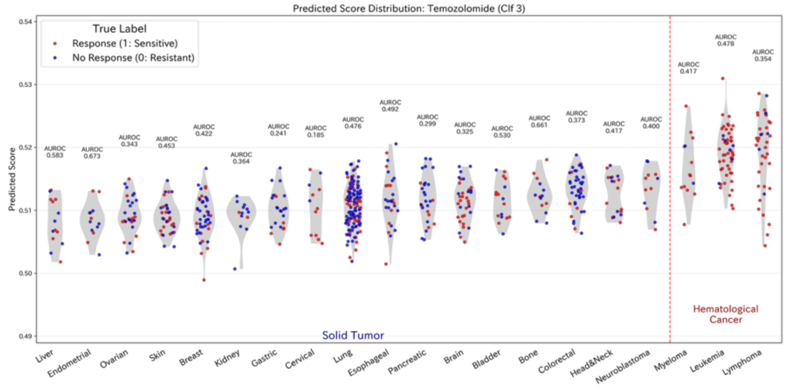
Predicted Score Distribution: Temozolomide (Clf 3)
True Label
Response (1: Sensitive)
No Response (0: Resistant)
Predicted Score
Solid Tumor
Hematological Cancer
Liver
Endometrial
Ovarian
Skin
Breast
Kidney
Gastric
Cervical
Lung
Esophageal
Pancreatic
Brain
Bladder
Bone
Colorectal
Head&Neck
Neuroblastoma
Myeloma
Leukemia
Lymphoma

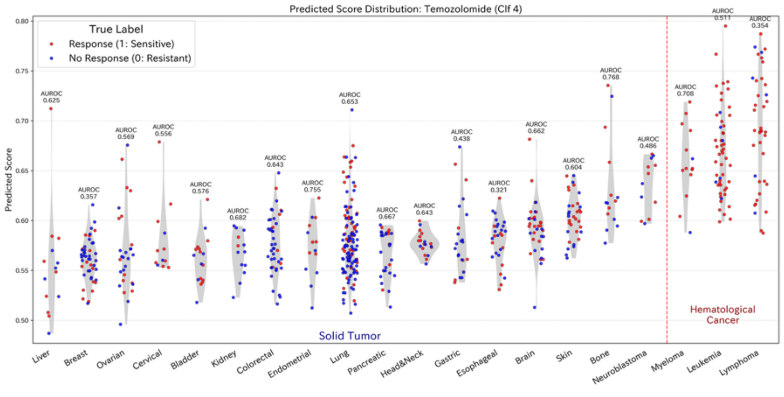
Predicted Score Distribution: Temozolomide (Clf 4)
True Label
Response (1: Sensitive)
No Response (0: Resistant)
Predicted Score
Solid Tumor
Hematological Cancer
Liver
Breast
Ovarian
Cervical
Bladder
Kidney
Colorectal
Endometrial
Lung
Pancreatic
Head&Neck
Gastric
Esophageal
Brain
Skin
Bone
Neuroblastoma
Myeloma
Leukemia
Lymphoma

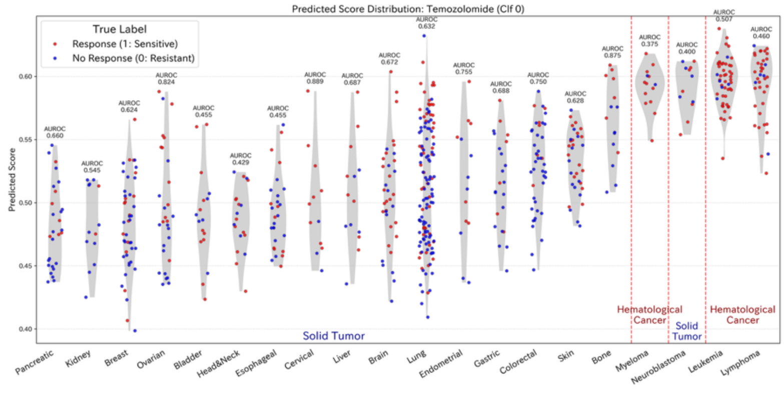
Predicted Score Distribution: Temozolomide (Clf 0)
True Label
Response (1: Sensitive)
No Response (0: Resistant)
Predicted Score
Solid Tumor
Hematological Cancer
Solid Tumor
Hematological Cancer
Pancreatic
Kidney
Breast
Ovarian
Bladder
Head&Neck
Esophageal
Cervical
Liver
Brain
Lung
Endometrial
Gastric
Colorectal
Skin
Bone
Myeloma
Neuroblastoma
Leukemia
Lymphoma

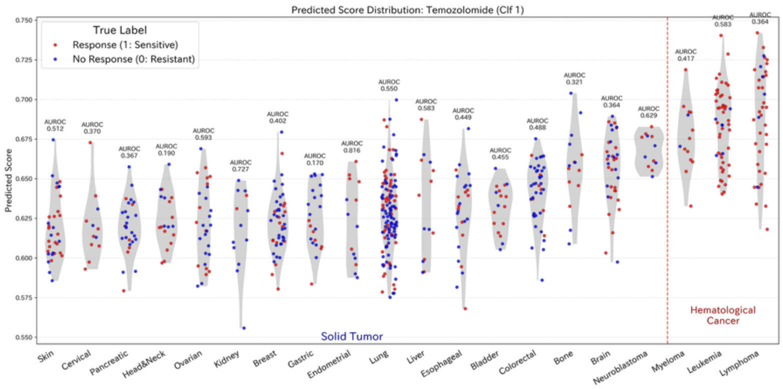
Predicted Score Distribution: Temozolomide (Clf 1)
True Label
Response (1: Sensitive)
No Response (0: Resistant)
Predicted Score
Solid Tumor
Hematological Cancer
Skin
Cervical
Pancreatic
Head&Neck
Ovarian
Kidney
Breast
Gastric
Endometrial
Lung
Liver
Esophageal
Bladder
Colorectal
Bone
Brain
Neuroblastoma
Myeloma
Leukemia
Lymphoma

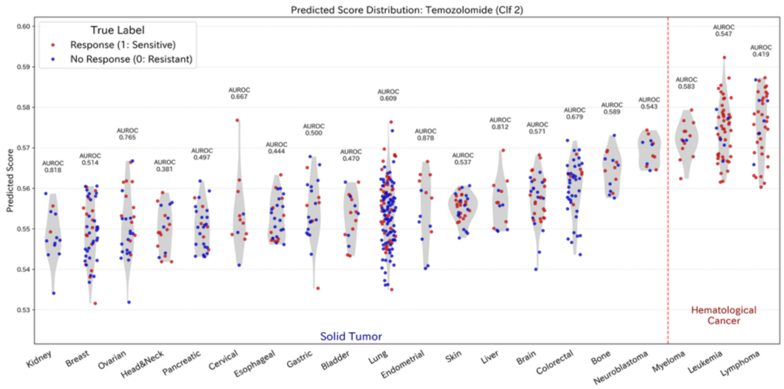
Predicted Score Distribution: Temozolomide (Clf 2)
True Label
Response (1: Sensitive)
No Response (0: Resistant)
Predicted Score
Solid Tumor
Hematological Cancer
Kidney
Breast
Ovarian
Head&Neck
Pancreatic
Cervical
Esophageal
Gastric
Bladder
Lung
Endometrial
Skin
Liver
Brain
Colorectal
Bone
Neuroblastoma
Myeloma
Leukemia
Lymphoma

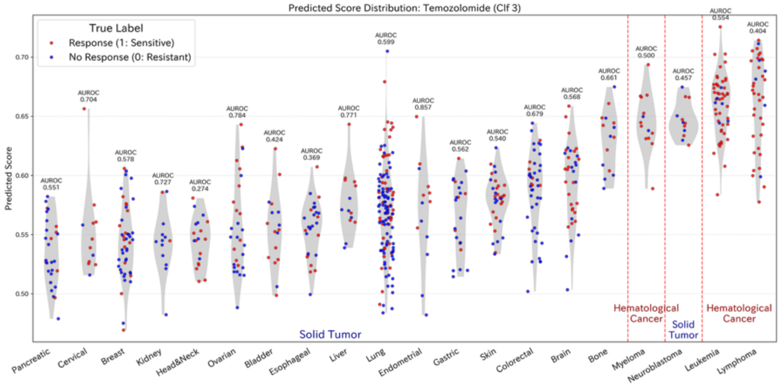
Predicted Score Distribution: Temozolomide (Clf 3)
True Label
Response (1: Sensitive)
No Response (0: Resistant)
Predicted Score
Solid Tumor
Hematological Cancer
Solid Tumor
Hematological Cancer
Pancreatic
Cervical
Breast
Kidney
Head&Neck
Ovarian
Bladder
Esophageal
Liver
Lung
Endometrial
Gastric
Skin
Colorectal
Brain
Bone
Myeloma
Neuroblastoma
Leukemia
Lymphoma

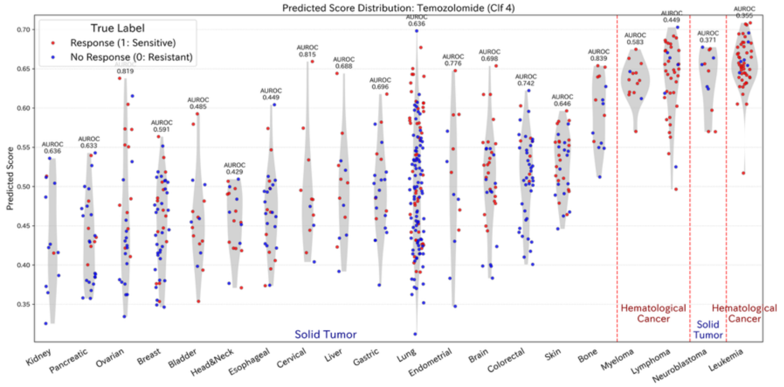
Predicted Score Distribution: Temozolomide (Clf 4)
True Label
Response (1: Sensitive)
No Response (0: Resistant)
Predicted Score
Solid Tumor
Hematological Cancer
Solid Tumor
Hematological Cancer
Kidney
Pancreatic
Ovarian
Breast
Bladder
Head&Neck
Esophageal
Cervical
Liver
Gastric
Lung
Endometrial
Brain
Colorectal
Skin
Bone
Myeloma
Lymphoma
Neuroblastoma
Leukemia

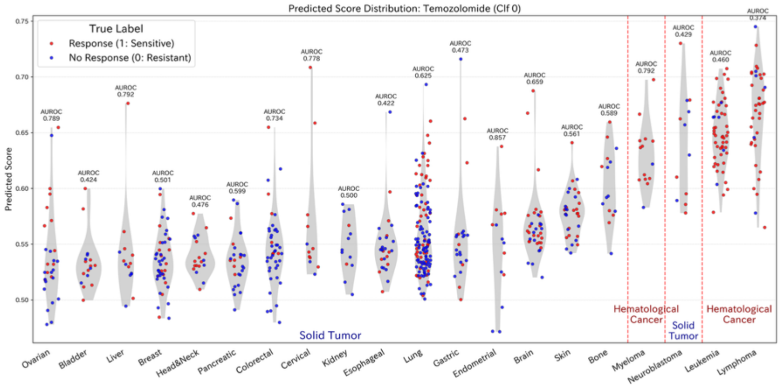
Predicted Score Distribution: Temozolomide (Clf 0)
True Label
Response (1: Sensitive)
No Response (0: Resistant)
Predicted Score
Solid Tumor
Hematological Cancer
Solid Tumor
Hematological Cancer
Ovarian
Bladder
Liver
Breast
Head&Neck
Pancreatic
Colorectal
Cervical
Kidney
Esophageal
Lung
Gastric
Endometrial
Brain
Skin
Bone
Myeloma
Neuroblastoma
Leukemia
Lymphoma

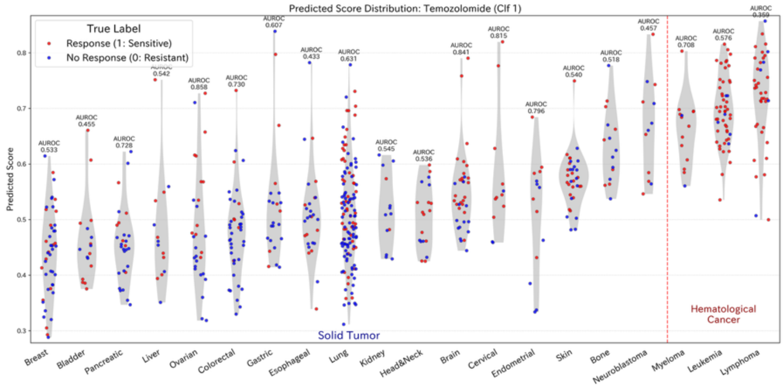
Predicted Score Distribution: Temozolomide (Clf 1)
True Label
Response (1: Sensitive)
No Response (0: Resistant)
Predicted Score
Solid Tumor
Hematological Cancer
Breast
Bladder
Pancreatic
Liver
Ovarian
Colorectal
Gastric
Esophageal
Lung
Kidney
Head&Neck
Brain
Cervical
Endometrial
Skin
Bone
Neuroblastoma
Myeloma
Leukemia
Lymphoma

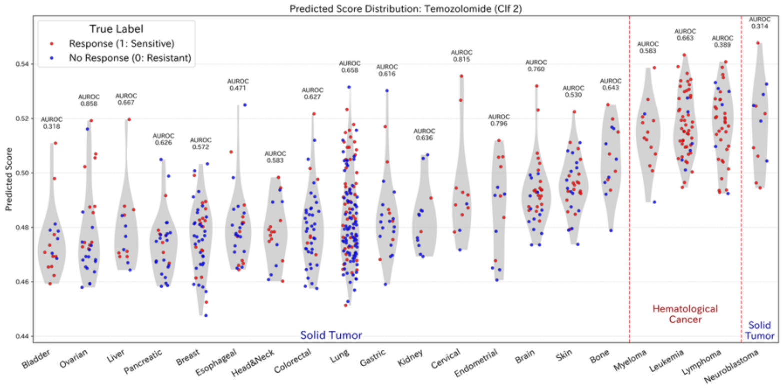
Predicted Score Distribution: Temozolomide (Clf 2)
True Label
Response (1: Sensitive)
No Response (0: Resistant)
Predicted Score
Solid Tumor
Hematological Cancer
Solid Tumor
Bladder
Ovarian
Liver
Pancreatic
Breast
Esophageal
Head&Neck
Colorectal
Lung
Gastric
Kidney
Cervical
Endometrial
Brain
Skin
Bone
Myeloma
Leukemia
Lymphoma
Neuroblastoma

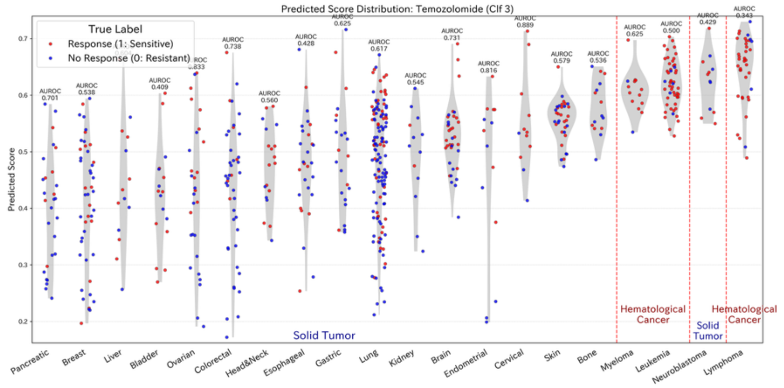
Predicted Score Distribution: Temozolomide (Clf 3)
True Label
Response (1: Sensitive)
No Response (0: Resistant)
Predicted Score
Solid Tumor
Hematological Cancer
Solid Tumor
Hematological Cancer
Pancreatic
Breast
Liver
Bladder
Ovarian
Colorectal
Head&Neck
Esophageal
Gastric
Lung
Kidney
Brain
Endometrial
Cervical
Skin
Bone
Myeloma
Leukemia
Neuroblastoma
Lymphoma

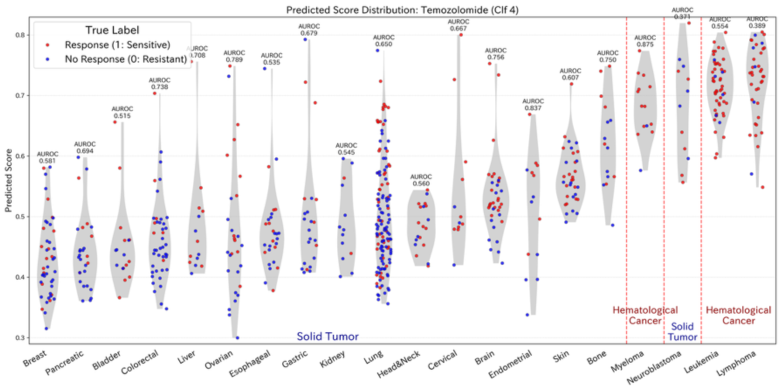


**Figure S1. Prediction score distributions across cancer types for all five drugs. Comprehensive results for all 25 predictors (5 seeds × 5 folds) are shown for 5-Fluorouracil, Cisplatin, Gemcitabine, Sorafenib, and Temozolomide.**
